# Regulatory landscape of widespread stop codon readthrough in *Drosophila*

**DOI:** 10.64898/2026.06.10.731275

**Authors:** Nadezhda Makarova, Daniel Martín de la Fuente, Mihajlo Stašuk, Ivet Ortiga Martínez, Jonas Pöhls, Irwin Jungreis, Manolis Kellis, Federica Battistini, Diego Garrido Martín, Alain Lescure, Agnes Toth-Petroczy, Marco Mariotti

## Abstract

In stop codon readthrough (SCR), ribosomes continue to translate past the stop. While programmed SCR is rare in mammals, it appears remarkably widespread in insects, yet its regulation and biological significance remain poorly understood. Here, we systematically characterize SCR in *Drosophila melanogaster* by integrating ribosome profiling, comparative genomics, proteomics, and functional screening. We identify ∼1,400 SCR genes, reveal sequence features associated with programmed SCR, and distinguish constitutive and tissue-restricted modes. We identify conserved *cis*-elements sufficient to stimulate constitutive SCR and demonstrate that many remain functional in mammalian cells. Unexpectedly, the major distinction between insect and mammalian cells was not termination efficiency, but insect responsiveness to SCR-promoting regulatory elements. SCR was further associated with increased overall protein output, suggesting a function beyond proteome diversification. Our results establish SCR as a pervasive regulatory mechanism in *Drosophila* and provide a new framework for its study through the Drosophila Stop Codon Readthrough Atlas.

## Introduction

Stop codon readthrough (SCR) is a recoding event in which a ribosome inserts an amino acid at a stop codon, allowing translation to continue in-frame until a downstream stop codon is encountered ^1^. Under normal conditions, termination is mediated by the eRF1-eRF3 complex in eukaryotes, which recognizes stop codons in the ribosomal A site and catalyzes peptide release ^2,3^. Yet, near-cognate tRNAs may instead be accommodated and promote elongation, resulting in SCR ^4–6^. This process primarily depends on the local stop codon context, though its determinants remain incompletely understood. In animals including *Drosophila melanogaster* and many other organisms, a specialized form of SCR enables recoding of UGA to selenocysteine (Sec) ^7^. This process is mechanistically well defined and relies on a dedicated cognate tRNA^Sec^ and a distal *cis*-element known as the SECIS ^8^. In contrast, other forms lack clear signatures and remain less systematically documented. In this work, we focus on non-Sec SCR.

SCR can arise either as a by-product of imperfect translation termination (non-programmed SCR), or as an evolutionarily conserved recoding mechanism (programmed SCR). Non-programmed SCR reflects occasional and typically rare decoding errors. The resulting extensions can destabilize proteins and trigger mRNA surveillance pathways ^9–11^. In contrast, programmed SCR leads to the consistent production of C-terminally extended isoforms for a substantial fraction of translation events ^12–14^, whose functional relevance has been shown for some *Drosophila* genes ^15–17^. In programmed SCR, the protein-coding extension is typically evolutionarily conserved and supported by *cis*-acting features, such as optimized nucleotide context and stop-proximal RNA structures.

Two main complementary approaches have been used to detect SCR: comparative genomics and ribosome profiling (Ribo-Seq). The first approach seeks evidence of programmed SCR by identifying protein-level sequence conservation downstream of stop codons across related species ^18–20^. Ribo-Seq, in contrast, offers direct experimental SCR evidence by sequencing ribosome-protected mRNA fragments at nucleotide resolution, thereby revealing active translation within 3′UTRs on a transcriptome-wide scale ^21,22^. Notably, both approaches revealed striking taxonomic differences among metazoa. While evolutionary approaches yielded only a handful of programmed SCR candidates in mammals, hundreds of genes were predicted in insects: a study estimated that the occurrence of programmed SCR in *Drosophila* approached ∼900 genes, and specifically pinpointed 333 well-supported cases ^23^. Consistently, Ribo-Seq studies reported substantially more cases in *Drosophila* than in yeast and human ^24^. Comparative sequence analyses further identified hundreds of SCR genes in *Anopheles* and *Bombus*, and suggested that extensive programmed SCR emerged after the divergence from centipedes, implying that this phenomenon is widespread across insects ^23,25^.

Despite the prevalence of SCR in *Drosophila*, only a limited number of genes have been investigated *in vivo*: *kelch* (*kel*), *traffic jam* (*tj*), *ventral veins lacking* (*vvl/dfr*), *headcase* (*heca/hdc*), and *synapsin* (*syn*)^15,16,26–29^. In many cases, SCR was studied by altering stop codons, SCR extensions, or proposed recoding signals, revealing a broad spectrum of phenotypic outcomes depending on the gene examined. Importantly, several genes display enhanced readthrough in neuronal tissues ^15–17,28,30,31^. The underlying mechanism is yet unclear, and is thought to involve tissue-specific *trans*-factors. In *Drosophila*, the only identified factor so far is the brain-enriched eRF1 isoform eRF1H. This differs from the canonical protein in the C-terminal region required for eRF3 interaction, and was shown to promote SCR of a reporter carrying the Tobacco mosaic virus (TMV) readthrough motif when expressed in S2 cells ^16^. In addition, RNAi screens identified *Upf1*, a component of the nonsense-mediated decay (NMD) pathway, as regulator of SCR, though its role does not appear to be restricted to neurons ^30^.

Collectively, previous studies established SCR as a prominent feature of insect gene expression, yet its prevalence across biological contexts, molecular determinants, amino acid incorporation patterns, and functional impact remain largely unresolved. Given the hundreds of genes predicted to undergo SCR in insects, addressing these questions arguably requires a comprehensive top-down approach capable not only of revealing common regulatory principles, but also resolving the underlying architecture of SCR into distinct regulatory modes.

Here, we present a systematic multi-omics characterization of SCR in *D. melanogaster* that integrates large-scale ribosome profiling analyses with comparative genomics, proteomics, and functional reporter assays. This enabled us to generate the largest catalog of *D. melanogaster* SCR genes to date, stratified into confidence tiers according to features associated with programmed readthrough. We uncover sequence and structural signatures linked to SCR, reveal distinct modes of SCR activation across biological conditions, and identify conserved *cis*-regulatory elements sufficient to stimulate readthrough in insect cells, most of which are portable to mammalian cells. We also show that *cis*-stimulated SCR occurs at much higher rates in *Drosophila* cells than in mammalian cells despite comparable baseline termination efficiency, indicating enhanced regulatory responsiveness. Furthermore, our analyses reveal amino acid incorporation biases at stop codons and suggest that SCR can increase the translational output of the mRNA beyond the production of SCR-derived isoforms. Together, our work sheds light into the regulatory landscape of *Drosophila* SCR and provides a dedicated new online resource for its exploration, the *Drosophila* Stop Codon Readthrough Atlas.

## Results

### Identification of novel SCR genes by ribosome profiling

To systematically identify genes showing evidence of SCR at a genome-wide scale, we gathered a comprehensive collection of publicly available ribosome profiling datasets for *D. melanogaster*. Our analysis included 79 samples from 9 datasets (Supplementary Table T1), spanning a broad range of developmental stages and tissues, including adult heads and bodies, embryos at multiple stages, larvae, and cultured S2 cells. Notably, these datasets were generated using heterogeneous experimental protocols, with differences in nuclease digestion and rRNA depletion strategies. To interrogate Ribo-Seq data, we generated an extended genomic annotation consisting of three regions per protein-coding gene (Figure 1A): (i) Canonical CDS, as annotated at Flybase without in-frame stops; (ii) SCR extension, i.e. the region between the annotated stop codon and the next in-frame stop; (iii) Negative control region, i.e. the region downstream of the second stop codon. A single transcript per gene was considered, selected through criteria to minimize false-positive SCR detection (Methods). To improve detection power, we implemented an aggregation strategy to average Ribo-Seq density at each nucleotide position across all samples, followed by a noise-removal procedure designed to eliminate position-specific artefactual coverage that may yield false positives (Supplementary Figure S1, Methods).

**Figure 1.**
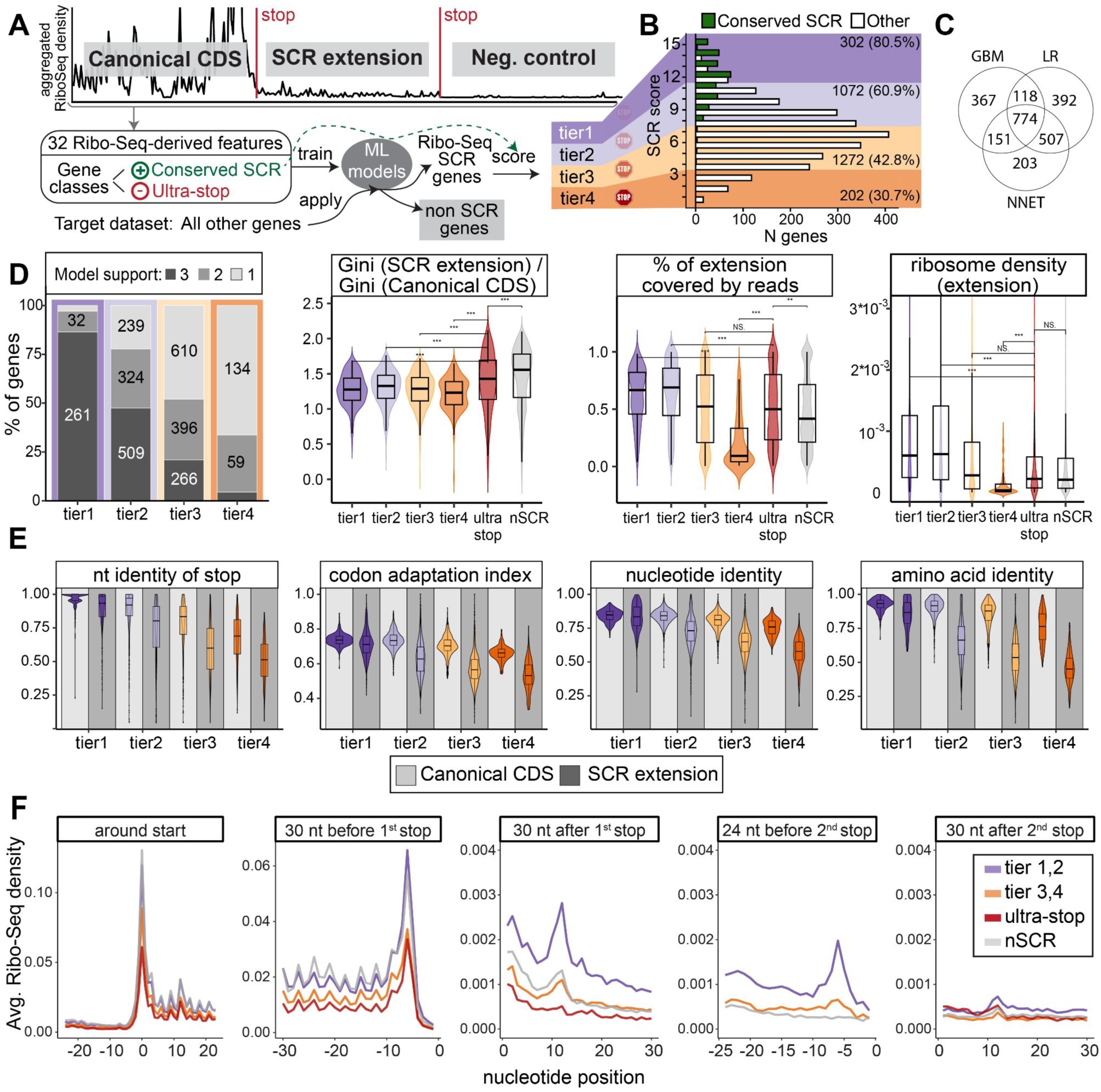
Identification of stop codon readthrough (SCR) genes using ribosome profiling. **(A)** Workflow for identification of SCR genes. The scheme at the top illustrates a representative known SCR gene (*gish* ^20^), highlighting the relevant gene regions. Aggregated, noise-filtered ribosome Ribo-Seq density data, stratified by region, are used as input for machine-learning classifiers. These classifiers identify putative SCR genes based on the similarity of their Ribo-Seq profiles to those of SCR genes previously detected by conservation analyses, used here for training. Candidate genes are subsequently scored using an heuristic procedure and assigned to confidence tiers. **(B)** Overview of putative SCR gene set. Bar color differentiates the SCR genes previously identified by conservation (green) ^20,23^ from our new Ribo-Seq-based predictions (white). Background colors correspond to tiers. Numbers on the right denote gene counts per tier, with estimated precision based solely on classifier performance shown in parentheses (Supplementary Tables T4,T5) **(C)** Overlap of SCR gene predictions across machine-learning models. Only unlabelled predictions (white colored in histogram B) are displayed here. The Venn diagram shows the overlap of SCR predictions obtained using Gradient Boosting Machine (GBM), Logistic Regression (LR), and Neural Network (NNET) classifiers. Overlap of SCR gene predictions for positive and negative training sets are shown in Supplementary Figure S4. **(D)** Examples of features used for Ribo-Seq-based SCR identification (pre-scoring). These include: (i) proportion of genes supported by 1, 2, or 3 classifier models. Absolute gene numbers are shown for categories comprising >10% of genes. (ii) ratio between the Gini coefficients of Ribo-Seq densities for the SCR extension and the canonical CDS. Higher Gini values indicate increased heterogeneity in coverage, suggesting that the signal is less indicative of active translation and more likely represents stochastic or artifactual reads. (iii) Percentage of SCR extension nucleotides covered by Ribo-Seq reads. (iv) Ribosome density in the SCR extension. Statistical comparisons are indicated (NS, not significant; **, p < 0.01; ***, p < 0.001, two-sided Wilcoxon rank-sum test). (E) Distribution across SCR gene tiers of metrics used for scoring. Metrics were calculated separately for the canonical CDS and SCR extension (background color). Refer to Methods for the full list of scoring metrics. “Identity”measures indicate stop, nucleotide, and protein conservation in *Drosophila* whole-genome alignments (WGAs). Note that nucleotide and protein conservation are displayed also for the canonical CDS, yet only the SCR extension was included in scoring. The codon adaptation index expresses the codon usage bias and ranges from 0 (not optimized) to 1 (highly optimized). **(F)** Aggregated Ribo-Seq densities along genomic positions. Genes were aligned to the annotated start, annotated stop codon, or next in-frame stop (facets), and read densities were averaged across gene groups (legend on the right) at each nucleotide position. Ultrastop genes were aligned identically, using their native second stop codon rather than the pseudoextensions used for machine learning. Boxplots show median (center line), interquartile range (IQR; box), and 1.5×IQR whiskers.

Earlier studies relied on manually crafted filters on Ribo-Seq-derived metrics to identify SCR candidates ^22,24,32,33^. Because such approaches depend on arbitrary thresholds, we instead used machine learning to recognize the characteristic signatures of SCR from aggregated Ribo-Seq data. We trained our framework with the profiles of “conserved SCR” genes identified by evolutionary conservation across *Drosophila* (Figure 1A; Methods). As negative set, we used “ultrastop” genes, which terminate with two in-frame stop codons separated by at most one sense codon. We trained three classifier models which achieved 75-88% accuracy (Supplementary Table T2), although this likely underestimates their true performance, since some conserved SCR genes may be active only in biological contexts absent from available Ribo-Seq samples (Supplementary Figure S2). Classification emerged from the combined contribution of multiple Ribo-Seq features rather than any single dominant metric (Supplementary Figure S3). At this stage, we intentionally excluded conservation and sequence features from model training, to capture experimental evidence of both programmed and non-programmed SCR from Ribo-Seq metrics.

We performed SCR discovery by applying our models to 6,158 unlabelled genes, i.e. all protein coding genes with sufficient coverage and adequate 3’UTR length that were not included in training (Methods). Collectively, the three classifier models yielded 2,512 candidate SCR genes, including 774 genes predicted by all models (Figure 1C). Because somatic mutations or RNA editing at the stop codon may result in apparent SCR, we analyzed raw sequencing reads to identify such cases, which ultimately led to discarding eight genes from downstream analyses (Methods). 336 out of 366 genes from the conserved SCR set were recovered by at least one model. Together, these sets were combined to generate a comprehensive list of 2,848 putative SCR genes (Supplementary Table T3). Because our primary goal at this stage was sensitive detection of SCR, we retained genes supported even by a single classifier despite the expected increase in false positives. We next refined this broad candidate set by incorporating additional evidence associated with programmed SCR as described below, thereby defining high-confidence subsets.

#### Composite SCR scoring reveals high-confidence programmed SCR genes

To define high-confidence SCR gene sets, we developed a composite SCR score combining Ribo-Seq evidence with evolutionary and sequence-related characteristics associated with programmed SCR. These 15 features were not used during model training, and included: (i–iii) Ribo-Seq classifier output; (iv) SCR extension length; (v–vi) G/C-ending codon frequency in CDS and extension; (vii–viii) codon usage bias in CDS and extension; (ix–x) stop codon conservation; (xi) stop-spanning reads; (xii) in-frame stop-spanning reads; (xiii–xiv) extension conservation at nucleotide and protein level; and (xv) Ribo-Seq enrichment at the second stop codon (Methods, Figure 1E, Supplementary Figure S5). The composite score was the number of classifiers that predicted SCR plus the number of features (iv-xv) that exceeded a feature-specific threshold, namely the 25th percentile among conserved SCR genes predicted by all three classifiers. For downstream analyses, genes were manually assigned into four tiers based on their composite SCR score (Figure 1A). As shown hereafter, tiers 1-2 represent a high-confidence group of SCR genes highly enriched for programmed SCR, and were later used for discovery of SCR-related features. By contrast, tiers 3-4 progressively accumulate non-programmed cases and false positives, effectively serving as negative sets (Supplementary Tables T4,T5).

Multiple lines of evidence supported the interpretation of the SCR score as an indicator of genuine programmed SCR. First, previously described conserved SCR genes were strongly enriched in tiers 1 and 2 (Figure 1B). Second, tier 1-2 genes displayed clearer and more homogeneous Ribo-Seq density across the SCR extension, together with stronger agreement between the three prediction models, than tiers 3-4 (Figure 1D, Supplementary Figure S6,S7 Supplementary table T5). Third, tier 1-2 genes showed substantially higher nucleotide and amino acid conservation within the SCR extension region (Figure 1E). Crucially, these tier-dependent trends were preserved when all genes used for training were excluded from the analysis, supporting the integrity of the pipeline (Supplementary Figure S6,S7,S8).

Analyses of aggregated Ribo-Seq data further corroborated the quality of our SCR gene set (Figure 1F). Tier 1-2 genes exhibited elevated ribosome density downstream of the stop codon compared to all other gene groups, while their canonical CDS expression was similar to non-SCR genes (i.e. genes not predicted by any classifier). Ultrastop and tier 3-4 genes showed reduced ribosome density not only in SCR extensions but also across canonical CDSs, indicating generally lower expression. In addition, tiers 1-2 exhibited a pronounced Ribo-Seq peak at the second stop codon (Figure 1F), whose magnitude correlated with SCR score (Supplementary Figure S9). This signal reflects ribosome accumulation at the second in-frame stop, supporting productive translation of the SCR extension.

Gene ontology enrichment analyses highlighted that tier 1 and 2 genes were significantly enriched for cell-surface receptors, vesicle-associated components, and genes involved in nervous-system function and developmental processes, signaling, and transcription regulation, whereas tier 3 and 4 showed no specific enrichment (Supplementary Figure S10). These results recapitulated and expanded analogous analyses previously performed on smaller SCR gene sets predicted by evolutionary conservation ^20^, and are consistent with SCR prevalence in neurons ^16,28,30^.

Because our dataset includes samples digested with different nucleases, we assessed digestion-related biases and found that nuclease choice affected ribosome profiling patterns but preserved SCR-associated termination signatures (Supplementary Figures S9, S11). Consistent with prior reports in *D. melanogaster* ^24,34^, we also observed substantial ribosome density in 5’UTRs (∼2-fold lower than canonical CDSs; 3’UTRs were ∼10-fold lower than CDSs, Supplementary Figure S12). Yet, this signal lacked clear 3-nt periodicity, and showed no enrichment nor depletion near annotated start codons, arguing against widespread productive translation (Supplementary Figure S13,S14). Importantly, 5′UTR ribosome aggregated density did not correlate with SCR score; instead, differences across tiers were largely explained by overall ribosome load on the transcript, indicating that 5′UTR density scales with overall translation and is not related to SCR (Supplementary Figure S15).

Altogether, our predictions substantially expanded the repertoire of *Drosophila* SCR, offering new opportunities to investigate their functional and mechanistic diversity. Tiers 1-2 included 1,018 SCR genes not previously identified by either conservation ^23^ or Ribo-Seq-based approaches ^24^. Notably, experimentally characterized cases were detected and consistently assigned high scores (Table 1).

**Table 1.** Well-characterized *Drosophila* SCR genes and their corresponding scores and tier classifications.

| gene name<br>(symbol) | SCR score<br>(max 15) | tier | reference |
| --- | --- | --- | --- |
| *headcase (heca/hdc)* | 12 | tier 1 | Steneberg et al. 2001<sup>26</sup> |
| *synapsin (syn)* | 12 | tier 1 | Klagges et al. 1996; Nuwal et al. 2010<sup>27,35</sup> |
| *ventral veins lacking (vvl/dfr)* | 14 | tier 1 | Zhao et al. 2021<sup>15</sup> |
| *traffic jam (tj)* | 14 | tier 1 | Karki et al. 2022<sup>16</sup> |
| *kelch (kel)* | 11 | tier 2 | Xue et al. 1993; Robinson et al. 1997; Hudson et al. 2021;<sup>28,29,36</sup> |

### SCR associates with specific nucleotide and protein sequence signatures

We analyzed sequence-related features in our gene sets, seeking characteristics associated with SCR (Figure 2). Since our scoring is meant to capture the likelihood of programmed SCR, features mechanistically or functionally implicated in SCR should exhibit a consistent gradient across tiers. This is the case, for example, of the length of 3’UTRs and SCR extensions, which was significantly larger in tiers 1-2 than in other genes, as expected (Figure 2E). As we assessed stop codon identity (Figure 2A), we noticed that top tier genes preferentially carried UGA as the readthrough stop codon (63% in tier1), consistent with previous observations. No strong preference was detected for the second stop, except an unexplained overrepresentation of UAA in the weakest tier. Note that, while the identity of the second stop codon varied between genes, it appeared more evolutionarily conserved in top scoring genes (Figure 1E). We then examined the stop codon context (Figure 2B). The UGA-C motif, widely reported as the leakiest stop context, showed a gradual increase in usage from low- to high- tiers. The C at +4 position (i.e. 3’-adjacent to the stop) exhibited a similar trend for all stop codons. Conversely, A at +4 showed the opposite trend across all stops, consistent with stronger termination, as reported ^37^. Strikingly, top scoring SCR genes tended to have an A at +5 (part of the known SCR motif UGA-CA[A/G]N[U/C/G]A ^14^), independently of their stop codon. Further downstream, positions +6 to +8 showed little nucleotide bias despite their proximity to the stop codon, whereas position +9 displayed a strong G enrichment in tier 1 genes across all three stop codons (Supplementary Figure S16). Upstream of the stop, we observed a stop-independent overrepresentation of A at position −2 of genes in all tiers except the top one, which may be a general feature of efficient termination in insects. Position −4 followed a gradient similar to that of position +4 (more C and less A in the top SCR sets) in genes with UAG and UGA stop codons, though with smaller extent. Finally, top tier UAA-ending genes exhibited extreme nt biases at positions −11 (A), −10 (G), and −5 (A), while UGA-ending genes showed strong enrichment for G at position −7 (Supplementary Figure S16). Overall, our analyses pinpointed SCR-dependent biases in the stop codon and its surrounding sequence.

**Figure 2.**
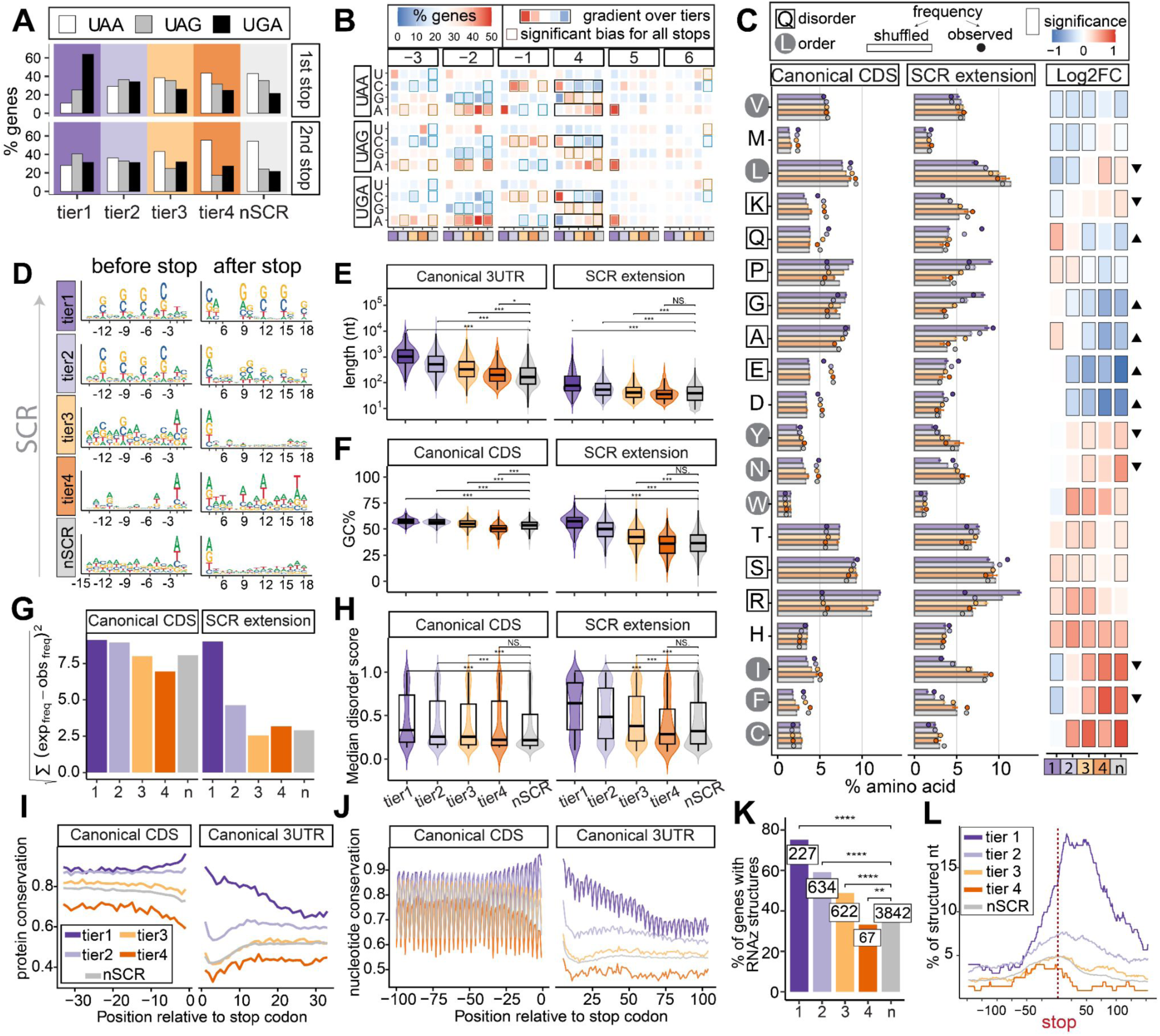
Signatures of SCR in insects. (A) Stop codon usage (color-coded) across tiers, for canonical CDSs and SCR extensions. (B) Stop codon context. For each tier (x-axis) and stop codon (facet rows), we show the percentage of genes (heatmap color) with each nucleotide (y-axis) at each position relative to the stop codon (facet columns). Positions are numbered relative to the stop codon: +4 is the first downstream nucleotide and −1 the last one upstream. Borders around tiles indicate statistically significant deviations from the expected 25% frequency that are consistent across all stops (FDR < 0.05 for each stop codon, exact binomial test; and combined Fisher’s aggregated FDR < 0.001). Black rectangles indicate positions identified by regression analysis, where nucleotide frequency across tiers was tested for association with SCR tier. Only the top 5 associations are highlighted (FDR < 0.001, binomial logistic regression trend test across tiers). Full regression results are provided in Supplementary Figure S16. **(C)** Amino acid composition in canonical CDSs and SCR extensions. The x-axis shows the percentage of each amino acid (y-axis) among protein sequences for each tier. Bars represent the expected amino acid frequencies from in-silico translation of shuffled nucleotide sequences within each tier (mean ± SD), while circles indicate the observed amino acid frequencies without shuffling. The third column represents the different prevalence of each amino acid between SCR extensions and canonical CDSs, expressed as log_2_ fold-change (log2FC). The x-axis shows SCR tiers; the y-axis shows amino acids. Borders around tiles indicate the statistical significance of the extension vs canonical comparison (FDR < 0.01, Fisher exact test). Triangles denote amino acids whose extension-versus-CDS log2FC correlated with SCR tiers (FDR < 0.05, linear regression trend test); upward indicates enrichment in top SCR tiers. Amino acids were ordered by hierarchical clustering based on their log2FC profiles across tiers (complete linkage method), and are additionally classified by disorder propensity ^39^: disorder-promoting residues are boxed, order-promoting residues circled, and others unmarked. (D) Sequence logos for each gene set (tiers and non-SCR genes) showing the last 15 nucleotides before the stop codon (left) and the first 15 nucleotides after the stop codon (right). (E) Distribution of log_10_-scaled lengths of SCR extensions and canonical 3’UTRs (defined as the mRNA region downstream of the canonical CDS) across tiers. (F) Distribution of GC content in canonical CDSs and SCR extensions across tiers. (G) Overall deviation from random amino acid composition across tiers. For each tier, we computed the Euclidean distance between the observed amino acid frequencies and those expected solely based on nucleotide composition, estimated from sequence shuffling. (H) Distribution of median AIUPred disorder scores in translated SCR extensions and canonical CDSs across tiers. (I) Mean protein conservation in the 30-codon stop codon context estimated from *Drosophila* WGAs (Methods). Average sequence identity values were calculated per gene, then aggregated across genes within each tier. **(J)** Mean nucleotide conservation in the 100-nt stop codon context, calculated analogously to panel (I). (K) Percentage of genes containing at least one conserved RNA structure (RNAz score > 0.8) located anywhere along the transcript. Absolute gene counts are shown in boxes. (L) Percentage of nucleotides that are base-paired in predicted conserved RNA structures near stop codons. *p < 0.05, **p < 0.01, ***, p < 0.001, ****p < 0.0001, NS: not significant; by two-sided Wilcoxon rank-sum test (E, F, H) or two-tailed Fisher exact test (K). Boxplots show median (center line), IQR (box), and 1.5×IQR whiskers.

Sequence logo analyses confirmed these trends (Figure 2D) and highlighted that tier 1-2 genes exhibit elevated GC content with pronounced 3-nt periodicity, peaking at third codon positions. This pattern was observed both upstream and downstream of the stop (except for the first two codons of the SCR extension), and resulted in significant compositional differences when comparing overall regional GC content across tiers (Figure 2F). These patterns are consistent with translational optimization of SCR genes through codon usage bias, as GC-ending codons are reportedly preferentially used and more efficiently translated in *Drosophila* ^38^. Of note, position −1 (i.e. immediately preceding the stop) did not exhibit GC enrichment, suggesting that other selective pressures drive nucleotide composition here.

These results were supported by complementary analyses of sequence conservation across *Drosophila* whole-genome alignments (WGAs) (Methods). Tier 1-2 genes showed increased nucleotide conservation compared to other genes in the last ∼25 nt of the canonical CDS peaking at the stop codon, with an even stronger trend in the ∼75 nt downstream (Figure 2J). Notably, conservation displayed clear 3-nt periodicity in top tier genes extending beyond the stop, consistent with purifying selection at the protein level. The +4 nucleotide was among the most conserved positions even in non-SCR genes, indicating a general constraint on termination. At the protein level, the amino acid preceding the stop was more conserved in tier 1-2 genes (Figure 2I), suggesting that P-site identity may influence SCR efficiency, although no specific amino acid enrichment was observed.

We also examined amino acid composition of SCR extensions relative to canonical CDSs and across tiers (Methods). Compared with translated shuffled controls that preserve nucleotide composition, canonical CDSs showed substantial deviations, consistent with selection acting at the protein level. This pattern was evident both for individual amino acids and using a global aggregated metric (Figure 2C,G). SCR extensions displayed weaker deviations overall, but these progressively increased toward higher SCR tiers, and most deviations occurred in the same direction as those observed in canonical CDSs (Figure 2C). This indicates that extensions of high-confidence SCR genes are subject to stronger protein-level selective constraints than those of other genes. Consistently, PhyloCSF scores, which quantify evolutionary signatures of protein-coding conservation ^18^, decreased with SCR tier and this effect was absent from alternative reading frames (Supplementary Figure S17). Within this general trend, we identified amino acids which were differentially represented in SCR extensions relative to canonical CDSs, with tier-dependent trends (Figure 2C, “Log2FC” facet): glutamine, glutamate, aspartate, glycine and alanine were enriched in the extensions of top tier genes, whereas leucine, lysine, asparagine, isoleucine and phenylalanine were depleted. Strikingly, enriched residues predominantly corresponded to disorder-promoting amino acids, while depleted residues were associated with ordered protein structure ^39^. This effect was especially pronounced for glutamine and glutamate, and could not be explained by nucleotide composition alone. We therefore quantified predicted disorder in SCR extensions (Methods) and found that extensions of top tier genes were indeed more disordered than non-SCR genes (Figure 2H). Together with the evidence of protein-level selection, these results suggest that intrinsic disorder is a widespread and potentially functional feature of programmed SCR in insects, possibly mediating regulatory effects (e.g., on protein interactions, stability, or condensate formation). Conversely, because hydrophobic residues at the C-terminus are associated with protein destabilization ^9^, their depletion in extensions of SCR genes may also contribute to protein stabilization.

### SCR may alter subcellular localization

To explore functional consequences of SCR, we compared predicted subcellular localization of canonical and SCR-extended proteins using DeepLoc2 (Methods). We identified 22 genes in which SCR was associated with a shift in localization, with a decreasing trend across SCR tiers (p = 0.005, one-sided Cochran-Armitage test). While previous studies searched for localization signals within SCR extensions ^24^, they did not examine whether the readthrough isoforms were predicted to relocalize relative to their canonical counterparts. We observed localization shifts between diverse subcellular compartments (Supplementary Figure S18; Supplementary Table T6). Altered localization represents a plausible functional outcome of SCR, and these predictions provide testable hypotheses for experimental validation. However, this feature appears restricted to a minority of cases rather than a general feature of *Drosophila* SCR.

### High throughput screening reveal new SCR stimulatory sequences

To investigate the molecular basis of programmed SCR, we set out to analyze readthrough-promoting *cis*-regulatory sequence elements. Since many recoding elements consist of structured RNAs ^1^, we used RNAz ^40^ to predict conserved RNA structures across WGA-derived transcript alignments of *Drosophila* species (Methods). Notably, genes with high SCR scores were enriched in conserved RNA structures (Figure 2K). Structured nucleotides preferentially accumulated in the window around the stop codon of tier 1-2 genes (from ∼25nt upstream to ∼75nt downstream), supporting the presence of localized RNA elements promoting SCR (Figure 2L).

We proceeded to experimentally test the SCR-stimulatory activity of candidate *cis-*acting elements using dual-reporter constructs, a widely employed strategy in SCR studies ^41^. These reporters contain two signal proteins, typically luciferases or fluorescent proteins, separated by an insert that includes an in-frame stop codon. The first signal serves as a marker of plasmid delivery and expression, whereas the second one reports translation beyond the stop codon. Thus, we can obtain a quantitative measure of SCR stimulation by inserting a candidate stop-containing sequence, and comparing signals to appropriate controls. These may include (i) a sense variant used for SCR rate normalization, and (ii) a frameshifted variant carrying a 1-nt insertion adjacent to the stop codon as a negative control; signal from this construct implies confounding processes, e.g. cryptic splicing, internal transcription initiation, or translation (re)initiation, rather than genuine SCR. Our constructs also incorporate two codon-diversified T2A peptide sequences flanking the stop codon region. These viral recoding elements prevent formation of a peptide bond, separating protein products and thus avoiding fusion-dependent artifacts in readthrough assays ^42^.

After validating our first dual-fluorescent reporter through pilot experiments with the known SCR stimulator from the *tj* gene ^16^ (Supplementary Figure S19), we scaled up this approach to a genome-integrated screening in *Drosophila* cells, inspired by recent CRISPR screens ^43,44^ (Methods). Briefly, we employed a AttB-flanked iRFP-eGFP reporter with an internal variable region positioned between T2A peptides (Figure 3A, Supplementary Figure S20) and carrying a library of 10,029 fixed-length (129-nt) inserts designed as described below (Figure 3B). The plasmid library was delivered and integrated into S2R+ cells via recombinase-mediated exchange, replacing a preexisting genomic mCherry-encoding cassette. Following expansion and monitoring via flow cytometry (Supplementary Figure S21), cells were sorted into four fractions based on eGFP signal: ultra-low, low, medium, and high. DNA from each fraction and from unsorted cells was isolated in two biological replicates, followed by PCR amplification and sequencing of the inserts. Quantifying inserts across fractions enabled identification of putative SCR-stimulatory elements. Selected candidates were subsequently validated individually using a luciferase-based dual-reporter system, suited for precise quantification of SCR rate in both insect and mammalian cells (Figure 3C; Methods).

**Figure 3.**
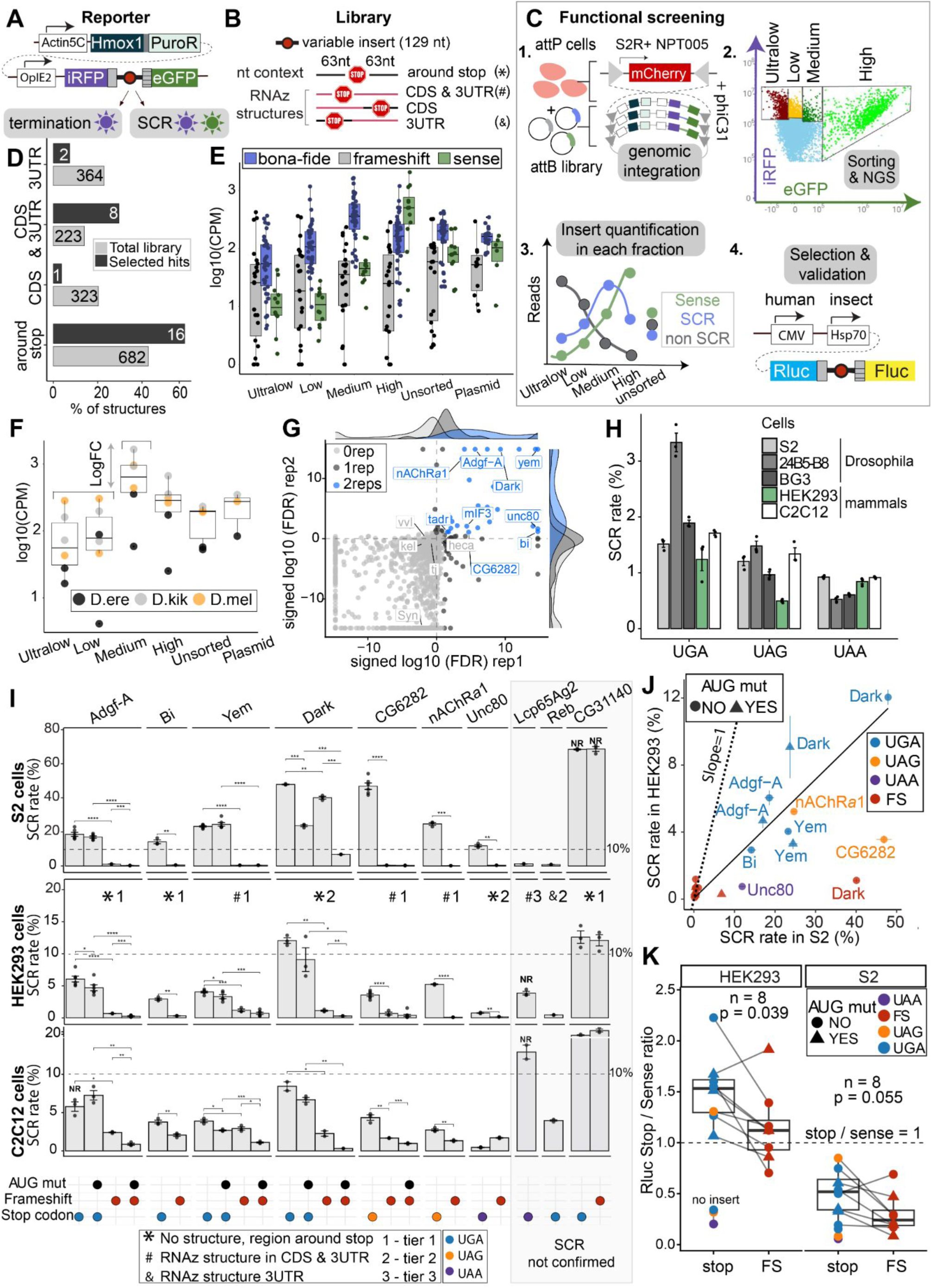
Identification of *cis*-acting SCR stimulators in S2 cells through functional screening. **(A)** Scheme of reporter designed for integration based screening in S2R+ cells. The Actin5C promoter drives the expression of Hmox1 and PuroR genes separated by a P2A peptide (dark grey rectangle). Hmox1 produces biliverdin, the metabolite required for iRFP fluorescence in insect cells. PuroR is a selection marker. The OpiE2 promoter drives the expression of dual fluorescent reporter. The variable insert containing the stop codon is flanked by codon-diversified T2A peptides (light grey rectangles). (B) Types of inserts tested in the library. All inserts were 129nt long, and consisted in symmetrical stop context windows for “around-stop” candidates (i.e. those other than RNAz hits). For stop-distal RNAz hits (i.e. “CDS” and “3’UTR” candidates), the native 6nt stop-codon context (black lines) was fused to the predicted structure (red lines), extended to reach the required fixed size. **(C)** Workflow of functional screening. The S2R+ insect cell line contains an integrated mCherry cassette flanked by attP sites, while the library backbone plasmid carries attB sites. Following PhiC31-mediated genomic integration, the reporter cassette replaces mCherry. After integration was completed and unintegrated plasmids were diluted through cell division, cells were sorted by flow cytometry. Cells were first gated based on iRFP, used as an integration marker, and then sorted according to eGFP signal, used as the SCR readout. Cells were collected into four fractions. Genomic DNA was extracted, and NGS libraries were prepared. For each insert, read counts were calculated in each sorted fraction. Inserts with no SCR activity were expected to be enriched in low-eGFP fractions, and SCR-stimulating inserts in high-eGFP fractions. Hits were selected based on the distribution of reads across fraction. Validated was performed using dual-luciferase constructs (Rluc: Renilla; Fluc: Firefly) driven by a dual-promoter system: CMV for expression in mammalian cells and Hsp70 for insect cells. (D) Composition of the screening library before screening (grey) and among selected hits (black). (**E**) Insert quantification across cell fractions of negative controls (frameshifted variants, grey), positive controls (sense variants, green) and a set of *bona fide*, validated SCR elements (from genes *Dark*, *miF3*, *tadr*; Methods). (**F**) Strategy for identification of hits from screening data. Shown are log_10_(CPM) counts for inserts derived from the SCR gene *Dark* in *D. melanogaster* (orange) and orthologous sequences from two additional *Drosophila* species (black and grey). Elements were classified as SCR stimulators when all evolutionary replicates showed consistent enrichment in the medium-eGFP fraction relative to the ultralow- and low-eGFP fractions (Methods). (**G**) Screening results and replicability. The x- and y-axes indicate hit strength in each sorting replicate, as “signed log10(FDR)”: the sign of the log fold change multiplied by −log10(False Discovery Rate). Lightgrey: structures not significantly enriched in either replicate; darkgrey: structures significantly enriched in at least one replicate; blue: structures significantly enriched in both replicates. Labels with blue color mark individually validated candidates; labels in grey indicate known SCR genes that did not reach significance in the screening. Significance threshold: FDR < 0.1 in both replicates (Methods). (**H**) Basal level of SCR (% relative to the sense version) measured from plasmids lacking SCR stimulators (n = 3) in 5 cell lines: *Drosophila* S2 (macrophage-like, derived from late embryo), 24B5-B8 (muscle-like, derived from late embryo), BG3 (central nervous system), human HEK293, and mouse C2C12. (**I**) Dual-luciferase validation of selected hits in *Drosophila* S2 cells and portability to human HEK293 cells and mouse C2C12 cells. NR indicates “non-reliable”, meaning that the observed Renilla values (i.e. the luciferase before the stop codon) were insufficiently low (Methods). Insert types and tier are indicated as defined in the bottom legend. Statistical comparisons are indicated (*p < 0.05, **p < 0.01, ***, p < 0.001, ****p < 0.0001, by one-sided Student’s t-tests: paired when n = 3 per group and samples were matched; otherwise unpaired). If AUG codons were present downstream of the stop codon, we generated versions in which they were mutated to UUG. Bars represent mean values and error bars indicate SEM. The number of replicate measurements varied between constructs (n = 2-6). (**J**) Correlation between SCR rates measured in S2 and HEK293 cells via dual-luciferase. Only elements that passed validation were included. The color indicates the stop codon of the various constructs, with red (”FS”) indicating frameshifted variants. Point shape indicates whether downstream AUG codons were mutated. The dotted line corresponds to a slope of 1, indicating identical SCR rates in both cell types. Correlation statistics: R = 0.7, p.value = 0.00039; or R = 0.9 and p.val = 3.14 * 10^-7^ when excluding *CG6282* UAG and *Dark* FS. (**K**) SCR-associated enhancement of reporter signal in S2 cells and HEK293 cells. Rluc signal (upstream of the stop codon) was normalized to the corresponding sense construct and compared between SCR constructs and their paired frameshift (FS) variants. P-values were calculated using paired Wilcoxon tests. Boxplots show median (center line), IQR (box), and 1.5×IQR whiskers.

The library was designed to test four classes of candidate *cis-*elements from our *Drosophila* SCR gene dataset (Figure 3B). Briefly, these included: (i) RNAz-predicted secondary structures overlapping the stop codon (”CDS-3’UTR” class); (ii) RNAz structures in the canonical CDS, repositioned for testing near their native 6-nt stop-codon context (”CDS”); (iii) RNAz structures located downstream of the stop, similarly repositioned (”3’UTR”); and (iv) the stop codon context (−63 to +63 nt) of genes in our SCR dataset lacking detectable conserved RNA structures (”around stop”). Altogether, the library comprised 1,592 native elements from *D. melanogaster*. In addition, it included multiple related inserts derived from each native element (Methods). Most importantly, these included two variants carrying non-native stop codons, as well as two or more evolutionary replicates, i.e. orthologous sequences from other *Drosophila* species extracted from WGAs. Evolutionary replicates enabled detection of conserved functional signals, under the assumption that functional programmed-SCR elements preserve activity across species. The library also included a set of manually selected *bona fide* elements from genes *Dark*, *miF3*, and *tadr*, whose SCR activities were independently validated by dual fluorescent reporter and Western blot prior to screening (Supplementary Figure S19). Lastly, the library included 6 sense and 11 frameshifted inserts serving as positive and negative controls, respectively.

Initial analyses recovered the expected enrichment of sense controls in the high fraction, and of *bona fide* SCR elements in the medium fraction, but also revealed poor replicate concordance and no depletion of frameshifted controls (Figure 3E, Supplementary Figure S22). We attributed this to multiple plasmid integrations per cell, possibly due to multiple AttP loci in S2R+ cells or off-target integration, exacerbated by extensive puromycin selection. This issue resulted in false-positive signals through hitchhiking with active inserts, and prevented the comprehensive quantitative SCR estimation of all library inserts. Nevertheless, evolutionary replicates provided an internal filter against such artifacts, since hitchhiking is expected to occur randomly among inserts. We therefore required multiple orthologous versions of the same *D. melanogaster* element to reproducibly enrich in higher fluorescence fractions (Figure 3F, Methods), thereby defining a small stringent set of screening hits. Of the 1,095 elements with sufficient coverage, 27 hits were enriched in the medium fraction and one in the high fraction (Figure 3G, Supplementary Table T7), collectively corresponding to 22 genes. Most candidates mapped to regions in the proximity of the stop codon, with only 2 derived from distal 3’UTR elements (Figure 3D). Notably, sequence logo analysis of the 6-nt context on each side of the stop codon of selected candidates revealed the known SCR motif CA(A/G)N(U/C/G)A ^14^ (Supplementary Figure S23), consistent with the nucleotide biases across SCR tiers presented earlier. The upstream region showed enrichment for the previously unreported CACNNA motif.

We validated screening hits by individually cloning 10 selected inserts into the dual-luciferase reporter (Supplementary Figure S24), additionally testing frameshifted variants as negative controls and sense variants for SCR rate normalization. For inserts containing downstream AUG codons, we also generated AUG to UUG mutants. We argue that these controls allow to exclude all aforementioned possible sources of artefactual signal. Transfection assays in *Drosophila* S2 cells showed that the stop-overlapping elements from *Adgf-A*, *bi*, and *yem* promoted robust SCR at rates of 15-25% (Figure 3I). By contrast, CG31140 was deemed a false positive: its apparent SCR matched the frameshift control and showed strongly reduced first-luciferase signal (Supplementary Figure S25), consistent with splicing-mediated skipping to the second reporter. The *Reb* element, which was predicted in the 3’UTR at a distance of 93 nt from the stop, also failed to replicate, showing no detectable activity. The gene *Dark* presented a more complex scenario: though the native construct reached ∼50% apparent SCR, elevated signal was also observed in the frameshift control. Mutation of downstream AUG codons reduced activity in both constructs (to ∼20% and ∼6.8%, respectively), suggesting that downstream initiation contributes, yet only partially, to the observed activity. Also considering that analogous assays in mammalian cells (see below) showed specific signal, we concluded that this element genuinely stimulates SCR; yet, additional *Drosophila*-specific RNA-processing or translational events likely occur and remain to be investigated. Next, the UAG-containing elements from *nAChRα1* and CG6282 showed robust SCR at rates of 23% and 50%, respectively. Among UAA candidates, *unc80* induced ∼10% SCR, while *Lcp65Ag2* did not validate. The latter showed reduced first-luciferase output (Supplementary Figure S25,S26), pointing to possible splicing artefacts, and was also weakly supported by Ribo-Seq (SCR score 5, tier 3). Overall, 7 of 10 tested elements validated in *Drosophila* cells, identifying efficient SCR stimulators across all three stop codons.

Following validation, we revisited screening data and analyzed constructs carrying alternative stop codons for each validated stimulator. Although the absence of evolutionary replicates precluded formal statistics, these data enabled a qualitative assessment of stop codon specificity of validated SCR stimulators (Supplementary Figure S27,S28). Native UAG (*nAChRα1*, CG6282) and UAA (*unc80*) stimulators apparently retained activity across stop codons. In contrast, most UGA-containing elements (*Adgf-A*, *bi*, *tadr*, *Dark*) showed reduced SCR stimulation with UAG and none with UAA (except the UGA-native *mif3* and *yem* elements, which remained active with any stop). These results may suggest greater stop-codon flexibility of UAA/UAG stimulators and stronger context dependence among UGA stimulators; yet, broader experimental tests will be needed to determine whether this pattern generalizes.

### Most *Drosophila* SCR elements are portable to mammals

To test whether insect SCR elements preserved their activity in mammalian systems, we performed dual-luciferase assays using the same set of plasmids described above in HEK293 (human embryonic kidney) and C2C12 (mouse myoblast) cells (Figure 3C,I and Supplementary Figures S25,S26,S29). Remarkably, six of the seven validated elements retained SCR activity in both mammalian cell lines. The sole exception was the UAA-containing element from *Unc80*, which showed no SCR in C2C12 cells and only weak activity in HEK293 cells. Importantly, readthrough rates were remarkably lower in mammalian cells than *Drosophila* cells across all constructs, not exceeding ∼15% in HEK293 and 10% in C2C12 cells. Nevertheless, we observed a striking correlation between SCR rates in *Drosophila* and mammalian cells (Figure 3J, Supplementary Figure S30). The correlation was even stronger between mouse and human cells, suggesting that the determinants of efficiency of SCR elements are largely shared across mammals. Our experiments indicate that the SCR-stimulating elements identified by our approach are largely portable to mammals, while also suggesting the presence of *trans-*factors that boost the magnitude of SCR efficiency in *Drosophila* cells.

### Translation termination is not inherently inefficient in *Drosophila*

Given their higher SCR rates, we next asked whether inefficient termination is a general feature of *Drosophila*. To estimate baseline readthrough in the absence of *cis* elements, we tested the three stop codons in the same short local context (GGUACC - stop - CGAUCC) in three *D. melanogaster* cell lines: S2 (macrophage-like, derived from late embryo), 24B5-B8 (muscle-like, derived from late embryo), and BG3 (central nervous system); and the two mammalian cell lines HEK293 and C2C12. The context included the SCR-prone C at +4, a feature shared by all sequence elements that passed validation. Measured baseline readthrough rates were similar across all cell lines (∼0.5–2%), except for UGA in 24B5-B8 cells (∼3.3%; Figure 3H, Supplementary Figure S31). Stop codon leakiness generally followed the expected order UGA > UAG > UAA, with HEK293 cells as the sole exception (UGA > UAA > UAG). Thus, in the absence of a dedicated *cis*-acting

SCR stimulator, differences between cell lines were modest, consistent with similar termination efficiencies. This indicates that *Drosophila* cells are inherently more responsive to SCR stimulation via *cis* elements than mammalian cells, rather than globally inefficient at terminating.

### Proteomics identifies peptides supporting stop codon readthrough

To expand experimental support for SCR, we analyzed publicly available proteomics data. Unlike Ribo-Seq, proteomics provides direct evidence of translated products and may therefore identify the amino acids incorporated at stop codons during readthrough. We collected 123 publicly available *D. melanogaster* tandem mass spectrometry (MS/MS) datasets (Supplementary Table T8) and searched them against custom databases allowing all 20 amino acids at stop codons, enabling detection of peptides mapping to canonical CDSs and SCR extensions (Methods). We identified 231 proteins with peptides in C-terminal extensions, ∼50% of which were also supported by Ribo-Seq (Figure 4A). The remaining cases were mostly supported by peptides spanning the stop codon and exhibited very short extensions, limiting their detection by Ribo-Seq (Supplementary Figure S32). Notably, higher-confidence tiers were more frequently detected and supported by a greater number of peptides (Figure 4B), largely reflecting their longer extensions and higher SCR activity. Overall, the majority of SCR peptides originated from head-derived samples, though contributions from other tissues were also observed (Figure 4F). Intersecting proteomics, Ribo-Seq, and conservation evidence yielded 54 top-confidence SCR genes (Figure 4D).

**Figure 4.**
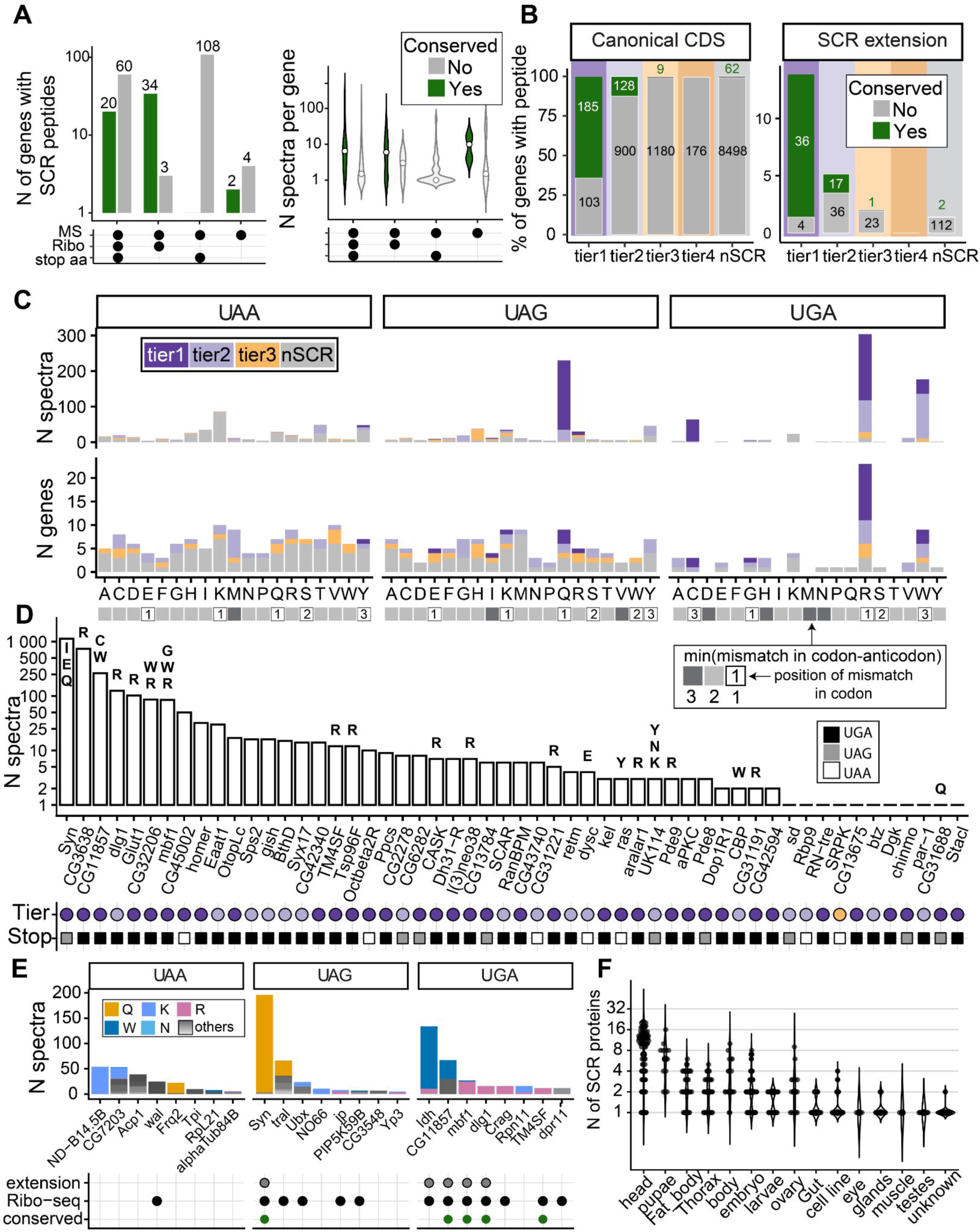
Proteomics reveals amino acids incorporated at stop codons during SCR. **(A)** Left: number of SCR proteins identified by proteomics (MS), stratified by SCR detection in ribosome profiling (Ribo) and by the presence of peptides enabling identification of the amino acid incorporated at the stop codon (stop aa). Colors indicate whether the gene was predicted to undergo SCR based on evolutionary conservation. Right: distribution of number of spectra per protein for detected SCR proteins. White circles indicate median values. (B) Percentage of genes with SCR detected by proteomics per tier, relative to canonical CDSs. Colors indicate conserved SCR cases. Absolute gene counts are shown on the plot. (C) Amino acid substitution profiles for each stop codon, colored by SCR tier. The upper panel shows the total number of spectra from all SCR genes; the lower panel shows data aggregated by gene. Below, the minimal nucleotide distance between the stop codon and cognate codons for each amino acid is shown, color-coded. For cases with a single nucleotide mismatch, its position within the stop codon is indicated (1, 2, or 3); e.g. “1” denotes that the first nucleotide of the stop codon differs from the corresponding nucleotide in the best-matching cognate tRNA for that amino acid. **(D)** Genes with SCR detected by all three approaches: conservation, proteomics, and ribosome profiling. Letters indicate the amino acids identified at the stop codon position, if any. Below, the tier and stop codon of each gene are indicated. (E) Top eight genes by number of spectra for each stop codon, showing the amino acid incorporated at the stop. Bottom annotations indicate whether each gene was predicted to undergo SCR by Ribo-Seq or conservation. “Extension” indicates the detection of additional peptides ascribed to the SCR extension but not spanning the stop codon. (F) Violin plot showing the number of SCR proteins detected by MS via peptide identification in the SCR extension, per sample, stratified by sample type. Sample types were manually curated from raw proteomics metadata and are ordered by decreasing SCR detection.

### Proteomics reveals candidate decoding mechanisms at stop codons

MS/MS data allowed us to characterize amino acid incorporation at stop codons (Methods), although sensitivity was limited by the shallow coverage of shotgun proteomics and low abundance of SCR products. UGA yielded the clearest signal, showing predominant incorporation of arginine and tryptophan consistently across genes and spectra (Figure 4C). In contrast, other stops lacked consistent preferences. UAA showed no specific bias. For UAG, glutamine incorporation was supported primarily by the known SCR gene *synapsin* (Figure 4E).

To investigate decoding mechanisms, we classified amino acid incorporations according to the most parsimonious stop-anticodon pairing model (Figure 4C). Whereas tryptophan incorporation at UGA is explained by classical third-position wobble, arginine at UGA requires a mismatch at the first codon position, generally considered to be under strong decoding constraints. Interestingly, peptides involving first or third codon position mismatches showed higher identification confidence scores for all stop codons (Supplementary Figure S33), suggesting that such decoding mechanism may not be restricted to UGA. In agreement, glutamine incorporation at UAG (which implies a first-position mismatch) was previously reported for the SCR gene *vvl* ^15^. Altogether, these results provide clues into stop codon decoding during insect SCR, possibly involving non-canonical wobble pairing. Yet, these patterns are based on sparse proteomic evidence and should be interpreted cautiously until confirmed through targeted experiments.

### SCR activity varies across tissues and developmental stages

Given the biological diversity of our Ribo-Seq dataset, we asked whether SCR was sample-specific or restricted to particular genes under defined conditions. We developed a single-sample detection framework based on Ribo-Seq metrics, following criteria analogous to Dunn *et al* ^24^. To avoid expression-dependent biases, we used thresholds scaled by expression and calibrated on conserved SCR genes (Methods). In the absence of detectable SCR, genes were classified as either expressed without readthrough or insufficiently expressed for detection, using a canonical CDS threshold of 10 FPKM set by exploratory analyses (Methods, Supplementary Figure S34). We thus selected 881 tier 1-2 genes with SCR in ≥2 samples and performed hierarchical clustering based on expression and SCR patterns (Methods, Figure 5).

**Figure 5.**
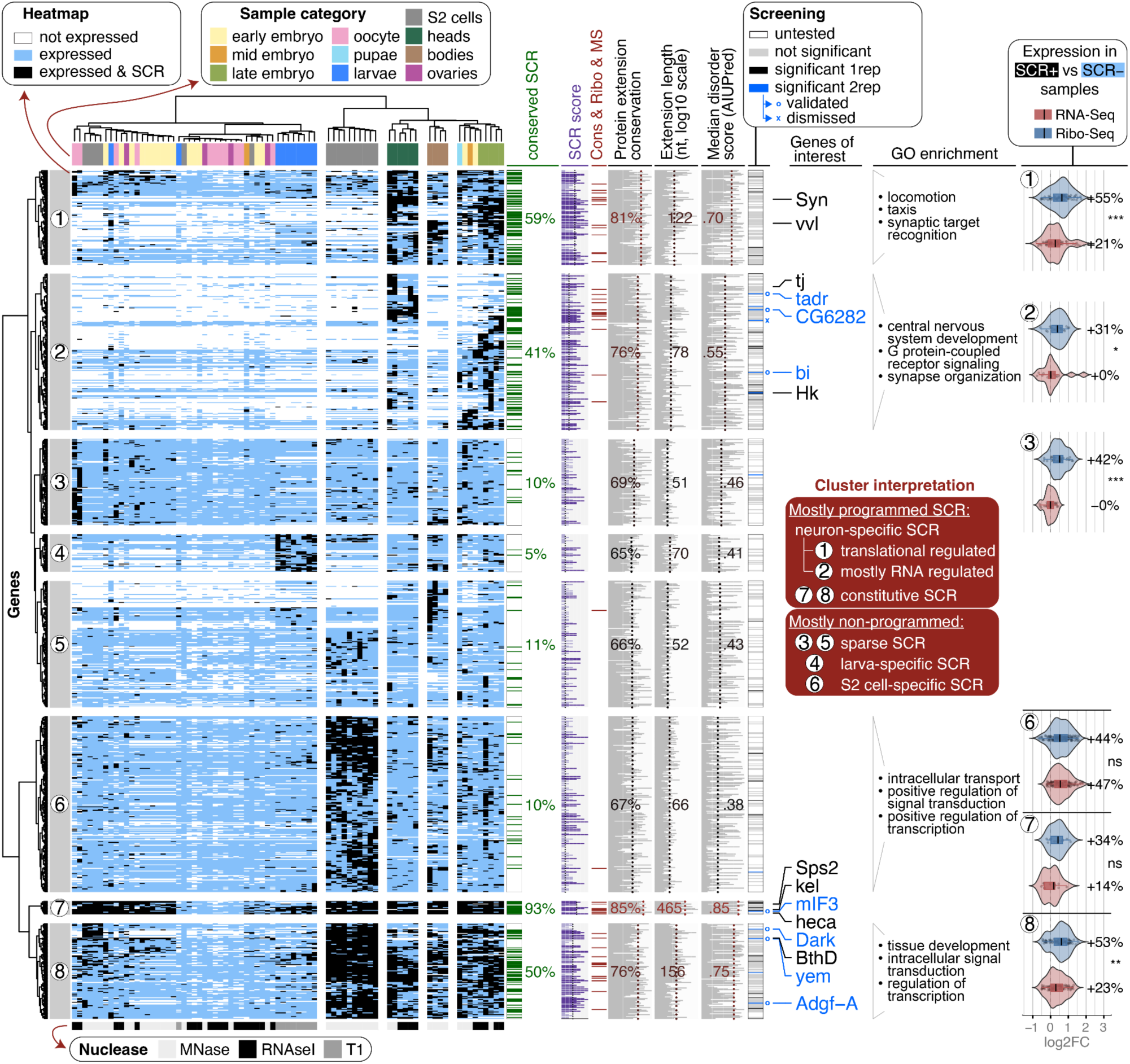
Regulatory landscape of SCR activation across tissues and developmental stages. Clustered heatmap summarizing sample-specific SCR detection. Rows correspond to genes (tier1-2 detected in at least 2 samples) and columns to Ribo-Seq samples. Sample category and nuclease treatment are indicated by colored bars at the top and bottom, respectively. Heatmap colors denote expression and SCR status: white, low or absent expression (insufficient for reliable SCR detection); blue, expressed without SCR; black, SCR detected (Methods). A heatmap of SCR rates is provided in Supplementary Figure S35. Hierarchical clustering resolved eight gene clusters (left), whose interpretation is summarized in the red inset. Vertical annotation tracks show gene-level features and their aggregated values within each cluster: conserved SCR genes; SCR score; SCR genes predicted by conservation, Ribo-Seq and proteomics; protein conservation of the SCR extension; extension length; median extension disorder (AIUPred); and functional-screening results (legend on top). For quantitative features, cluster medians are indicated by dashed lines and labels, color-coded according to their relative rank among clusters (black, lowest; red, highest). Characterized SCR genes are labeled on the right; *Bthd* and *Sps2* are selenoproteins ^46^. Representative enriched Gene Ontology categories are shown for clusters with significant enrichment. The rightmost panel summarizes expression changes associated to SCR. Violin/boxplots show per-gene log2 fold changes in RNA-seq and Ribo-Seq expression, calculated from mean expression values in SCR-positive (black cells in heatmap) vs SCR-negative (blue cells) samples. Percentages indicate median increases, and significance was assessed by paired Wilcoxon tests with BH correction (*p < 0.05, **p < 0.01, ***, p < 0.001, ****p < 0.0001, NS: not significant). Only clusters containing at least 12 genes with ≥6 SCR-positive and ≥6 SCR-negative samples were analyzed; extended results are shown in Supplementary Figure S36.

Samples clustered into two major groups. The first comprised ovaries, oocytes, and early embryos, and showed few SCR events. The second, included S2 cells, adult heads and bodies, and late embryos, and was markedly enriched in SCR. Clustering reflected biological context rather than nuclease type (Figure 5, bottom). Overall, SCR was detected in a wide range of tissues and developmental stages. Given the reported abundance of SCR in neurons, the high prevalence observed in heads and late embryos was expected. However, S2 cells exhibited even more SCR-positive genes, with only partial overlap with neuron-rich samples. This pattern is unlikely to be explained by gene expression, as most S2-specific SCR genes were sufficiently expressed in heads to permit detection. Consistently, Ribo-Seq-based estimates indicated substantially higher SCR rates in heads and S2 cells than in other sample types, although with considerable heterogeneity among genes (Supplementary Figure S35).

### Distinct clusters of SCR genes reveal sample-specific activation and functional signatures

To further characterize the regulatory landscape of SCR, we clustered SCR genes based on the framework described above (Methods). This resolved eight major groups with distinct SCR profiles across samples, which we then examined for signatures of programmed SCR and context-specific regulation (Figure 5). Conserved SCR genes, together with features associated with programmed SCR (score, extension conservation, and extension length), were strongly enriched in clusters 1, 2, 7, and 8. Extension disorder was likewise elevated across all these clusters.

Clusters 1 and 2 appeared to represent neuron-specific programmed SCR. They included known neuron-associated SCR genes such as *synapsin* (*Syn*) ^27,35^, *ventral veinless* (*vvl*) ^15,45^, and *traffic jam* (*tj*) ^16^ and were enriched for neuronal and synaptic functions. SCR activity in these clusters was detected primarily in heads and late embryos. Notably, genes in cluster 1 were broadly expressed across tissues but underwent SCR mainly in neuron-rich samples, suggesting translational regulation. In contrast, genes in cluster 2 showed more restricted expression patterns that largely coincided with SCR occurrence, consistent with predominant transcriptional control.

Clusters 7 and 8, which also exhibited prominent features associated with programmed SCR, displayed widespread and robust SCR activity across samples. Cluster 7 exhibited constitutive SCR across nearly all stages and tissues except oocytes, whereas cluster 8 was active across most samples within the SCR-rich group. These clusters included well-characterized examples such as *kelch* (*kel*) ^28,36^, *headcase* (*heca*) ^26^, and selenoproteins *BthD* and *Sps2* ^46^; cluster 8 was enriched for functions related to tissue development, signalling, and transcriptional regulation. We interpret clusters 7 and 8 as constitutive programmed SCR.

In contrast, clusters 3-5 exhibited sparse or context-restricted SCR (cluster 4 predominantly in larvae), weak SCR-associated features, low conservation, and no significant functional enrichment. These properties suggest that they consist primarily of non-programmed SCR, together with a subset of genuine programmed events that escaped robust detection due to limited sample coverage. Lastly, cluster 6 showed SCR activity largely restricted to S2 cells and enrichment in intracellular regulatory functions; however, its low conservation and restriction to cell culture make its biological significance difficult to interpret.

Results from the functional screening independently supported this classification (Figure 5). Nearly all screening hits (and all validated stimulators) mapped to clusters 2, 7, and 8. Only two hits (untested in validation) originated from clusters 3-6, reinforcing the interpretation that these clusters are largely dominated by non-programmed SCR. Most strikingly, cluster 1 yielded no screening hits despite its strong enrichment in conserved SCR genes. This observation is consistent with a model in which efficient SCR in this cluster depends on neuron-specific *trans*-acting factors absent from S2 cells, rather than autonomous *cis*-regulatory elements. Altogether, these analyses begin to resolve the apparent heterogeneity of insect SCR, revealing distinct regulatory modes and functional classes within the broad set of SCR genes.

### SCR boosts protein production

Inspection of raw signals from the dual-luciferase assays (Figure 3I) revealed a striking and informative pattern. Across all *Drosophila* and mammalian cell types tested, the first luciferase signal was consistently higher in SCR constructs than in frameshift controls (Figure 3K, Supplementary Figure S37B). The relationship to sense controls varied between systems: in S2 cells, both SCR and frameshift constructs showed markedly reduced signal relative to sense controls; in mammalian cells, frameshift constructs were largely indistinguishable from sense controls, while SCR constructs showed similar (C2C12) or slightly elevated (HEK293) signal (Figure 3K, Supplementary Figures S25,S26,S29,S37). Observed across multiple candidates and independent transfections, these results suggest that SCR enhances overall protein output from the mRNA, including the canonical product, particularly in *Drosophila* cells.

This boosting effect may arguably result from increased transcript stability and/or enhanced translation. A major determinant of mRNA stability is NMD, which degrades transcripts undergoing premature translation termination. NMD operates through two alternative mechanisms: an exon junction complex (EJC)-dependent pathway predominant in mammals, and an EJC-independent pathway determined by termination-poly(A) distance, common in other eukaryotes including insects ^47^. The sharp difference in the first luciferase signal between frameshift (lowest), SCR (mid), and sense (highest) constructs in S2 cells is consistent with EJC-independent NMD activity, since transcript stability is expected to increase as termination occurs closer to the poly(A) tail. This is in agreement with previous observations that knockdown of the NMD factor *Upf1* in fruit flies increased the mRNA levels of a SCR reporter ^30^. Fittingly, the signal of frameshift constructs resembled sense controls in HEK293 cells, where this mechanism is reportedly much weaker; yet, SCR constructs still showed increased output, which cannot be explained by transcript stabilization alone. This led us to hypothesize that SCR may boost protein output also through enhanced translation.

To explore this hypothesis, we leveraged our single-sample SCR calls. Focusing on tier 1-2 genes in Ribo-seq datasets with matched RNA-seq, we compared samples in which the same gene did or did not undergo detectable SCR (Methods; Figure 5, Supplementary Figure S36). On average, SCR occurrence was associated with a 25% increase in RNA-seq levels, and a significantly higher increase of 43% in Ribo-Seq signal. These effects varied across gene clusters, with programmed SCR clusters 1 and 8 showing the strongest Ribo-Seq increases. We next examined translation efficiency, defined as the ratio between Ribo-Seq and RNA-seq signal (Supplementary Figure S38). Around the stop codon, translation efficiency was significantly elevated in SCR-positive samples, as expected from delayed termination and ribosome pausing at readthrough-prone stops. Importantly, a similar trend remained evident across the canonical CDS after excluding regions proximal to the start and stop codons, and was more pronounced in tier 1 than tier 2 genes. These observations suggest that SCR occurrence is associated with enhanced ribosome occupancy beyond the termination region itself.

Although SCR detection biases may contribute to the observed Ribo-Seq patterns, these results align with our reporter assays and support a model in which SCR promotes overall protein production through both transcript stabilization and translational enhancement. Dedicated experiments will be required to validate this model and dissect the underlying mechanisms.

### Introducing the *Drosophila* Stop Codon Readthrough Atlas

Our findings reinforce *Drosophila* SCR as a widespread and functionally diverse regulatory phenomenon. We argue that its impact should be considered on a gene-by-gene basis to understand its role in diverse aspects of insect biology. To enable this, we developed the *Drosophila* Stop Codon Readthrough Atlas, a webserver that integrates all SCR-related data into a unified, gene-centric resource (https://mariottigenomicslab.bio.ub.edu/dmel_scr_atlas/). Users can query genes by symbol or FlyBase identifier and explore a wide range of information (Figure 6), including SCR score and tier, nucleotide and amino acid sequences, evolutionary conservation and WGAs, predicted disorder and subcellular localization, 3D structure predicted by AlphaFold2, aggregated and sample-specific Ribo-Seq coverage, proteomic support, conserved RNA structures, and results from reporter-based assays of candidate SCR elements. Altogether, this resource provides a unified platform to explore, interpret, and prioritize SCR events, and we expect it to accelerate functional and mechanistic studies in *Drosophila*.

**Figure 6.**
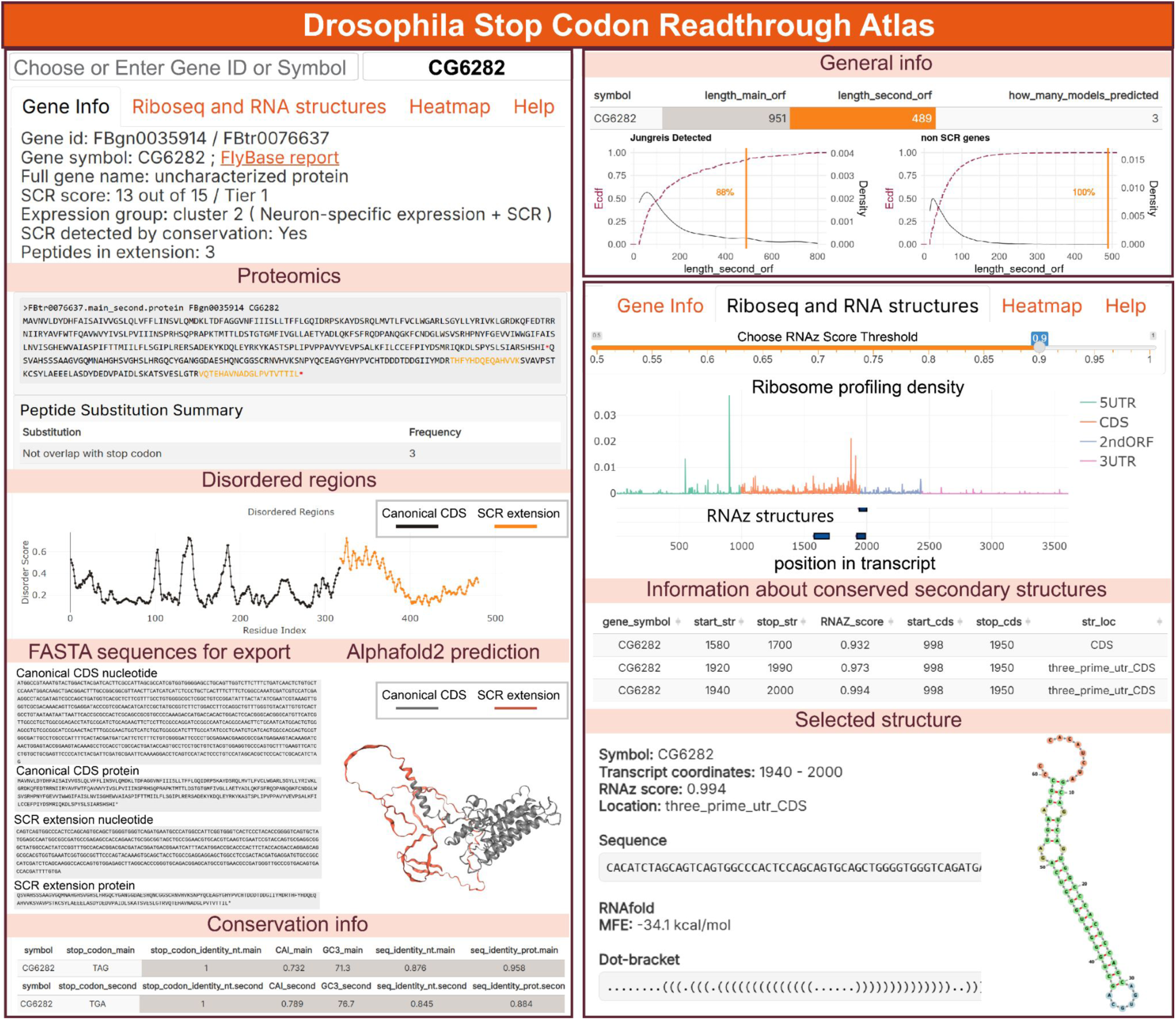
The *Drosophila* Stop Codon Readthrough Atlas. Users can enter a FlyBase gene ID or gene symbol in the query field. Information is provided for all candidate SCR genes identified by Ribo-Seq (any tier; n = 2,848). The example shown corresponds to gene CG6282, identified in the screen; panels and content were rearranged for visualization. The Atlas is organized into four tabs. The first tab reports: SCR classification results (score and tier), proteomics results highlighting SCR extension regions with matched peptides, disorder scores across the extension, nucleotide and protein sequences of the canonical CDS and SCR extension, conservation-related metrics, AlphaFold2 structure predictions for extended protein isoforms (available for candidates with SCR extensions longer than 40 amino acids) and more (not shown). For each metric, distributions across all genes and conserved SCR genes can be visualized upon selection (see top-right). The second tab is dedicated to Ribo-Seq and RNA structures (right): aggregated nucleotide-resolution density profiles across the transcript (without noise correction), together with conserved RNA structures predicted by RNAz with an adjustable score threshold. Clicking on a structure reveals its sequence and predicted secondary structure by RNAfold ^48^. The third tab reports activation patterns, and corresponds to a heatmap visualization analogous to Figure 5, wherein the queried gene is highlighted; this is not shown in this Figure. The heatmap tab is available only for tier1-2 genes whose SCR was detected in ≥2 samples; gene absence may reflect insufficient expression in individual datasets. The fourth tab is a help page (not shown).

## Discussion

Previous research has established SCR as an abundant phenomenon in insects supported by evolutionary conservation ^19,20,23,25^ and limited genome-wide experimental data ^24^, and highlighted tissue-specific patterns ^16,28,30^. Here, we provide the most comprehensive characterization of *D. melanogaster* SCR to date through a multi-omics integration approach.

A major obstacle to SCR discovery in insects is the limited quality of available Ribo-Seq data. Compared to mammalian and yeast studies, insect Ribo-Seq suffers from suboptimal protocols resulting in reduced coverage, weaker triplet periodicity, and overall lower signal-to-noise ^49^. Despite these limitations, our framework was able to detect reliable SCR events at genome scale through dataset aggregation, noise correction, machine learning, and integration with evolutionary conservation. To further expand and refine the regulatory landscape described here, broader sampling across biological conditions and improved protocols, particularly for efficient depletion of insect rRNA, will be required.

Predicted SCR genes recapitulated essentially all major signatures previously associated with programmed readthrough, including enrichment of UGA stop codons, cytosine at the +4 position, long 3′UTRs, elevated GC content, and disordered protein extensions ^14,20,22,23,32,50–52^. Moreover, they displayed characteristic Ribo-Seq patterns, including increased ribosome occupancy at the readthrough stop, sustained read density downstream, and termination peaks at the second in-frame stop. Interestingly, SCR genes displayed a diverse repertoire of motifs downstream of the stop codon. The most prevalent was the CANNNG motif, reminiscent of the CA(A/G)N(U/C/G)A context previously associated with readthrough in yeast ^14,51^ and TMV ^53^. A smaller subset of genes carried the CUAG motif, previously linked to mammalian ^22,52^ and viral readthrough ^53^.

However, the nucleotides flanking the stop codon capture only a fraction of the determinants of programmed SCR. Strikingly, evolutionary conservation peaked within a window surrounding the stop codon (approximately −25 to +75 nt), coinciding with a strong enrichment of conserved RNA structures. RNA structures are established mediators of translational recoding ^1^. While their enrichment in *Drosophila* conserved SCR genes had been previously reported ^20,54^, we here extended this observation by pinpointing the precise localization of this signal and showing that it is independent of transcript length. Together, these observations indicate that programmed SCR in *Drosophila* is commonly controlled by proximal *cis*-regulatory elements acting in concert with favorable stop-codon contexts. Importantly, because we discovered that stop codons lacking *cis*-stimulatory elements terminate with similar efficiencies in insect and mammalian cells, we argue that the evolutionary expansion of SCR in insects was driven not by relaxed termination, but by enhanced sensitivity to SCR elements. We propose that insects possess specialized regulatory mechanisms that amplify the effects of SCR stimulators, whose characterization now emerges as a central challenge for the field. We reason that *cis*-acting SCR stimulator elements may control termination by interfering with the binding of termination factors (e.g., via ribosomal conformational changes), and/or by altering ribosome dynamics at the stop codon (e.g., inducing stalling). Their effects may be exerted directly on the translating ribosome or indirectly through the recruitment of yet unknown *trans*-acting factors.

An important advance of our study is the ability to examine genome-wide SCR regulation across diverse biological conditions. While previous work established tissue-specific SCR in individual genes ^16,28,30^, its generality remained unclear. By combining sample-specific SCR detection with unsupervised clustering, we identified distinct regulatory patterns, including constitutive and tissue-restricted modes. We interpret constitutive SCR as largely driven by autonomous *cis*-elements, whereas tissue-specific SCR likely depends on additional *trans*-acting factors, particularly in neurons. This provides a framework for resolving the remarkable heterogeneity of insect SCR according to regulatory principles.

To experimentally determine SCR stimulators, we developed a dual-fluorescent SCR reporter system in *Drosophila* cells capable of interrogating thousands of sequences. Previous high-throughput reporter assays primarily examined disease-associated nonsense mutations and engineered stop-codon contexts and were restricted to human cells ^52,55^. To our knowledge, ours is the first screen targeting programmed SCR, and the first pooled functional assay of any kind in insect cells, other than the CRISPR-based screens that inspired it ^43,44^. The system has several limitations, including imperfect control of reporter copy number, restricted insert size, and evaluation in a single (non-neuronal) cellular context; slow cell division and lengthy optimization cycles hindered further refinement. Consequently, our screen likely captured only a subset of SCR stimulators, preferentially detecting strong and autonomous elements. This interpretation is consistent with the enrichment of validated hits among the constitutive SCR clusters and their ability to remain active in mammalian cells, albeit at reduced efficiency. This unexpected portability across large evolutionary distances highlights insects as a rich evolutionary reservoir of functional readthrough elements with potential biomedical and biotechnological applications.

The requirement for conserved activity across orthologous sequences probably excluded recently evolved regulators. On the other hand, positive hits likely represent functionally optimized drivers of genuine programmed SCR. Notably, the screen recovered SCR stimulators from genes not previously identified by evolutionary conservation (e.g., *Adgf-A*, *Dark* among our validated genes; additional cases listed in Supplementary Table T7), underscoring the value of direct functional interrogation. The diversity of sequences, stop codons, and predicted RNA structures among validated hits (Supplementary Figure S39) further supports the view that insect SCR is governed by multiple regulatory mechanisms.

Our analyses also provide new insights into the biological consequences of SCR. Functionally, our results suggest that SCR may play a dual role: (i) generating protein isoforms with distinct C-terminal regions that shape functional properties, and (ii) modulating the overall protein production level. Related to protein diversification, multiple functional paradigms emerged. In some cases, SCR introduces recognizable features such as localization signals (as well documented ^56,57^) or structured elements (e.g., α-helices), but these appear case-specific rather than generalizable. In contrast, recurrent compositional patterns point to a broader functional trend: SCR extensions are enriched in disorder-prone amino acids, in particular glutamine. Notably, intrinsically disordered regions are well known to act as interaction platforms and to contribute to transcriptional regulation and phase separation ^39,58^, making these plausible functions of SCR extensions. Strikingly, SCR genes are enriched in functions related to transcriptional regulation, signaling, development, and synaptic processes, suggesting that the biological impact of SCR is amplified through downstream transcriptional and signaling cascades.

Related to the impact on gene expression, SCR may modulate protein output at both the transcript and translation levels. Our reporter assays in S2 cells are consistent with SCR decreasing mRNA sensitivity to EJC-independent NMD. Supporting this interpretation, Chen et al. (2020) showed that knockdown of NMD factor increased expression of SCR reporter constructs. Yet, the same study found little effect on an endogenous SCR transcript, highlighting that reporter architectures may artificially magnify NMD-mediated regulation ^30^. This caveat may also apply to our system, as the distance between canonical and SCR termination sites is substantially larger in our dual-luciferase reporter (1650nt) than in endogenous SCR genes (median 612nt among tier 1-2 genes). Following observations from reporter assays in HEK293 cells, we hypothesized that SCR may also promote protein output through an effect on translation. Our genome-wide analyses provided preliminary support for this idea, although the evidence remains correlative. We speculate that factors promoting SCR may simultaneously enhance translation, with initiation factors representing attractive candidates given their established involvement in SCR ^59–62^. Alternatively, the increased protein output may reflect more efficient ribosome recycling following readthrough, owing to the decreased termination-poly(A) distance. Further experiments will be required to disentangle the interplay of transcript stability pathways, translational machinery, and SCR.

Beyond their mechanistic and functional implications, our analyses also provide insight into how programmed SCR may originate and evolve. We view programmed SCR as arising from sporadic readthrough events that are retained and progressively optimized when their protein products confer a selective advantage. In this framework, SCR score serves as a rough proxy for evolutionary maturity, ranging from recently emerged or non-programmed events to ancient programmed SCR. Along this axis, we observed progressive enrichment of SCR-promoting stop contexts, emergence of conserved downstream RNA structures, and amino acid composition shifts indicative of increasing protein-level selection. Remarkably, codon usage optimization toward GC-rich codons occurred not only within the SCR extension but also in the CDS upstream of the stop codon.

Overall, our work defines SCR as a widespread and functionally diverse regulatory layer in insects, whose impact must be considered on a gene-by-gene basis. To facilitate this, we provide a dedicated webserver for convenient data exploration, and we encourage its adoption by the broad *Drosophila* community. Concomitantly, our findings highlight several key unresolved questions about insect SCR. First, it will be important to experimentally test SCR determinants across cell types. Extending our pooled framework to additional *Drosophila* cell lines and improving reporter design using mammalian-inspired genetic tools ^52,63^ should substantially enhance this approach. At the mechanistic level, the *trans*-acting factors that promote efficient SCR in insects, and especially in neurons, remain largely unknown. The brain-enriched eRF1H isoform is likely only one component of a broader regulatory network ^16^, in which tRNA abundance, identity and modifiers emerge as prime candidates, particularly given the potential role of non-canonical wobble pairing proposed here. As noted, our results also underscore the importance of systematically dissecting the interplay between transcript stability and translation dynamics. From an evolutionary perspective, the diversity and distribution of SCR across insect species and related lineages remain only partially explored ^23^. Understanding how it varies across taxa, and how it has been gained, lost, or optimized during evolution, will be essential to place these findings in a broader evolutionary context. Finally, the structural basis of SCR remains largely unknown: while ribosome structure was resolved for Sec insertion ^8^, readthrough mediated by near-cognate tRNAs is still not captured by any tridimensional structure, and the structure of the relevant RNA elements was only predicted *in silico*. Together, these challenges provide a tentative roadmap for future studies aimed at understanding the mechanisms, evolution, and biological significance of SCR in insects.

## Methods

### QUANTIFICATION AND STATISTICAL ANALYSIS

#### Ribosome profiling

Fastq files for all available Ribo-Seq datasets obtained from *Drosophila melanogaster* (listed in Supplementary Table T1) were downloaded from the GEO database. Adapters were identified using DNApi (version 1.1) ^64^. Reads were trimmed with Cutadapt (version 2.8) ^65^ using the parameters --minimum-length 20, --maximum-length 40, and -q 20 to remove low-quality bases and retain reads of appropriate length. For the dataset GSE99920, trimmed reads were reverse-complemented using the FASTX-Toolkit to correct the library orientation prior to alignment. Trimmed reads were aligned to the *D. melanogaster* genome (FlyBase release dmel_r5.27) using STAR (v2.7.1a) ^66^, using the extended genomic annotations described below. STAR was run with the following options: “--alignSJDBoverhangMin 1 --seedSearchStartLmax 15

--outFilterScoreMinOverLread 0.66 --outFilterMatchNminOverLread 0.66 --outFilterMatchNmin 0

--outFilterMismatchNmax 3”.

To accurately assign ribosomal P-site positions, read-length-specific 5′-end offsets were estimated independently for each sample using the psite utility from the Plastid package (v0.5.1) ^67^. Offset predictions were visually inspected around annotated start codons and manually adjusted when the P-site peak was misaligned. Subsequent analyses were conducted in Python using the Plastid package as a library. BAM files were loaded as BAMGenomeArray objects, and P-site-adjusted read mappings were applied using a VariableFivePrimeMapFactory derived from the offset tables obtained as described above. In samples lacking clear periodicity (those that employed RNAse T1, GSE99920 ^68^), a centered mapping strategy (CenterMapFactory, nibble = 12) was employed, meaning that reads were trimmed 12 nt from both ends. Read counts were normalized to library size to allow comparisons across samples.

Finally, the ribosome profiling densities for each sample were calculated at each nucleotide position. Ribosome profiling density is defined as the library-size-normalized coverage of P-site assigned reads at each nucleotide position. For all datasets except one, only reads of length 28-35 nt were considered as individual-length aggregated profiles showed robust 3-nt periodicity in this range for most samples, and this interval contained the majority of the reads (Supplementary Figure S1 (left panel) and S41). A broader read-length range (20-40 nt) was used for the Chen dataset (GSE99920), wherein no clear P-site offset could be defined. We also examined other read-length ranges. Shorter reads, typically associated with ribosomes bearing an empty A-site, did not show detectable accumulation at stop codons relative to the CDS when considering all genes, and differences between non-SCR and tier 1-2 genes were proportional to overall CDS Ribo-Seq density (Supplementary Figure S40).

#### *Drosophila* genome annotation preparation

For Ribo-Seq quantification, we employed the Flybase annotation of protein coding genes release dmel_r5.27, edited as follows. First, to minimize false positives in SCR detection arising from splice isoforms or misannotations, we selected one representative transcript per gene: after excluding transcripts lacking a 5’UTR or 3’UTR and those with an internal stop codon in the CDS, we selected those with the 3’-most stop codon. In case of ties, we further prioritized, in order: transcripts tagged as *principal* in the APPRIS database (version 25.08_v50; assembly version: BDGP6; Gene Dataset: Ensembl96); transcripts with the longest mRNA, and transcripts with the highest mean Ribo-Seq density across all samples. Next, each transcript was divided into three regions: (i) the canonical CDS (annotated CDS), (ii) the SCR extension (from after the canonical stop codon up to the next in-frame stop), and (iii) the negative control region, i.e. downstream of the SCR extension (minimum of either the SCR extension length or 114 nt). For Ribo-Seq quantification, we excluded the first 18 nt and last 15 nt of the canonical CDS, as well as the first 6 nt of the SCR extension, since these regions may display artificially elevated density due to slow initiation and termination. For selenoproteins, we manually corrected the annotation so that the canonical ORF extended until the Sec-encoding UGA codon, which we used as positive controls.

Transcripts were filtered to retain only those in which both the SCR extension and the downstream negative-control region were fully contained within the annotated 3′UTR. After filtering, 9,158 of 12,989 genes were retained for further analysis.

#### Aggregation of datasets and noise removal

After Ribo-Seq data processing and mapping, ribosome density was calculated independently for each region, and several metrics to evaluate whether the extension was actively translated were computed (see below). Initially, we attempted to detect SCR occurrence independently in each sample as done by Dunn *et al* ^24^. Yet, after extensive manual inspection of the Ribo-Seq profiles of putative candidates, we concluded that the limited coverage of single-sample data resulted in many false positives (e.g. due to spurious translation-unrelated peaks in 3’UTRs) and false negatives; i.e. over 66% of conserved SCR genes were not detected. Therefore, to take advantage of the substantially larger collection of Ribo-Seq datasets analyzed in this study, we aggregated signals across samples to improve SCR detection power.

Our Ribo-Seq density quantification is naturally structured along three axes: (i) transcript (one representative transcript per gene in the modified annotation); (ii) region, i.e. canonical CDS, the SCR extension, or negative-control region; and (iii) sample (each ribosome profiling library). Through the procedure outlined above, we obtained matrices of ribosome profiling densities across these axes, which were then subjected to aggregation and noise removal, described hereafter (Supplementary Figure S1). For each transcript and region, consider the Ribo-Seq density matrix *X* of size S × L, where S is the number of samples (libraries) and L is the length of the region in nucleotides; *Xs*, *i* denotes the library-size-normalized coverage at nucleotide position *i* in sample *s*. To increase signal-to-noise, we aggregated data across samples by computing the mean density at each transcript position *i*:

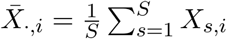

Because averaging can inflate sparse technical artifacts (isolated spikes), we applied a two-stage outlier filtering procedure (Supplementary Figure S1). First, candidate artifact positions were detected using position-wise Z-scores (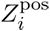) computed on the aggregated mean profile across positions within the same transcript-region:

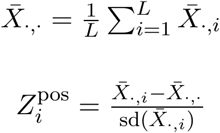

Positions with 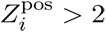 were flagged. Second, for each flagged position *i*, we computed sample-wise Z-scores for each sample:

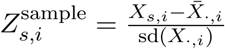

If 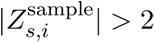 occurred in multiple samples, we treated the peak as a position-specific artifact and removed the entire codon containing that nucleotide. If instead the outlier was present in exactly one sample, that sample was excluded from the calculation of the aggregated density in that transcript-region. After filtering, the final aggregated Ribo-Seq density was calculated for each transcript-region pair as the mean coverage across non-excluded nucleotides and samples (Supplementary Figure S1). This procedure substantially improved the signal quality of Ribo-Seq profiles, as revealed by manual inspection.

#### SCR detection by machine learning

Based on aggregated, noise-removed per-nucleotide Ribo-Seq density vectors, we calculated 32 gene-level metrics and used them as features to train machine learning classifiers (Supplementary Figure S3). They included three coverage metrics for each region (canonical CDS, SCR extension and downstream negative control region): mean Ribo-Seq density (defined as a single value per gene and per region, calculated as the average across all nucleotide positions retained after noise removal; each position itself represents the mean across samples); percentage of nucleotides covered with reads; and Gini index of Ribo-Seq density distribution across nucleotides (a measure of inequality here used to capture non-homogenous artefactual coverage). Another 13 features were included to represent ribosomal profiles in proximity of stop codons: mean Ribo-Seq densities at each of the four codons preceding the canonical stop codon; at each of the four codons preceding the extension stop; and at each of the nine nucleotides preceding the extension stop. Finally, we included three features derived from Gini indices introduced above: ratio between the negative control region and the extension; ratio between the extension and the canonical CDS; and the ratio of these two ratios. No sequence- or length-based features were used for model training.

To avoid missing values and exclude non-informative genes, we applied several filtering steps to obtain a complete matrix of all features described above. We retained only genes with non-zero mean Ribo-Seq density in both the canonical CDS and the SCR extension, with a minimal length of SCR extension of 15 nt (this threshold was applied before noise removal, so that the minimal 3’UTR length is 30 nt).

For the downstream negative-control region, genes with zero coverage resulted in undefined Gini coefficients (NA values). In these cases, we replaced the missing Gini value with the maximum Gini coefficient observed across all genes in the negative-control region. A high Gini coefficient reflects strong inequality in read distribution across the region; thus, assigning the maximum value represents highly non-uniform coverage and a low likelihood that reads originate from active translation. Missing values at the single-nucleotide or single-codon level, which arose during the noise-removal procedure, were replaced with zero.

To develop a computational framework for systematic SCR detection, we first defined labeled gene sets for supervised model training. Our positive reference set representing likely genuine SCR events was compiled based on evolutionary conservation. We retrieved 333 transcripts (from 327 genes) previously predicted to undergo SCR ^23^and expanded this list by querying FlyBase (version FB2023_01 Dmel Release 6.50; QueryBuilder; DataClass: Gene; Field: Gene model comment; Search text: “Stop-codon suppression”), yielding 430 candidate genes. After applying the filtering criteria described above, 366 genes were retained and defined as the “conserved SCR” positive class. As a negative reference set, we defined 447 “ultrastop” genes. These genes naturally terminate with two in-frame stop codons, either consecutive or separated by a single sense codon. For each ultrastop gene, all Ribo-Seq-derived metrics were calculated along a pseudo-extension extending to the third in-frame stop codon. Because these genes are expected to lack genuine ribosome occupancy beyond their two stop codons, they provide an appropriate negative control set for SCR detection. All remaining protein-coding genes passing coverage and 3’UTR filters (n = 6,158) were treated as unlabeled and reserved for downstream prediction.

Three classifier models were trained independently: gradient boosting (GBM), logistic regression (LR), and a single-hidden-layer neural network (NNET) using the caret package from R (version 7.0-1) with default settings ^69^. For performance assessment, labeled genes were randomly partitioned into 80% training and 20% test subsets. Model training and parameter optimization were performed using repeated 10-fold cross-validation (5 repeats) within the training set, using ROC as the primary optimization metric. Logistic regression and neural network models were trained using centered and scaled predictors, whereas GBM models were trained without preprocessing; no hyperparameter optimization was performed. To account for class imbalance between conserved SCR and ultrastop genes, inverse class-frequency weights were applied during training. The optimal GBM model used 100 trees with interaction depth 2. The optimal neural network model used a single hidden unit with decay 0.1.

Final model performance was evaluated on the held-out validation set, resulting in 75-88% validation accuracy (Supplementary Table T2). Feature importance analyses indicated that classification emerged from the combined contribution of multiple partially complementary features rather than from any single dominant predictor, with different classifiers relying on distinct feature-weighting schemes (Supplementary Figure S3A). To assess the impact of missing-value imputation, additional models including binary indicators for imputed Gini values were tested. These indicators were not among the most important features across classifiers (Supplementary Figure S3B).

Afterwards, training was repeated on 100% of the data, and the resulting final models were applied to the 6,158 unlabeled genes to perform genome-wide SCR discovery. Predictions from individual models as well as their intersections were retained for downstream analysis (Supplementary Figure S4A, Figure 1C). To ensure we did not miss well-established candidates, we then ran all conserved SCR genes through the same pipeline and added them to the final set of potential SCR candidates if they were predicted by at least one model. This procedure ensured that all previously recognized conserved SCR genes and novel Ribo-Seq-supported candidates were represented in the final candidate set.

#### *Drosophila* whole genome alignments

We downloaded the “multiz27way” whole genome alignment (WGA) based on *D. melanogaster* dm6, and selected only the 23 entries from *Drosophila* species (https://hgdownload.soe.ucsc.edu/goldenPath/dm6/multiz27way/). Because the WGA annotation was based on a newer genome version, we first performed a liftover from dm3 to dm6 to ensure compatibility with our dataset. The resulting *Drosophila* WGA was used to calculate conservation at positions of interest (e.g. stop codons) and for other downstream applications (e.g. RNA structure prediction with RNAz).

#### Scoring strategy

We scored putative SCR genes based on 15 characteristics: (i-iii) Readthrough prediction by individual Ribo-Seq-based classifiers (GBM, NNET, and LR); (iv) Length of the SCR extension; (v-vi) Proportion of G/C-ending codons in the canonical ORF and in the SCR extension; (vii-viii) Codon usage bias of the canonical ORF and of the SCR extension; (ix-x) Conservation of the canonical stop codon and of the second stop codon (calculated as the mean pairwise nucleotide identity between the *D. melanogaster* stop codon and the aligned codon in other species, ignoring gaps); (xi) Number of reads spanning the stop codon; (xii) Number of in-frame reads spanning the stop codon; (xiii) Nucleotide-level conservation and (xiv) Protein-level conservation of SCR extension; (xv) Ribo-Seq termination peak enrichment at the second stop codon region, calculated as the ratio between the highest ribosome density observed within the four codons immediately upstream of the second stop codon and the mean density of the remaining codons in this region. This metric captures ribosome accumulation associated with translation termination at the second stop codon. For characteristics iv-xv, we used a threshold defined as the 25th percentile value calculated in a strongly supported SCR gene set, represented by conserved SCR genes predicted by all three classifier models. The final score was calculated for each putative SCR gene as the number of Ribo-Seq-based classifiers that predicted SCR plus the number of characteristics iv-xv that exceeded the threshold. Codon usage bias was calculated using the Codon Adaptation Index (CAI), which requires a reference set of highly expressed genes. In this study, the top 25% of genes were selected based on aggregated ribosome profiling density across the canonical ORF. Conservation was calculated across the *Drosophila* WGA described above. For subsequent aggregate analyses, scored genes were grouped as follows: tier 1 (scores 12-15), tier 2 (8-11), tier 3 (4-7), and tier 4 (1-3).

#### Aggregated Ribo-Seq profile analyses

Ribo-Seq densities aggregated across genes were generated from BAM files derived from ribosome profiling experiments. Samples were grouped according to nuclease treatment, and only comparable datasets were aggregated. Transcript annotations were obtained from a GTF file containing a single representative transcript per gene, with three annotated regions: 5’UTR, CDS, and 3’UTR. Genes were included only if the SCR extension exceeded 30 nucleotides in length. For each transcript, per-nucleotide ribosome coverage was extracted from the selected BAM files, corrected by read-length-specific P-site offsets as described above. At each nucleotide position, Ribo-Seq density values were directly averaged across samples. No additional noise filtering was applied. The resulting per-transcript coverage tables were subsequently aggregated across genes within tier groups. Region-specific windows were selected from the nucleotide coverage vectors to construct the final aggregated plots.

#### Analysis of mutations and RNA editing

To identify potential strain-specific, somatic, or phenotypic (e.g. RNA editing) mutations at stop codons, we performed variant calling from aligned sequencing reads from ribosome profiling samples, restricting the analysis to genomic coordinates of canonical CDS stop codons. We performed variant calling using bcftools (version 1.18) ^70^. First, we used bcftools mpileup to summarize all sequencing reads aligned to each stop codon, using the *D. melanogaster* reference genome dmel r5.57 and setting the maximum read depth to 10,000,000. Next, variants were identified using bcftools call, in multiallelic calling mode (option -mv) to detect single-nucleotide variants and short insertions or deletions. Only variants with a Phred-scaled quality score greater than 20 (corresponding to an estimated error probability below 1%) were retained for downstream analysis. We identified potential stop codon alterations in 26 genes within our Ribo-Seq SCR dataset. Out of those, 18 carried mutations replacing one stop codon with another stop. In contrast, 8 genes (*Ebp, CG13403, CG13403, CG13101, CG6321, CG13048, Naa80, CG14511, stum*) exhibited mutations converting the annotated stop into a sense codon. Because such substitutions artificially mimic readthrough in Ribo-Seq interrogation, these genes were excluded from further analysis.

#### Identification of conserved RNA secondary structures

We predicted conserved RNA secondary structures using RNAz (version 2.1.1) ^40^. The input data was derived from the *Drosophila* WGA, extracting regions corresponding to transcripts of *D. melanogaster*, typically joining multiple exons. These WGA-derived transcript alignments were scanned using sliding windows of 120 nt with a step of 20 nucleotides, to identify regions with evidence of structural conservation. Only windows meeting the following criteria were retained: maximum gap fraction ≤ 0.7, maximum masked fraction ≤ 0.1, minimum number of sequences ≥ 2, minimum pairwise sequence identity ≥ 50%, maximum number of sequences ≤ 23, minimum alignment length ≥ 120 nt. To reduce sampling bias in large alignments, 7 sequences were randomly selected three times from each window using rnazSelectSeqs.pl (--num-seqs=7 --num-samples=3), and only the highest-probability hit per window was retained.. Each alignment window was scored with RNAz, which predicts conserved and thermodynamically stable RNA structures based on minimum free energy (MFE), a stability z-score, and a structure conservation index, combined via a support vector machine classifier. Windows with probability >0.8 were considered for downstream analyses. These underwent transcript-to-genomic coordinate conversion to map their positions back onto the reference genome.

#### Analysis of protein localization

Protein sequences were generated for both the canonical CDS and the canonical CDS extended by the SCR region. Localization analysis was performed using deeploc2 (version 2.1) ^71^ with the “accurate” model setting. For each sequence, DeepLoc returned class probabilities for the following compartments: Cytoplasm, Nucleus, Extracellular, Cell membrane, Mitochondrion, Endoplasmic reticulum, Lysosome/Vacuole, Golgi apparatus, and Peroxisome. To identify SCR-dependent localization changes, we required a shift in the top predicted compartment between canonical and extended proteins with an absolute probability difference ≥30%. Moreover, we required the localization probability to be ≥70% for at least one protein isoform, ensuring high-confidence predictions. Information about genes with detected change in localisation is provided in Supplementary Table T6.

#### Intrinsic disorder analysis

This analysis was performed using AIUPred tool (version v0.9) ^72^. For each residue, the tool assigns a disorder score ranging from 0 to 1, with higher values indicating a greater probability of intrinsic disorder. For each gene, we calculated the median disorder score separately for the canonical CDS and the SCR extension by aggregating residue-level scores across each respective region.

#### Sequence logo

To examine nucleotide preferences around stop codons, fixed-length sequence windows centered on the stop codon were extracted from each gene and aligned relative to the first nucleotide of the stop codon. Sequence logos were generated using ggseqlogo (version 0.2) in R.

#### Design of library of candidate SCR stimulators

The candidate library was designed from an extended set of 3,284 SCR genes obtained using the first batch of results from a machine learning approach analogous to that described above. This gene list largely overlaps with the final set presented above, obtained after model refinement. The library consisted of 129-nt variable inserts and it was designed to prioritize conserved RNA structures. Specifically, the final library consists of four groups of sequences: (i) “CDS-3UTR” conserved structures, i.e. one RNAz prediction overlapping the stop codon per gene (highest RNAz probability score) in 223 genes; (ii) “CDS” conserved structures upstream of the stop (at most 2kb away) in top SCR genes, defined as with SCR score > 10 or predicted by conservation, i.e. one or multiple structures in 247 genes; (iii) “3UTR” conserved structures downstream of the stop (at most 2kb away) in top SCR genes, i.e. one or multiple structures in 273 genes; (iv) “around-stop” sequence windows centered on the stop codon in top SCR genes, one each for 670 genes. Conserved structures were frame-adjusted and extended to 129 nt. Symmetrical extension was applied when possible; otherwise, extensions were constrained to avoid additional in-frame stops. CDS-UTR sequences were truncated before downstream stops and extended upstream. CDS sequences terminated at the most at the stop codon and were extended upstream to reach the desired length, with the final 15 nt reserved for the native stop and its ±6 nt context. The same rules were applied to 3UTR sequences, with upstream-downstream inversion.

This procedure yielded 1,592 “base” elements from *D. melanogaster*, each represented by multiple variants (full information in Supplementary Table T9). For each element, two stop codon variants were included to cover all three stops. Then, evolutionary replicates were added from orthologous sequences in other *Drosophila* species using WGAs (four species for CDS-UTR, three for CDS and UTR elements). Mapping was performed relative to the stop codon for CDS-UTR and around-stop elements, and relative to the 3’ structure boundary for CDS and UTR elements. For structured candidates, species were selected iteratively to maximize structural similarity while minimizing sequence similarity. For around-stop candidates, species were selected iteratively to match the mean and standard deviation of sequence identity obtained in the CDS-UTR class. Duplicates were removed. Additional in-frame stops were recoded using two mutation schemes (i: TAA→TTA, TAG→TAT, TGA→TTA; ii: TAA→TAT, TAG→TGG, TGA→TGG), and cloning restriction sites (EcoRV, NotI) were altered when present.

#### Identification of SCR candidates from pooled screening

FASTQ files from paired-end sequencing of pooled screening experiments were processed using DiMSum (v1.3) ^73^ to obtain read counts for each insert in each cell fraction. Paired-end reads were first merged to reconstruct full insert sequences. Reads were then quality-filtered, and inserts were identified by exact sequence matching to a predefined list of expected inserts using a barcode-insert mapping file. The resulting count table was analyzed to identify functional SCR stimulators. Each sorting replicate was analyzed independently. Inserts with low representation were filtered out prior to analysis by excluding entries whose total counts across the four fluorescence bins fell below the 10th percentile, yielding 1,095 elements corresponding to 683 genes. For each candidate stimulator, we conducted all pairwise comparisons between cell fractions (i.e. ultra-low + low combined, medium, and high). Counts were modeled using a negative binomial generalized linear model, with cell fraction as the predictor of interest and species included as a covariate. Cell fraction p-values were adjusted using Benjamini-Hochberg FDR. For each candidate stimulator, sequences from *D. melanogaster* and two additional species were included. Additional sequence variants (i.e., constructs with alternative stop codons, orthologs from other species, or different mutational patterns) were excluded to ensure equal representation across stimulators. Stimulators lacking sufficient evolutionary replicates after filtering were excluded from further analysis. The final list of selected candidates (Supplementary Table T7) were those with FDR<0.1 for both sorting replicates. The results of this analysis without applying the threshold are presented in Supplementary Table T9.

#### Single sample SCR calling and clustering

We selected 1,374 genes from the machine learning-derived Ribo-Seq candidate set (score ≥8; tiers 1 and 2), computed per-sample Ribo-Seq features, and implemented a selection procedure to detect sample-specific SCR events, following the approach described in Dunn et al ^24^. Read counts were obtained using featureCounts (v. 2.0.1) ^74^ with default parameters, quantifying three regions per gene: canonical ORF, SCR extension, and negative control region. Gene counts were converted to FPKM (Fragments Per Kilobase of transcript per Million mapped reads) values. The SCR rate was calculated as the ratio between the SCR extension and canonical ORF FPKMs; the drop-off metric was defined as the analogous ratio between negative control region and SCR extension. In each case, a pseudocount of 0.01 was added to the denominator. To characterize the distribution of ribosome footprints across the SCR extension, we calculated nucleotide-level coverage statistics using Plastid, and derived the following features: percentage of SCR extension nucleotides covered by any read; and positions of the first and last detected reads along the normalized SCR extension length.

Next, we set to choose feature thresholds for sample-specific SCR detection. To define them, we assembled a reference SCR gene set per-sample, consisting of conserved SCR genes with SCR extension FPKM ≥ 0.1, SCR rate < 2, the first detected read located within the first quarter of the SCR extension, and the last detected read located within the last quarter. Our exploratory analyses revealed that SCR Ribo-Seq features in the reference set co-varied with overall expression level, motivating us to define expression-dependent thresholds (Supplementary Figure S41). Thus, we modelled the relationship between SCR extension FPKM and each Ribo-Seq feature separately, fitting generalized additive models with aggregate metrics of reference SCR genes. We fitted two models (one for lower and one for upper bounds) for SCR rate, and one lower bound model for the other features (SCR extension coverage and drop-off). Each model was fitted with ten data points, one for each expression decile. For lower bound models, these consisted of the 5th percentiles of feature values of reference genes in that decile; for upper bound models, we used the analogous 95th percentiles. Models were then applied to calculate the lower and/or upper thresholds of each metric for every candidate gene in each sample, matching its observed expression level. To increase robustness, we selected only genes predicted to undergo SCR in two or more samples (881 genes), which were used for clustering and heatmap visualization, as follows.

When sample-specific SCR was not detected, genes were classified according to whether their expression level was sufficient to permit SCR detection. Based on exploratory analyses (Supplementary Figure S34), we defined a minimum canonical CDS expression threshold of 10 FPKM. Thus, we constructed a discretized gene-sample matrix representing gene states: 1 (zero or low expression), 2 (expressed), and 10 (SCR). Genes and samples were then clustered using hierarchical clustering (Ward.D2) with Euclidean distance, and genes were partitioned into eight clusters. To evaluate clustering robustness, we introduced controlled label noise into the matrix by randomly perturbing a fraction of entries using constrained transitions (1→2, 10→2, 2→1 or 2→10), reflecting plausible misclassification between expression and SCR states. Noisy matrices were reclustered, and cluster assignments were compared with the original clustering. Cluster stability was quantified by calculating, for each original cluster, the fraction of its genes recovered in the best-matching cluster of the noisy solution. This analysis was repeated across multiple noise levels and replicates to assess how strongly clusters were affected by increasing annotation noise (Supplementary Figure S42).

To analyze expression changes associated to SCR, we selected genes with at least 6 SCR+ and 6 SCR- samples and matched RNA-Seq and Ribo-Seq available. For each gene, expression values were first transformed as log2(FPKM + 0.1), averaged separately across SCR+ and SCR- samples (i.e., with gene-specific sets of samples considered), and the log2 fold change was calculated as the difference between these means. Results were visualized on the full gene set or grouping by gene cluster (Supplementary Figure S36). The statistical significance of the difference between Ribo-Seq and RNA-Seq log2 fold changes was assessed using paired Wilcoxon tests.

#### Gene Ontology enrichment analysis

Gene set overrepresentation analysis was performed per gene cluster using the enrichGO function from the clusterProfiler package (version 4.16.0) ^75^. The background gene set consisted of all genes meeting the following criteria: SCR extension length greater than 15 nt, and at least one read detected in the SCR extension in at least one sample (8,338 genes in total). Both the p-value and q-value cutoffs were set to 0.05. Redundant GO categories with identical gene sets were then removed, and the top 6 categories for each cluster were selected based on q-value. From these, the top 3 categories were manually chosen for display in the heatmap annotation.

Gene Ontology enrichment analysis for high-SCR genes (tiers 1 and 2) was performed using the same background gene set, enrichment function, and significance thresholds (p-value and adjusted p-value). Redundant terms were reduced using the simplify function (cutoff = 0.65, based on adjusted p-values), followed by computation of pairwise term similarity and visualization with emapplot (top 30 categories).

#### Proteomics Analysis

The proteome database for MS/MS (tandem mass spectrometry) data interrogation was constructed as follows. One isoform per gene was selected as described in the gene annotation section, and the nucleotide sequences corresponding to the canonical CDS plus the SCR extension were obtained. These were translated into 20 alternative protein sequences per gene, replacing the stop codon with each of the 20 amino acids. The index of the stop-codon-derived amino acid was retained for each gene for later interpretation of peptide mapping. Decoy sequences were generated by reversing the amino acid sequences and used as negative controls.

Peptide identification was performed using deTELpy (v. 0.1.13) ^76^, which builds upon the FragPipe pipeline (which employs using MSFragger engine version 4.0 ^77,78^). Using the stop codon index, peptides were classified as canonical or SCR peptides, the latter potentially spanning the stop codon. False discovery rate (FDR) control was applied using a target-decoy strategy, first independently for each dataset (1% FDR threshold) and subsequently after aggregation across all datasets. Publicly available MS/MS proteomics datasets were obtained from the PRIDE database https://www.ebi.ac.uk/pride/, then filtered to retain only those with *D. melanogaster* as the sole source organism and generated by mass spectrometry instruments compatible with deTELpy. In total, 123 datasets were included (Supplementary Table T8). Only candidates with predicted SCR extensions of ≥4 amino acids were considered, as shorter sequences are more prone to ambiguous peptide assignment.

#### Drosophila Stop Codon Readthrough Atlas

The Webserver was built using RShiny v.1.11.1, hosted in our laboratory server. It internally uses precomputed data tables generated as described in this manuscript.

#### AlphaFold2 Structure Prediction

The amino acid sequence corresponding to the canonical protein and its predicted translational readthrough extension was modeled using AlphaFold2 (v2.3.2). Structure prediction was performed with the monomer model preset using the full AlphaFold2 database suite. Multiple sequence alignments were generated with HHblits and HHsearch, and structure inference was subsequently carried out using the precomputed alignments on a GPU-enabled computing node. The highest-ranked AlphaFold2 model was selected for further structural analyses.

#### PhyloCSF analysis

PhyloCSF ^18^ was run using the 12-flies dm3 alignment using the default “mle” strategy. We computed PhyloCSF scores for the regions starting 0, 1, or 2 nucleotides downstream of the annotated stop codon up to but not including the next stop codon in that frame, excluding regions with 0-length, those with any degenerate bases (N’s) in the assembly, and those in the mitochondrial chromosome. If the annotated transcript ended before the second stop codon was reached, we extended the transcript past the end without splicing. We selected transcripts with length of extension >= 20 codons. PhyloCSF-ΨEmp scores were computed as reported previously ^23^.

#### Data visualization

Figures were generated in R using ggplot2, with additional plot assembly and customization performed using the cowplot and ComplexHeatmap packages where appropriate, in some cases refining layout and appearance with Adobe Illustrator.

### EXPERIMENTAL MODEL DETAILS

#### Cell culturing

*Drosophila* S2, S2R+ NPT005, 24B>Ras attP-L5-CloneB8 (24B5-B8; embryo cells at late embryonic stage with characteristics of muscle tissue) and ML-DmBG3-c2-attP-25C6 (BG3; cells from the central nervous system) cells were obtained from the Drosophila Genomics Resource Center. S2 and S2R+ cells were cultured at 25 °C in Schneider’s medium (S0146-100ML, VWR or L0207-500, Sigma) supplemented with 10% fetal bovine serum (FCS; Hyclone SH30070.03). BG3 and 24B5-B8 cells were grown in Shields and Sang M3 Insect Medium (S3652-500ML, Sigma) supplemented with 10% FCS. BG3 cells were supplemented with 10μM insulin. Insulin powder (Insulin from bovine pancreas, I5500-50MG, Sigma) was diluted to 10mg/mL in acidic water (1µL HCl 32% per 1mL MiliQ) and stored at -80°C in 100μL aliquotes as a stock solution. Insulin supplemented media was not stored for more than 10 days due to insulin degradation. All insect cell lines were cultured without antibiotics, in sealed flasks without CO_2_ control under non-humidified conditions. S2 and S2R+ NPT005 cells were passaged every five days by gentle resuspension through pipetting, due to their low attachment. BG3 and 24B5-B8 cell lines were passaged every four days with trypsin (Trypsin-EDTA (0.05%), phenol red; Gibco, 25300054).

Mouse myoblast C2C12 cells and human HEK293 cells were cultured in high-glucose Gibco Dulbecco’s Modified Eagle Medium (Thermo, 61965026), supplemented with 10% FCS (Hyclone, SH30070.03) or 10% FBS (Thermo, A5256701) and Gibco Penicillin-Streptomycin 10.000 U/ml antibiotic mixture (Thermo, 15140-122). Cell confluence was maintained below 80%. Cells were split 1:5 every 48h, or 1:10 every 72h. All experiments on *Drosophila* and mammalian cells were conducted at passage <20 after thawing.

### EXPERIMENTAL METHODS DETAILS

#### Plasmid transfection

Transfections in *Drosophila* cells were performed using Effectene (301425, Qiagen). The transfection mix consisted of 520 ng DNA in 130 µL EC buffer, 4.13 µL Enhancer, and 13 µL Effectene. Flow cytometry, Western blotting, and luciferase assays were performed 72h after transfection since early experiments determined that the highest signal was detected at this time point. For flow cytometry, cells were seeded at 3 × 10^6^ cells/mL (2 mL in total) in 12.5 cm^2^ flasks two hours before transfection. At the time of cytometry, cells were collected by pipetting and resuspended in FACS buffer (1× PBS supplemented with 5% FBS, filtered). For luciferase assays, S2 and S2R+ cells were seeded in 24-well plates (500 µL per well, 1.8 × 10^6^ cells/mL) and transfected using the same Effectene protocol as for 12.5 cm^2^ flask but used for 3 replicates (45ul of mix per each well). For BG3 and 24B5-B8 cell lines, 500,000 cells per well were seeded in 24-well plates with their respective culture media 24h before transfection; the media was changed to avoid toxicity of the Effectene reagents 24h after transfection, and luciferase assays were performed 96h after transfection.

For studies in HEK293 cells, 70,000 cells per well were seeded in 48-well plates. Cells were transfected with Lipofectamine 3000 24 h after seeding, and luciferase assays were performed 24 h later. Lysates were diluted 20-fold in 1X PLB to avoid luminometer signal saturation. For experiments in C2C12 cells, 9,000 cells per well were seeded in 48-well plates. Prior to transfection, complete media was changed to antibiotic-free. Transfection was performed using the jetOPTIMUS reagent (Sartorius, 101000051). To make the transfection mix for one well, 250ng of DNA was suspended in 25 µL of DMEM and vortexed. 0,375 µL of the jetOPTIMUS reagent was added to the mix, and after vortexing and 10 minutes of incubation, the complete transfection mix was added dropwise to the wells. Four hours after transfection, antibiotic-free media was changed to complete media.

#### Plasmid constructs

All plasmid and library synthesis and cloning were performed by GenScript (Piscataway, NJ, USA) starting from a pcDNA3 vector backbone. The primary construct used for library integration and screening contained the following elements (Supplementary Figure S20): attB - Acti5C promoter - HMOX1 - P2A - PuroR - SV40 polyA - OpIE2 promoter - chimeric intron - emiRFP703 - T2A’ (i.e. T2A with synonymous mutations introduced at every third nucleotide) - EcoRV site - stop containing insert- NotI site - T2A - eGFP - BgH poly(A) - attB. The insert library was synthesized in pooled format and cloned into the plasmid backbone using the NotI and EcoRV restriction sites. For validation experiments, we performed transient transfection assays using a dual-luciferase reporter construct based on the same backbone (Supplementary Figure S24) containing: attB - CMV promoter - insect HSP70 promoter - chimeric intron - Renilla luciferase - T2A’ - Acc65I site - stop containing insert - BamHI site - T2A - Firefly luciferase - BgH poly(A) - attB. This luciferase reporter plasmid contained two promoters, CMV for mammalian expression and Hsp70 for insect expression. Prior to these experiments, AUG codons within the Hsp70 promoter region were mutated to minimize alternative initiation, and promoter function was confirmed.

#### Integration-based screening

S2R+ NPT005 cells (which carry a AttP-flanked mCherry cassette) were transfected with the screening plasmid library together with the φC31 integrase-encoding plasmid pBS130 (#26290) at a 1:1 ratio using Effectene, following the manufacturer’s instructions. Specifically, two hours before transfection, cells were seeded in two 75 cm^2^ flasks at a concentration of 3 × 10^6^ cells/mL, using 12 ml per flask. In total, 7.2 × 10^7^ cells were transfected using 1,560 ng of library plasmid and 1,560 ng of pBS130 in 780 µL EC buffer, supplemented with 25 µL Enhancer and 78 µL Effectene. In parallel, two control transfections were performed: one with the library alone to monitor dilution of non-integrated plasmids over time, and one with pBS130 plus the library backbone with the frameshifted stop context of the *tj* gene, as positive control for integration. Medium was changed the following day. One week after transfection, medium was replaced and puromycin was added to a final concentration of 5 µg/mL to samples containing pBS130. After one additional week, cells were diluted to 3 × 10^6^ cells/mL and maintained under puromycin selection for approximately 1.5 month (∼7 passages). Passaging was continued until cells transfected with the library alone (without integrase) became completely iRFP-negative, indicating loss of the non-integrated plasmid. A total of 1.3 × 10^6^ unsorted integrated cells were collected. Integration efficiency was approximately 70%. Puromycin-containing medium was replaced with regular growth medium one week prior to cell sorting. Cells were detached by pipetting in culture medium, centrifuged at 300 × g, and resuspended in FACS buffer (1× PBS supplemented with 5% FBS). Sorting was performed on an BD FACSDiscover™ S8 Cell Sorter (gating strategy in Supplementary Figure S43). Untransfected S2 cells were used to assess autofluorescence. S2R+ NPT005 cells served as a control for mCherry, S2 cells transfected with the OpIE2-GFP plasmid (Addgene #173505) as a GFP only control, and S2 cells transfected with iRFP-frameshifted_tj-GFP as an iRFP only control. These controls were used for fluorescence unmixing, allowing discrimination of low SCR-dependent signals.

Following a first round of cytometry analysis, a binning strategy for cell sorting was adopted in which four gates were defined along the GFP intensity axis, corresponding to ultra-low, low, medium, and high GFP levels. Cells were sorted into four populations based on GFP intensity using a 100 µm nozzle at 4 °C into 5 mL tubes containing 500 µL of Schneider’s medium. Due to differences in cell abundance across gates, different numbers of cells were collected per bin; however, these numbers were kept constant between biological replicates (see Supplementary Table T10). At minimum, 3 × 10^5^ cells were collected for the ultra-low bin, and up to 2.5 × 10^6^ cells for the highest-intensity bin. Cells were then centrifuged, washed with PBS, after that pellets were frozen at -20 °C prior to DNA extraction. Sorting was performed in two independent rounds to generate two biological replicates, each comprising ∼2 * 10^8^ cells with integrated plasmid.

Next, genomic DNA was extracted using the Quick DNA miniprep kit (Zymo D3024), using one column per each sorted sample and 6 columns for non-sorted cells. DNA was eluted in 50 µL in each column. All DNA from sorted samples was used for PCR amplification in 2* 50 µL reactions using primers binding to regions surrounding the variable insert (Supplementary Table T11). We used 25 ul of genomic DNA (with concentration from 5 ng to 15 ng/µl) in each 50µl reaction. PCR conditions were:98 °C 1 min (initial denaturation); 23 cycles of 98 °C 15 s (denaturation), 68 °C 20 s (annealing), 72 °C 20 s (elongation); 72 °C 7 min (final elongation). The forward primer annealed to the first T2A’ sequence, while the reverse primer annealed to the second T2A. Both primers contained a stretch of 1-7 random nucleotides (N2-N7) to increase sequence complexity during sequencing; equal amounts of primers containing N2 to N7 were mixed prior to PCR amplification. PCR products were purified using a Qiagen kit (#28704) and eluted in 50 µL MQ water. DNA concentration was measured using a NanoDrop spectrophotometer. 50 ng of each purified PCR product were used in a second PCR to attach Illumina adapters. PCR conditions were: 98 °C 30 s (initial denaturation); 5 cycles of 98 °C 10 s (denaturation), 60 °C 20 s (annealing), 72 °C 20 s (elongation); 72 °C 7 min (final elongation). PCR2 products were purified using AMPure beads (0.9× ratio) and sequenced on a NextSeq 500 using a 300-cycle mid-output kit in paired-end mode.

#### Luciferase-based validation assays

Selected candidate elements were cloned into a dual-luciferase reporter vector (see Plasmid constructs) and assayed using the Dual-Luciferase Reporter Assay System (Promega E1910). After transfection (see Plasmid transfections), cells were collected by pipetting, washed with PBS, and lysed in 150 µL Passive Lysis Buffer (PLB) for 15 min at room temperature. Lysates were centrifuged for 5 min at 13,000 × g, and supernatants were collected. Two microliters of lysate were assayed using 65 µL of LAR II and Stop & Glo reagents. Higher lysate amounts caused signal saturation. For C2C12 only, the PLB incubation time was increased to 30 min, and 20 µL of lysate were assayed using 80 µL of LAR II and Stop & Glo reagents. Luminescence was measured using a Berthold Lumat LB 9507 luminometer in single-tube mode. Raw Fluc and Rluc values are presented in Supplementary Table T12. Rluc counts were deemed non-reliable when below 90,000 RLU for C2C12 and 300,000 for all others, owing to differences in transfection efficiency.

#### Western blot

Seventy-two hours after transfection, S2 cells grown in 12.5 cm flasks (one flask per sample) were collected by pipetting and centrifugation at 900 × g for 5 min. Cell pellets were washed once with 1× PBS and lysed in lysis buffer (20 mM Tris-HCl pH 7.5, 150 mM NaCl, 1% NP-40 (Merck, 492016), supplemented with 1× protease inhibitor cocktail (Sigma, 11836153001)). Lysates were incubated at 4 °C for 30 min and subsequently clarified by centrifugation. The supernatant was collected as total protein extract. Protein concentration was determined using the Pierce Bradford assay (Thermo Fisher, 23227), and 5 µg of total protein per sample was resolved by SDS-PAGE and transferred to a nitrocellulose membrane (Thermo Fisher, LC2000). Membranes were probed with a primary antibody against GFP (rabbit; GenScript, A01388), followed by an HRP-conjugated secondary antibody (anti-rabbit IgG; GenScript, A01827). Detection was performed using enhanced chemiluminescence (ECL).

## Supporting information

Excel file E1. Supplementary Tables T1-T12 as separate sheets

## Acknowledgments

We thank Raghuvir Viswanatha, Stephanie Mohr, Arthur Luhur, Ilya Vorontsov, Gary Loughran, Montserrat Corominas, Jordi Bernués, Ignasi Toledano, Pavel V. Baranov, Antonio Torres-Mendez and Lucia Prieto Godino for valuable advice, insightful discussions, and technical support. We also thank the Flow Cytometry Facility at the Scientific and Technological Centers of the Universitat de Barcelona (CCiTUB) for assistance with sorting experiments, and the Boehringer Ingelheim Fonds for providing a travel grant to attend the EMBL course “The Fundamentals of High-End Cell Sorting.”

## Funding

MM is funded by grants RYC2019-027746-I, PID2020-115122GA-I00, and PID2023-147164NB-I00, funded by MICIU/AEI /https://doi.org/10.13039/501100011033 and by “ESF Investing in your future”, FEDER, UE; and received support from Comissió Interdepartamental de Recerca i Innovació Tecnològica (2021SGR00279). NM received support from predoctoral fellowship FPU2021-03659, from an EMBO Scientific Exchange Grant supporting an internship at the Max Planck Institute for Molecular Cell Biology and Genetics (MPI-CBG) in Dresden, and from a EuroLife PhD & PostDoc Mobility and Knowledge Exchange Grant supporting an internship at the University of Strasbourg. IJ was supported by the National Human Genome Research Institute of the National Institutes of Health under Award Number U24HG007234. The content is solely the responsibility of the authors and does not necessarily represent the official views of the National Institutes of Health. AL was supported by a AFM-Téléthon grant (TG2022 #24381).

## Declaration of interests

The authors declare no competing interests

## Declaration of generative AI and AI-assisted technologies in the writing process

During the preparation of this work, the authors used OpenAI ChatGPT in order to improve conciseness and readability of the manuscript. After using this tool, the authors reviewed and edited the content as needed and take full responsibility for the content of the published article.

## Supplemental information titles and legends

Document S1. Supplementary Figures S1-S43.

Excel file E1. Supplementary Tables T1-T12 as separate sheets. All table captions are included in the first sheet.

## Notes

### Competing Interest Statement

The authors have declared no competing interest.

