## Supplementary material for "Regulatory landscape of widespread stop codon readthrough in *Drosophila*": Document S1: Supplementary Figures S1-S43.

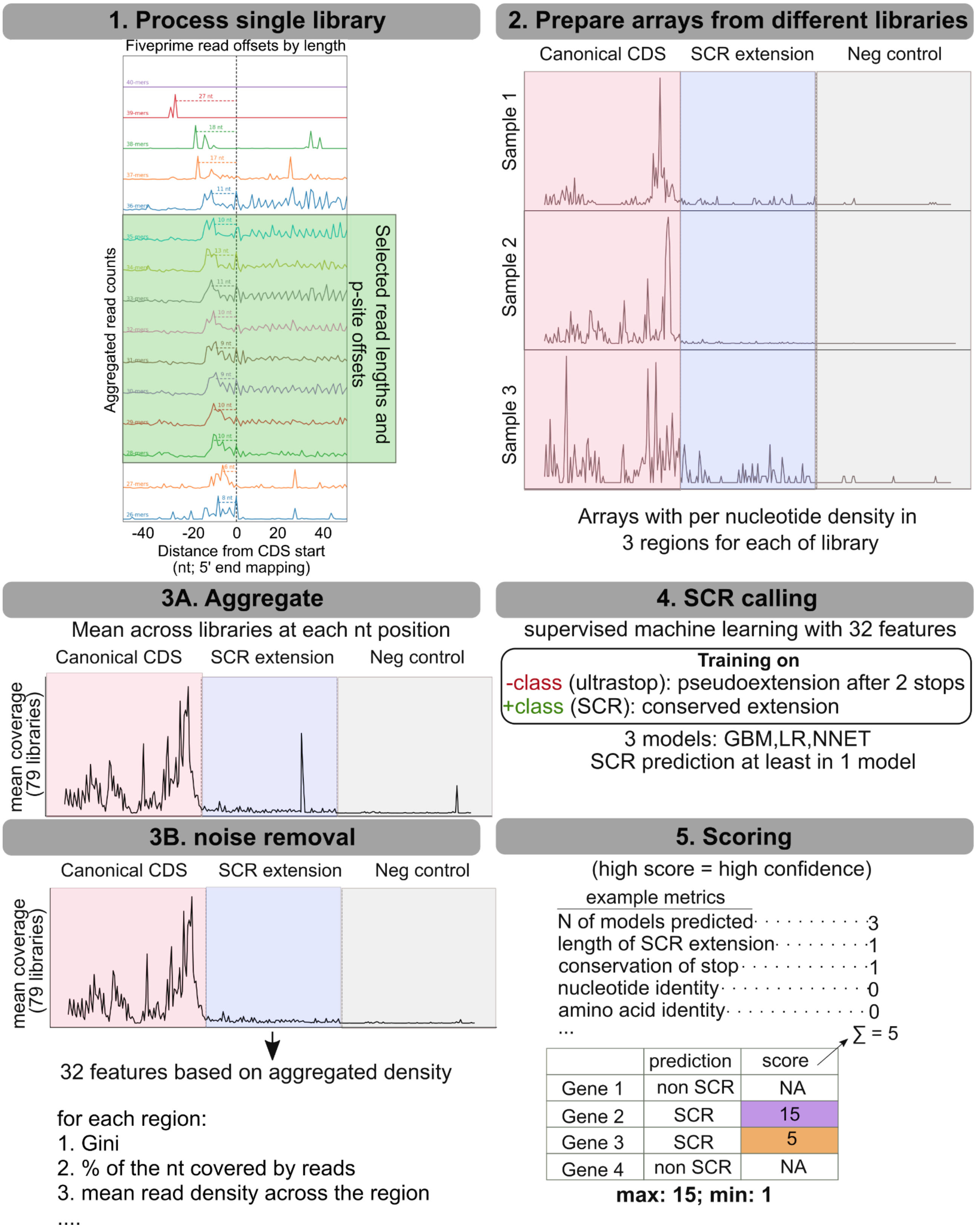


**Supplementary Figure S1.** Aggregation-based Ribo-Seq pipeline for SCR detection.

(1) Processing of individual libraries. For each Ribo-Seq sample, P-site offsets were determined manually and separately for each read length, and footprint lengths exhibiting clear 3-nt periodicity and a pronounced start-codon peak (green rectangle) were selected.

(2) Per-transcript Ribo-Seq density quantification. For each gene, nucleotide-resolution Ribo-Seq density was computed independently for each sample across three regions: the canonical CDS, the SCR extension, and a downstream negative control region.
(3) A. Cross-sample aggregation. Per-nucleotide densities were aggregated by computing the mean density at each nucleotide position, generating position-resolved profiles for each gene and region.
(3) B. Noise removal. To prevent amplification of technical artifacts during aggregation, positions exhibiting abnormally high density relative to the regional background were identified and excluded, removing spurious peaks from the aggregated signal.
(4) SCR prediction. The resulting profiles were used to compute multiple features capturing ribosome occupancy patterns, which were integrated using supervised machine learning models to predict SCR candidates.
(5). Scoring of SCR candidates. Predicted SCR genes were further evaluated using a scoring scheme that integrates model predictions with additional independent features, including Ribo-Seq-derived metrics, sequence properties, and evolutionary conservation. This approach assigns a cumulative score to each gene, enabling stratification into tiers.

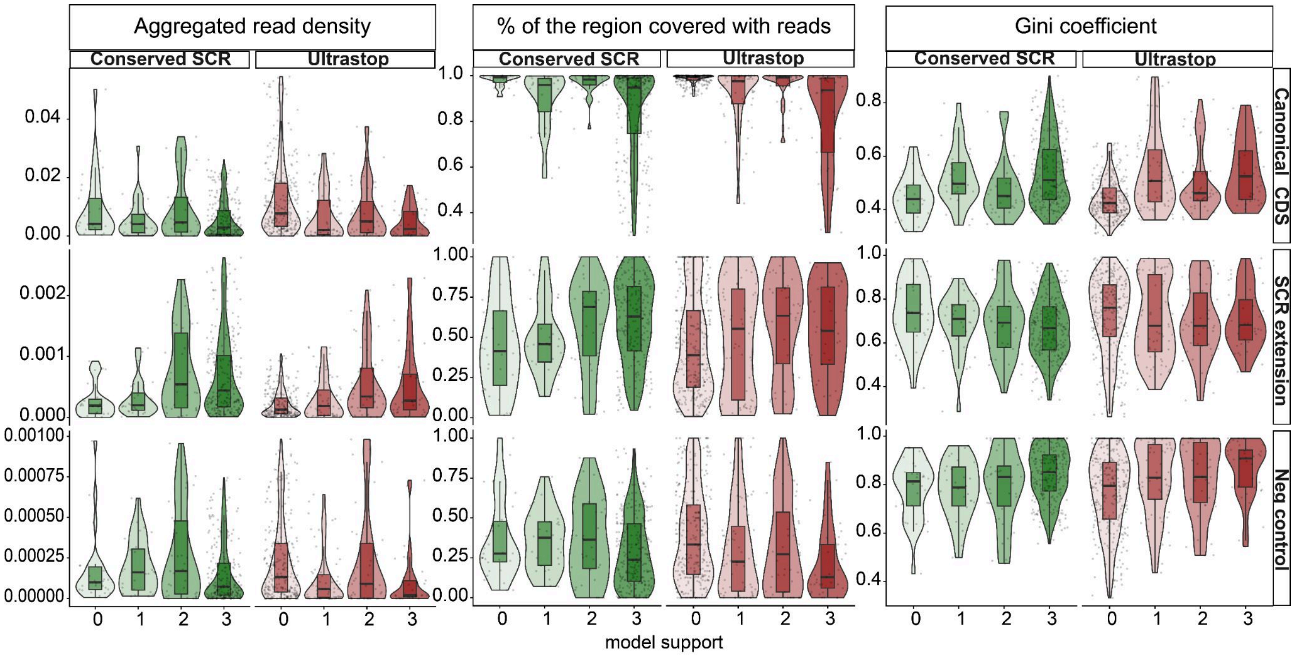


**Supplementary Figure S2.** Comparison of ribosome profiling-derived features across genes supported by different numbers of machine learning models. Violin plots show the distribution of aggregated read density, percentage of the region covered with reads, and uniformity of read distribution (defined by the Gini coefficient) for canonical CDSs, SCR extensions, and negative-control downstream regions. Genes are grouped according to the number of machine learning models supporting stop codon readthrough prediction (0–3 models). Conserved SCR genes (green), which were used as the positive training set, and ultrastop genes (red), which were used as the negative training set, are shown separately. Notably, conserved SCR genes predicted by 0 or 1 model exhibited almost no detectable ribosome density within SCR extensions. This likely reflects context-specific SCR activity absent from the analyzed Ribo-Seq datasets, resulting in weak experimental support despite evolutionary evidence of readthrough.

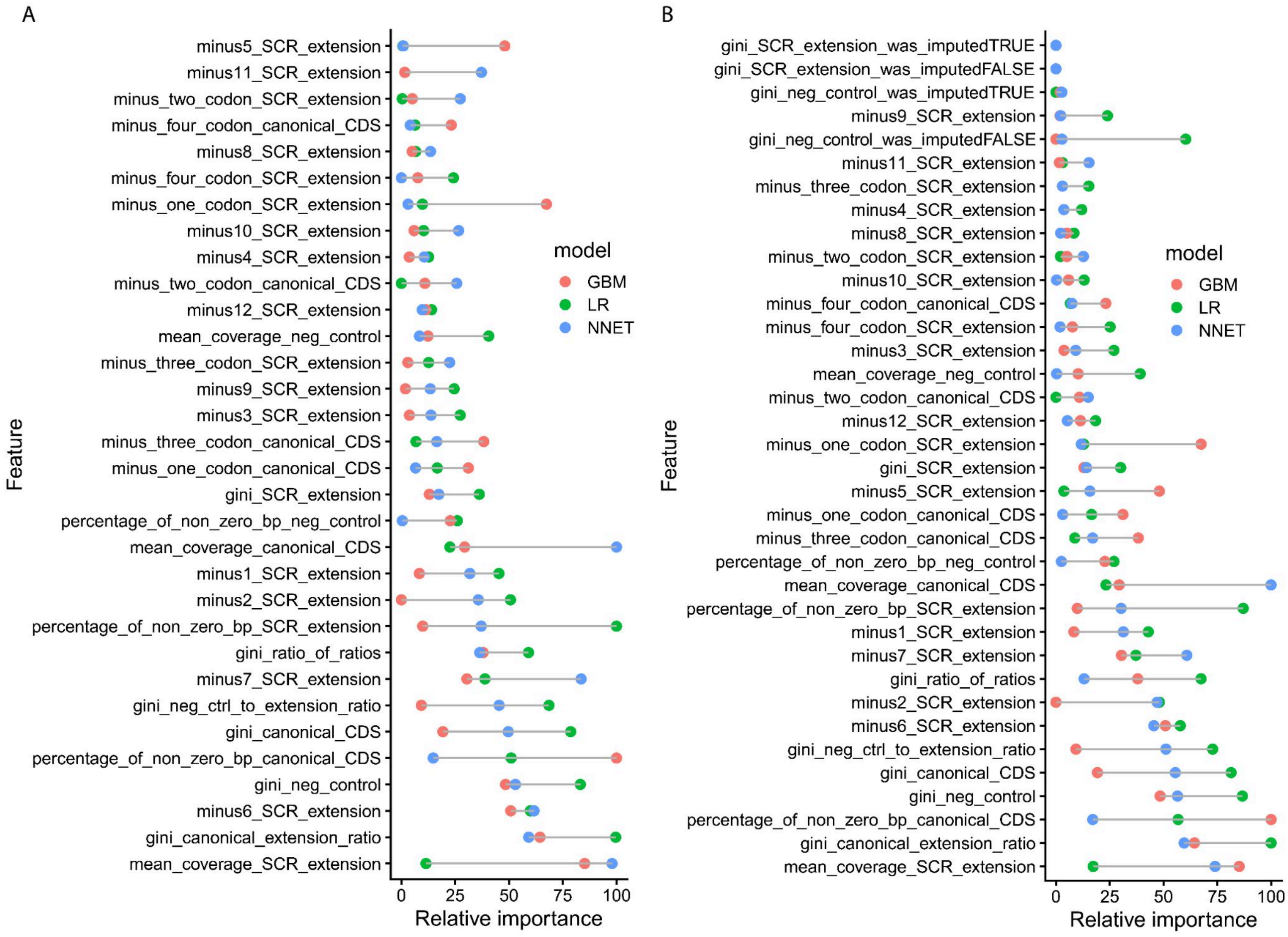


**Supplementary Figure S3.**

**(A)** List of 32 training features used in the machine-learning SCR classifier (Gradient Boosting Machine (GBM), Logistic Regression (LR), and Neural Network (NNET)), sorted by their maximum mean importance across models. This ranking reflects both the overall contribution and consistency of each feature across classifiers. Feature names include a suffix indicating the region used for calculation (canonical CDS, SCR extension, or negative control - region downstream of the second stop codon that is either 114 nt or equal to the length of SCR extension if it is smaller than 114).

**(B).** Imputation of missing values had minimal impact on ML predictions. In addition to the 32 primary features, binary indicators specifying whether Gini coefficients were imputed were included in the extended 36-feature set used for model training. Gini values for SCR extensions were imputed when the SCR extension remaining after noise correction was extremely short (3-9 nt; 8 genes total, of which 1 was predicted to undergo SCR). Gini values for the negative-control region were imputed when the region downstream of the second stop codon contained no reads (788 genes total, of which 389 were predicted by at least one model). The full list of 36 training features, including imputation indicators, is shown sorted by maximum mean importance across models.

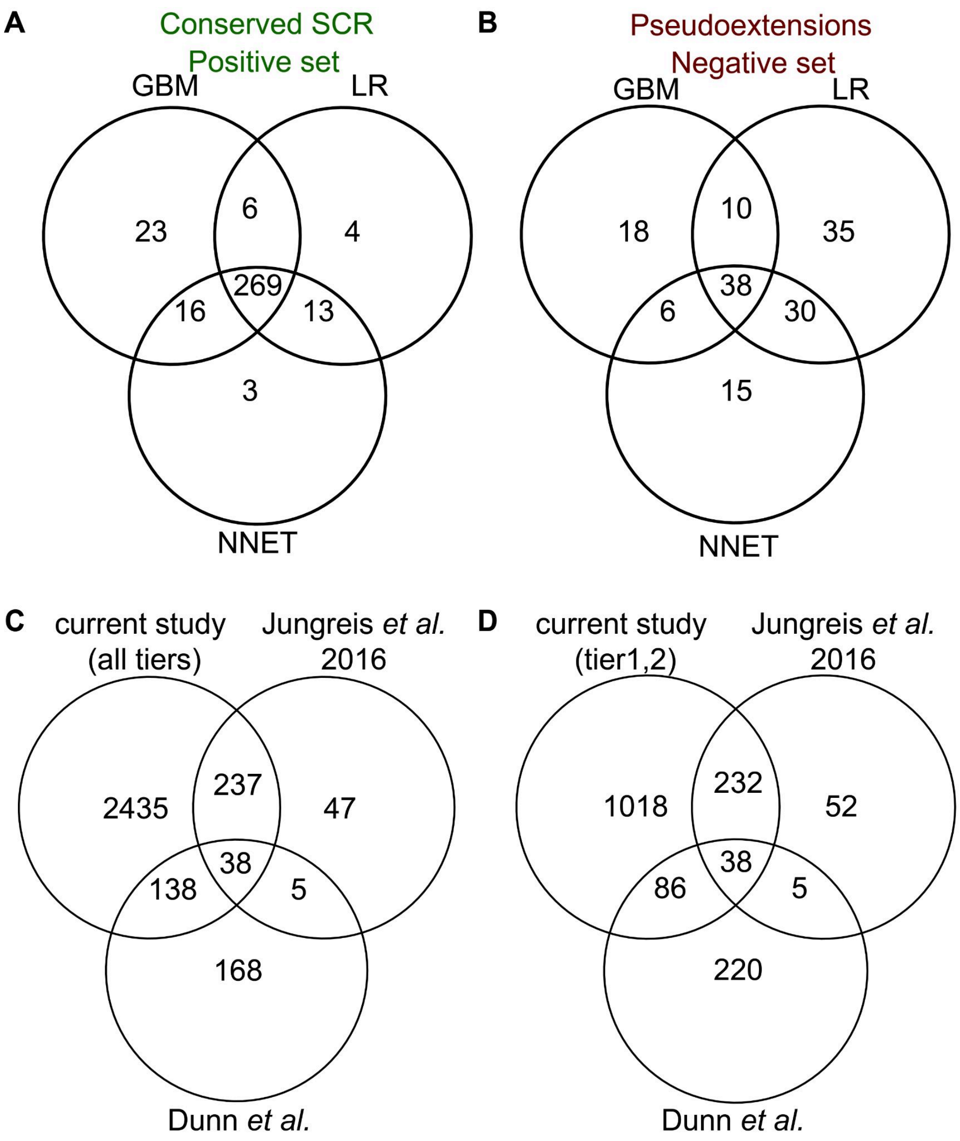


**Supplementary Figure S4.** Venn diagrams showing overlapping between gene sets.

**(A)** Overlap of SCR predictions within the positive training set. Predictions were obtained using Gradient Boosting Machine (GBM), Logistic Regression (LR), and Neural Network (NNET) classifiers. These genes were included as SCR candidates and were evaluated during the scoring procedure.
**(B)** Overlap of SCR predictions within the negative training set from ultrastop genes and their pseudoextension. These genes were not included as SCR candidates.
**(C)** Overlap of SCR candidate genes identified by the previous largest conservation-based study [23], the previous ribosome profiling SCR study in *Drosophila* [24], and the current study (2848 genes from all tiers). Overlaps are statistically overrepresented over random subsets: Fisher's exact test: PhyloCSF set [23], $p = 5.3 \times 10^{-139}$; ribosome profiling set [24], $p = 8.6 \times 10^{-39}$.
**(D)** Overlap of SCR candidate genes identified by the previous largest conservation-based study [23], the previous ribosome profiling SCR study in *Drosophila* [24], and the current study (1374 genes from tiers 1,2). Overlaps are statistically overrepresented over random subsets: Fisher's exact test: PhyloCSF set [23], $p = 2.5 \times 10^{-221}$; ribosome profiling set [24], $p = 2.6 \times 10^{-41}$.

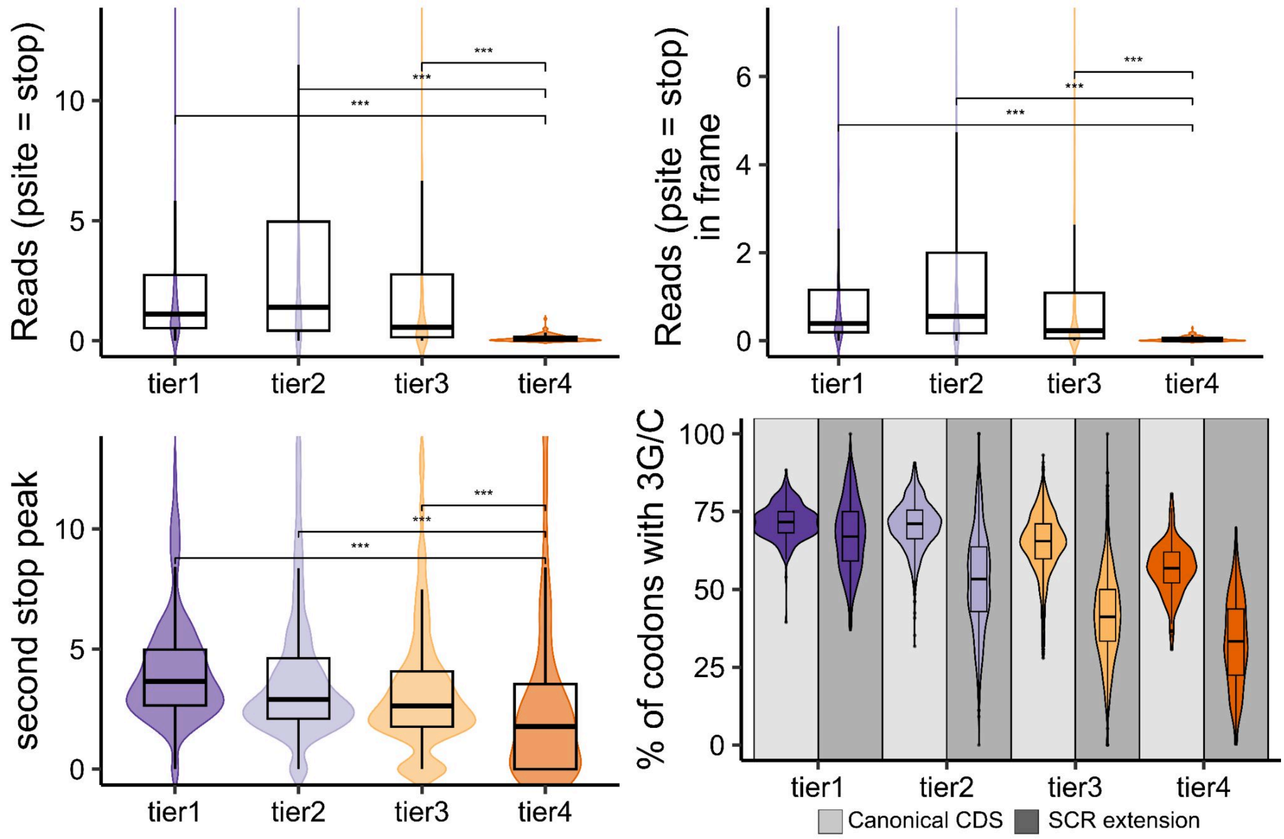


**Supplementary Figure S5.** Metrics used for SCR scoring.

**Top left**: "Stop reads" correspond to P-site-resolved ribosome footprints whose inferred P-site overlapped the stop codon, ensuring that only ribosomes actively decoding the termination codon are quantified rather than all reads merely spanning the region. For each gene, the mean of this value was calculated across Ribo-Seq samples.
**Top right**: "In-frame stop reads" denote the subset of stop reads whose P-site aligned with the canonical CDS reading frame, reflecting ribosomes elongating in-frame at the stop codon. For each gene, the mean of this value was calculated across Ribo-Seq samples.
**Bottom left:** "Second-stop peak" represents the fold change between the highest coverage value within the last 12nt of the SCR extension and the mean value in the same window, capturing the sharp termination peak expected at the second stop codon.
**Bottom right:** percentage of the codons ending with G/C in Canonical CDS (light grey background) or in SCR extension (dark grey background)
Statistical comparisons are indicated (NS, not significant; **, $p < 0.01$; ***, $p < 0.001$, two-sided Wilcoxon rank-sum test)

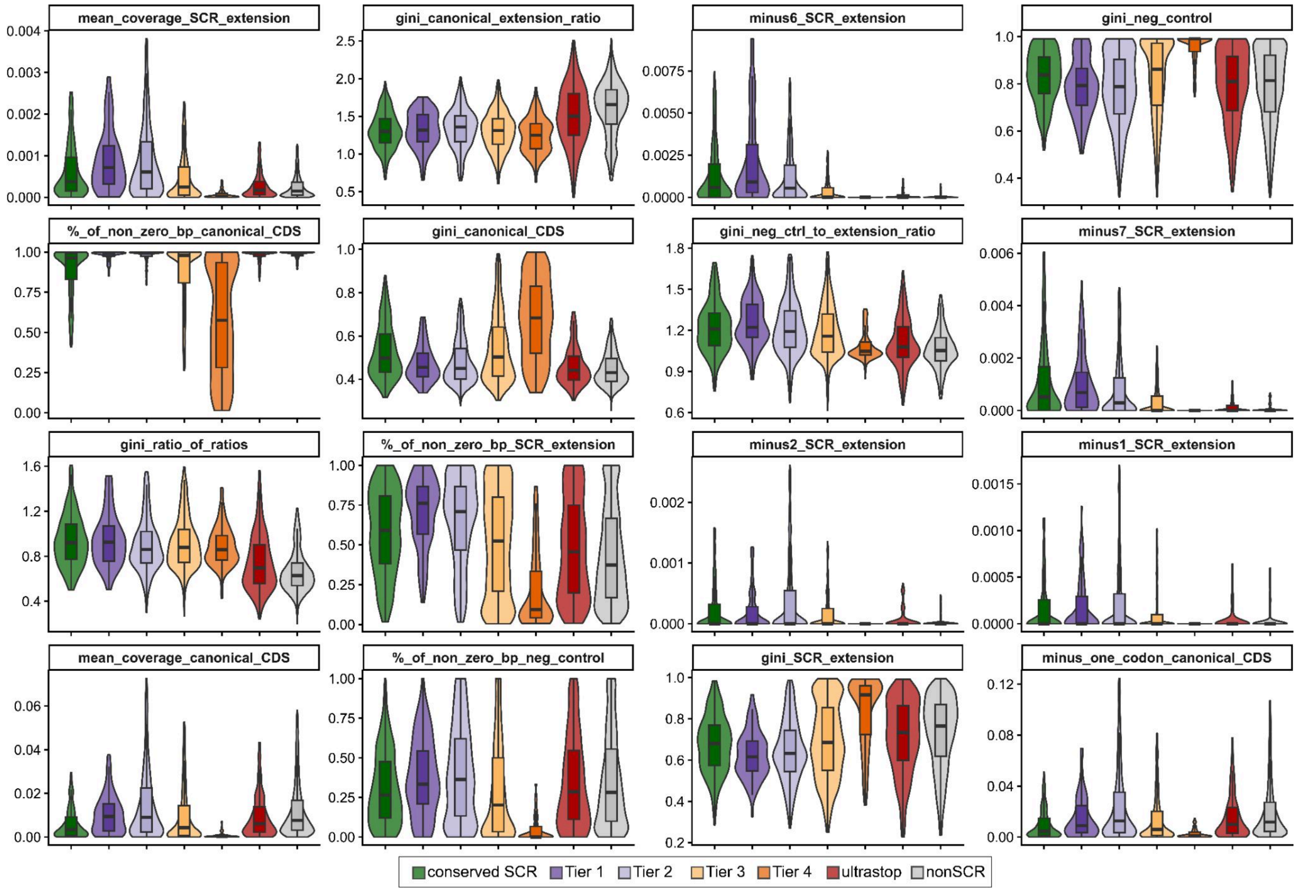


**Supplementary Figure S6.** Distribution of Ribo-Seq features used for machine-learning prediction across SCR confidence groups (see description in Methods). Violin and boxplots show ML input features for conserved SCR genes used as positive training examples, ultrastop genes used as negative training examples, ML-predicted genes assigned to tiers based on the composite SCR score, and non-SCR genes not predicted by any model. Features are ordered by aggregate importance across classifiers. Outliers were removed for visualization.

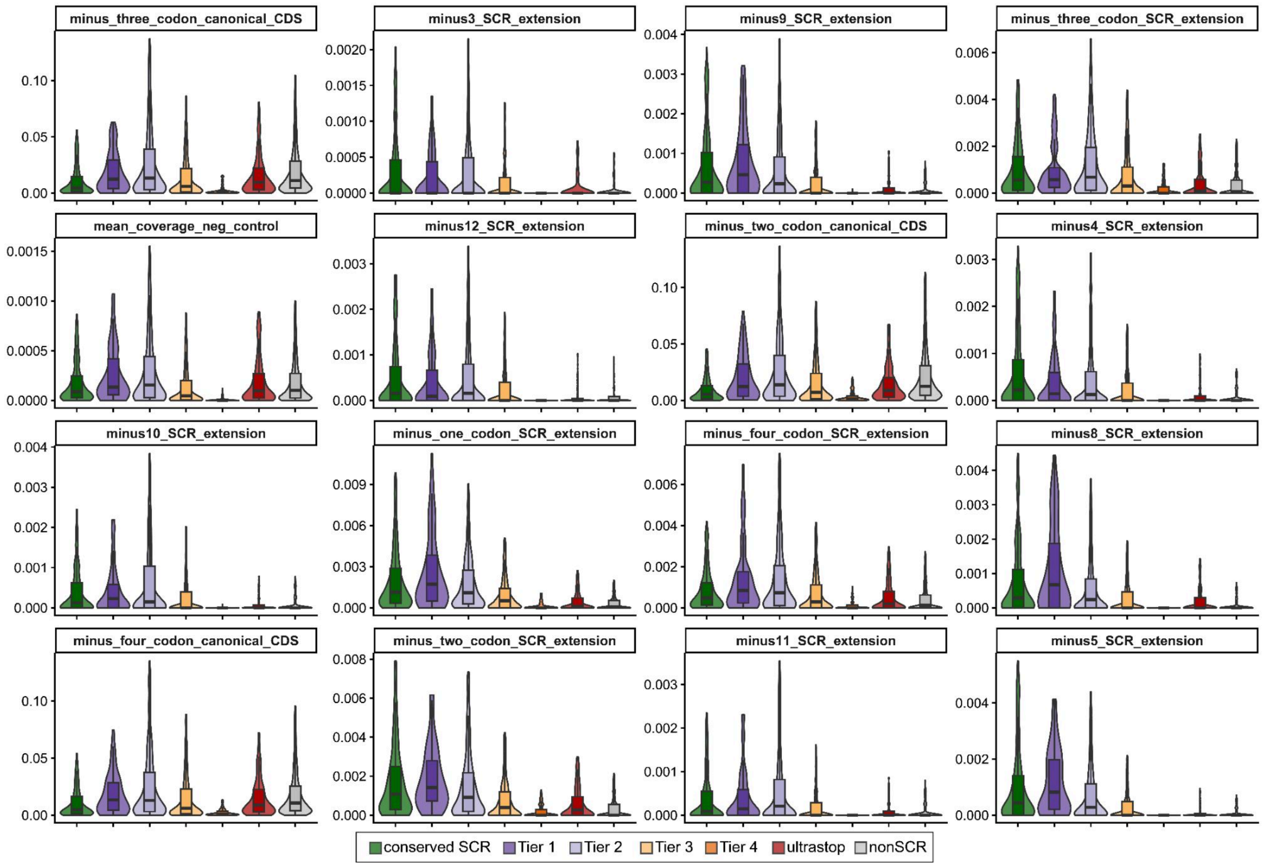


**Supplementary Figure S7.**

Distribution of Ribo-Seq features used for machine-learning prediction across SCR confidence groups (see description in Methods). Violin and boxplots show ML input features for conserved SCR genes used as positive training examples, ultrastop genes used as negative training examples, ML-predicted genes assigned to tiers based on the composite SCR score, and non-SCR genes not predicted by any model. Features are ordered by aggregate importance across classifiers. Outliers were removed for visualization.

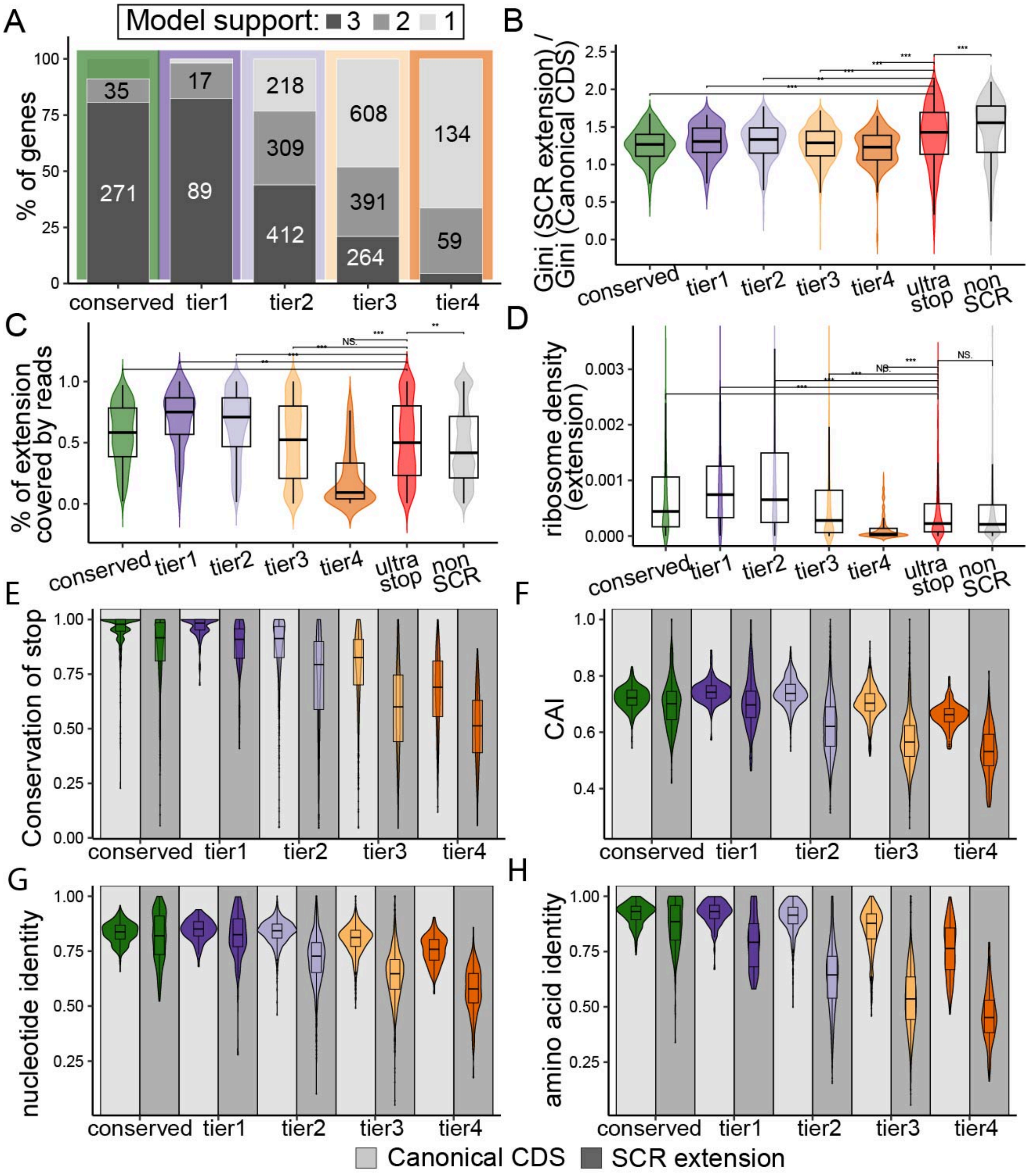


**Supplementary Figure S8.** SCR score tiers reflect both Ribo-Seq support and evolutionary signatures of programmable stop codon readthrough.

**(A)** Proportion of genes supported by 1,2,or 3 ML models among conserved SCR genes (used as the positive training set), across predicted tiers (tier1-tier4), ultrastop genes (used as the negative training set), and genes for which SCR was not predicted by all three models.
**(B-D)** Ribo-Seq-based metrics across conserved SCR genes, predicted tiers, Ultrastop genes, and non-SCR genes.

**(B)** Ratio between gini coefficient of SCR extension and gini coefficient of canonical CDS.
**(C)** Percentage of SCR extension covered by ribosome footprints.
**(D)** Ribosome density within the SCR extension.

**(E-H)** Evolutionary and sequence-based features across conserved SCR genes and predicted tiers.
**(E)** Stop codon conservation.
**(F)** Codon Adaptation Index (CAI) of the SCR extension.
**(G)** Nucleotide identity across species.
**(H)** Amino acid identity across species.

Statistical comparisons are indicated (NS, not significant; **, $p < 0.01$; ***, $p < 0.001$, two-sided Wilcoxon rank-sum test)

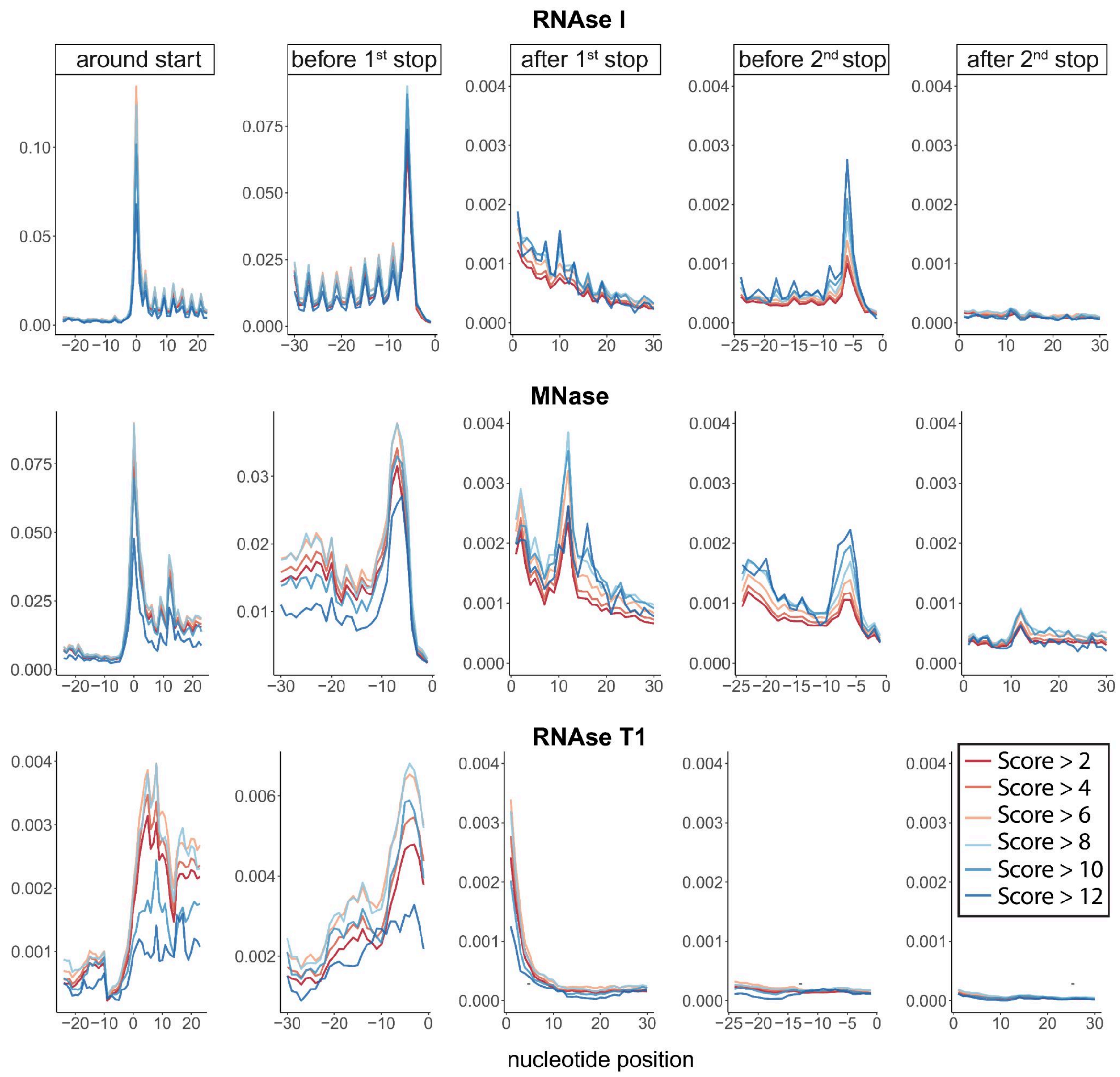


**Supplementary Figure S9.** Aggregated Ribo-Seq density stratified by SCR score.
Cumulative analysis was performed by grouping genes based on increasing SCR score thresholds (indicated by color and shown in the legend). Supplementary Figure S7 contains an analogous analysis grouping genes by tiers. Horizontal facets represent different transcript regions (around the start codon, end of the canonical CDS, beginning of the SCR extension, and end of the SCR extension), while vertical facets correspond to the nucleases used to generate the aggregated datasets. Only reads of 28-35 nt were considered. For MNase- and RNase I-treated samples, length-specific P-site offsets were applied individually for each sample. In contrast, for RNase T1-treated samples, a fixed 12-nt offset was trimmed from both ends of each read. RNase I and MNase datasets show robust profiles, whereas the single RNase T1 dataset has weaker signal but preserved coding enrichment, so that it was retained in density analyses. RNase I improved 3-nt periodicity and reduced post-stop density, as reported [79]. A termination peak at the second stop is evident across digestion types in tier 1-2 genes, supporting the robustness of our SCR predictions. Periodicity in SCR regions is weak overall, but clearly detectable in RNase I samples for genes with top SCR scores. MNase samples showed an additional ~12 nt downstream peak, independent of SCR. Its origin remains unclear and may reflect an MNase-specific termination signature in insect samples. Overall, this analysis shows that increasing SCR score is associated with a stronger termination peak and improved 3-nt periodicity in the SCR extension, particularly evident in RNase I-derived samples.

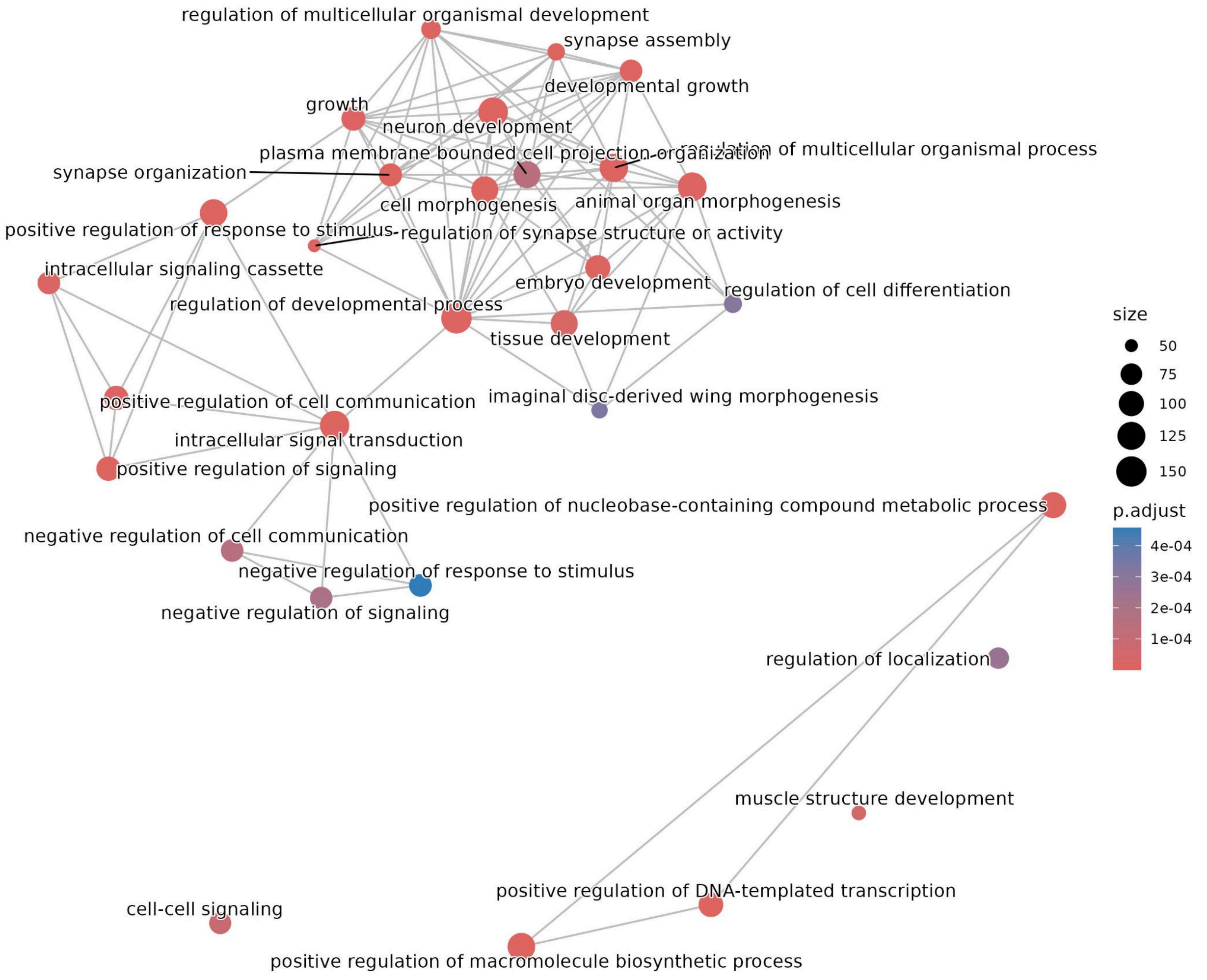


**Supplementary Figure S10.** GO enrichment of genes from high SCR scored genes (tier 1 and tier 2 together). Node size reflects the number of genes in each GO term. Terms sharing similar gene sets are positioned closer together and connected by grey lines.

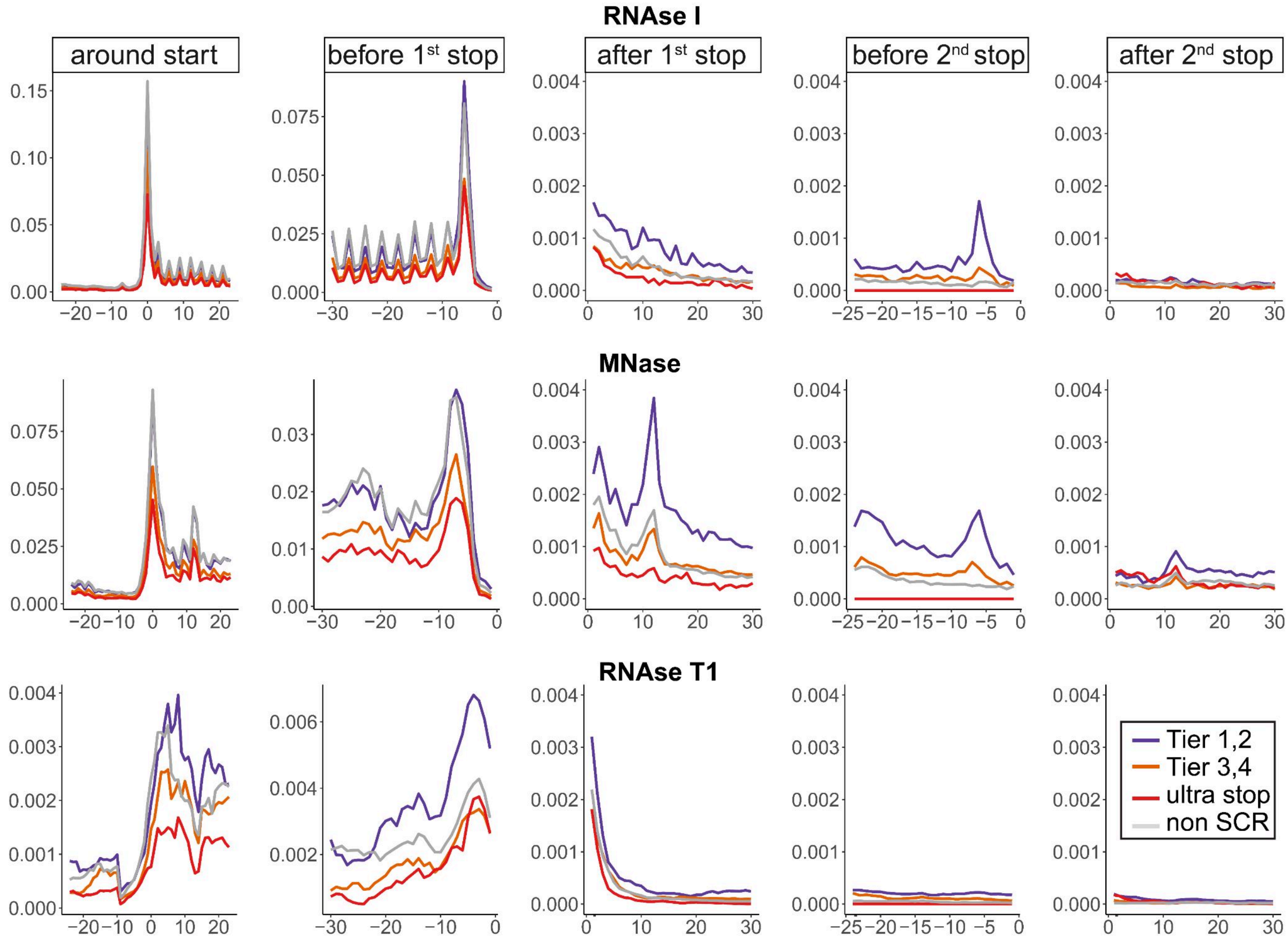


**Supplementary Figure S11.** Aggregated Ribo-Seq density stratified by tiers. See caption of analogous analysis in Supplementary Figure S9. Genes were grouped here by SCR tier, with ultrastop and non-SCR genes included as controls.

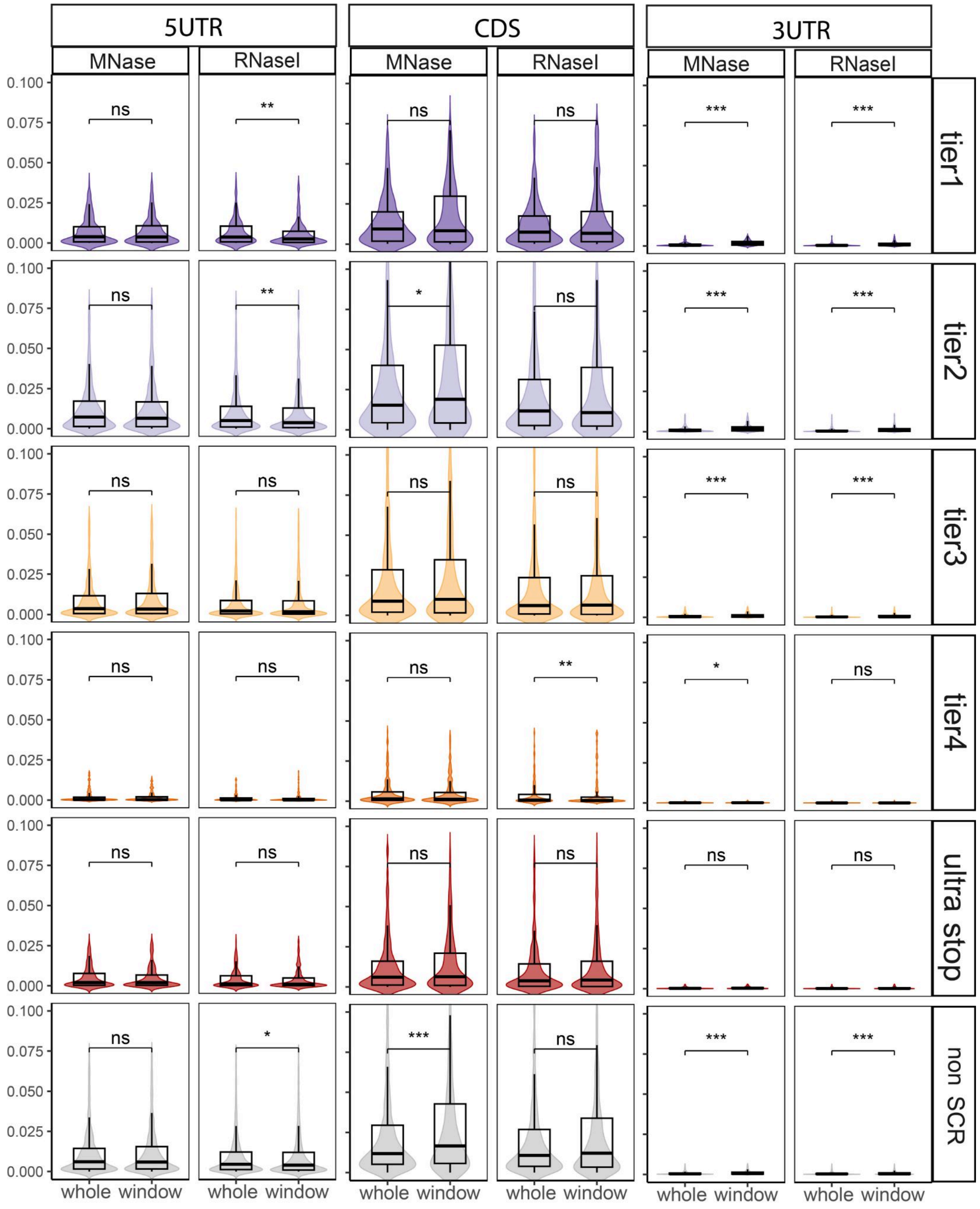


**Supplementary Figure S12**. Comparison of ribosome density across regions using aggregated datasets stratified by nuclease. "Whole" values correspond to the mean density across all nucleotides, computed for each gene and presented as a distribution across genes. While "window" values correspond to defined regions (last 24 nt of the 5′UTR, first 24 nt of the CDS, and first 30 nt of the 3′UTR). Significance is indicated as NS, *, **, and *** ($p < 0.05$, 0.01, and 0.001, respectively; paired two-sided Wilcoxon test).

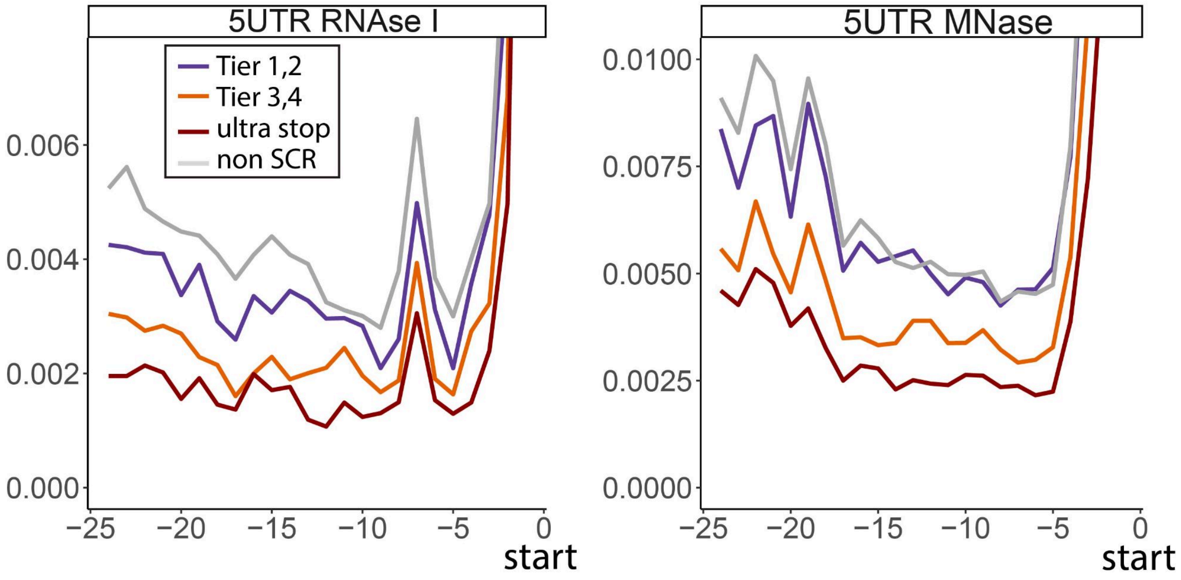


**Supplementary Figure S13.** Elevated Ribo-Seq density in 5'UTR does not show periodicity and has similar patterns across different gene groups. 0 - first nt of start codon. Only reads of 28-35 nt were considered. For MNase- and RNase I-treated samples, length-specific P-site offsets were applied individually for each sample.

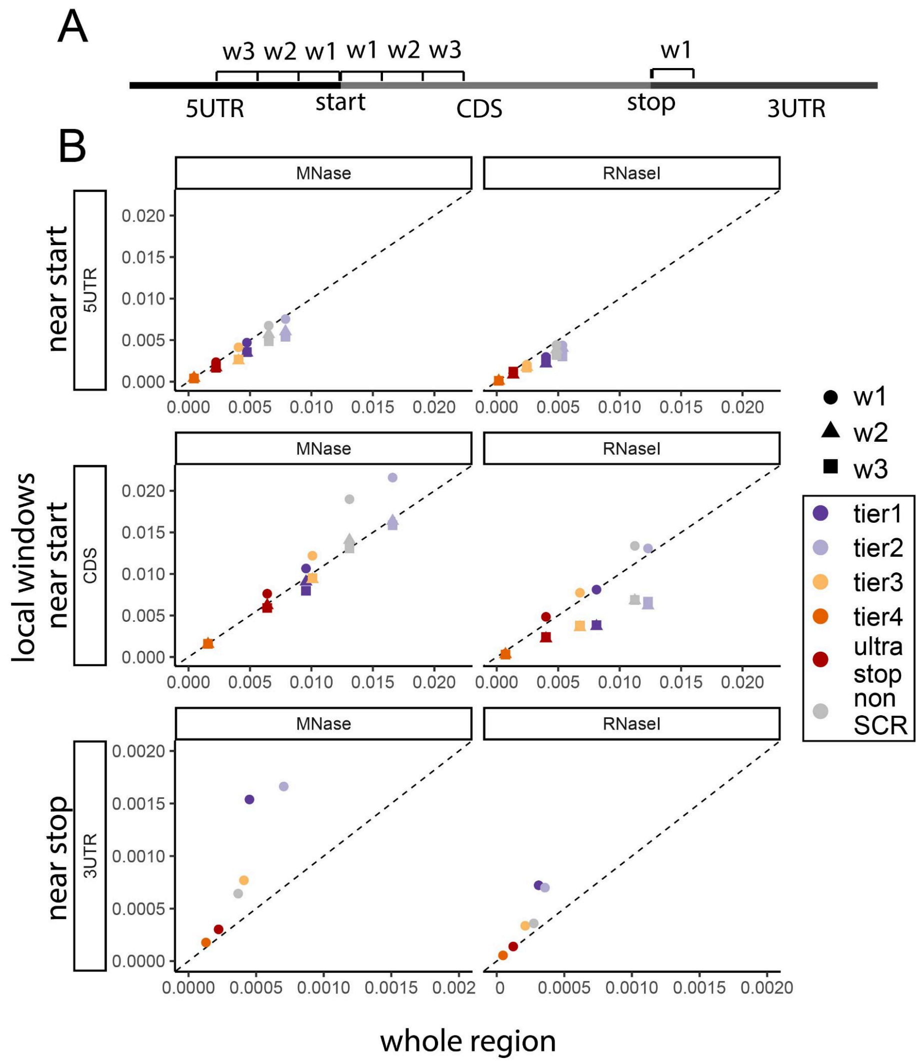


**Supplementary Figure S14.** Comparison of ribosome density between whole transcript regions (5'UTR, CDS, 3'UTR) and selected windows in these regions.

**(A)** Schematic of regions used for ribosome density analysis. For each gene, ribosome density was quantified either as the mean across full transcript regions (5′UTR, CDS, and 3′UTR) or as the mean within defined windows (w1-w3 for the CDS and 5′UTR; w1 only for the 3′UTR). Density was first aggregated at each nucleotide position across Ribo-Seq samples, later mean values for defined regions were calculated per each gene and lastly median densities were subsequently calculated across genes within each tier, yielding a single value per tier. Window size was 24 nt. For example, w1 in the 5′UTR corresponds to the last 24 nt upstream of the start codon, while w2 spans positions -48 to -24 relative to the start codon. Localized effects are expected to produce higher density in specific windows compared to the mean across the full region (illustrated here for the 3′UTR near the stop codon, corresponding to the SCR region). In contrast, regions with uniform ribosome occupancy, such as the 5′UTR, are expected to show similar densities across windows and the full region.
**(B)** Median ribosome profiling density per tier group, computed either over the entire region (x-axis) or within specific windows of the same region (y-axis; 24 nt windows for 5′UTR and CDS, and the first 30 nt for 3′UTR). Facets are organized by genomic region (rows) and nuclease type (columns). The dashed line indicates the identity (1:1) relationship. For the 3′UTR, axis limits were adjusted (10-fold lower scale) to improve visualization of the data.

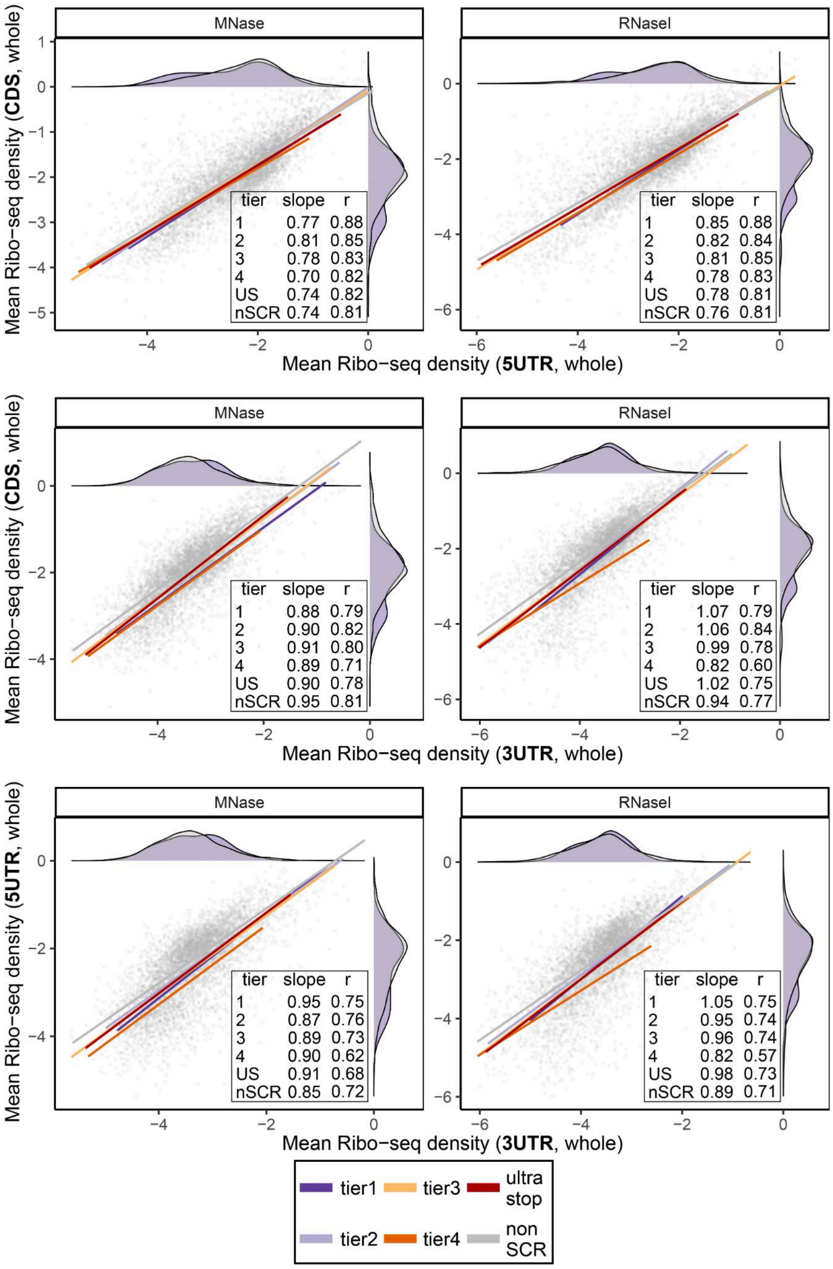


**Supplementary Figure S15.** Correlation between ribosome profiling densities across full-length 5′UTR, CDS, and 3′UTR regions.

For each gene, mean ribosome density was computed across all nucleotides in each region using aggregated datasets stratified by nuclease type, and $\log_{10}$-transformed. Genes were grouped into tiers, and linear regressions were fitted for pairwise comparisons (CDS vs 5′UTR, CDS vs 3′UTR, and 5′UTR vs 3′UTR). Regression slopes were used to compare relative scaling between regions across tiers ,while Pearson correlation coefficients (R) were used to quantify the strength of association between regional densities. Density plots are shown only for non-SCR and high-SCR genes (tier 1).

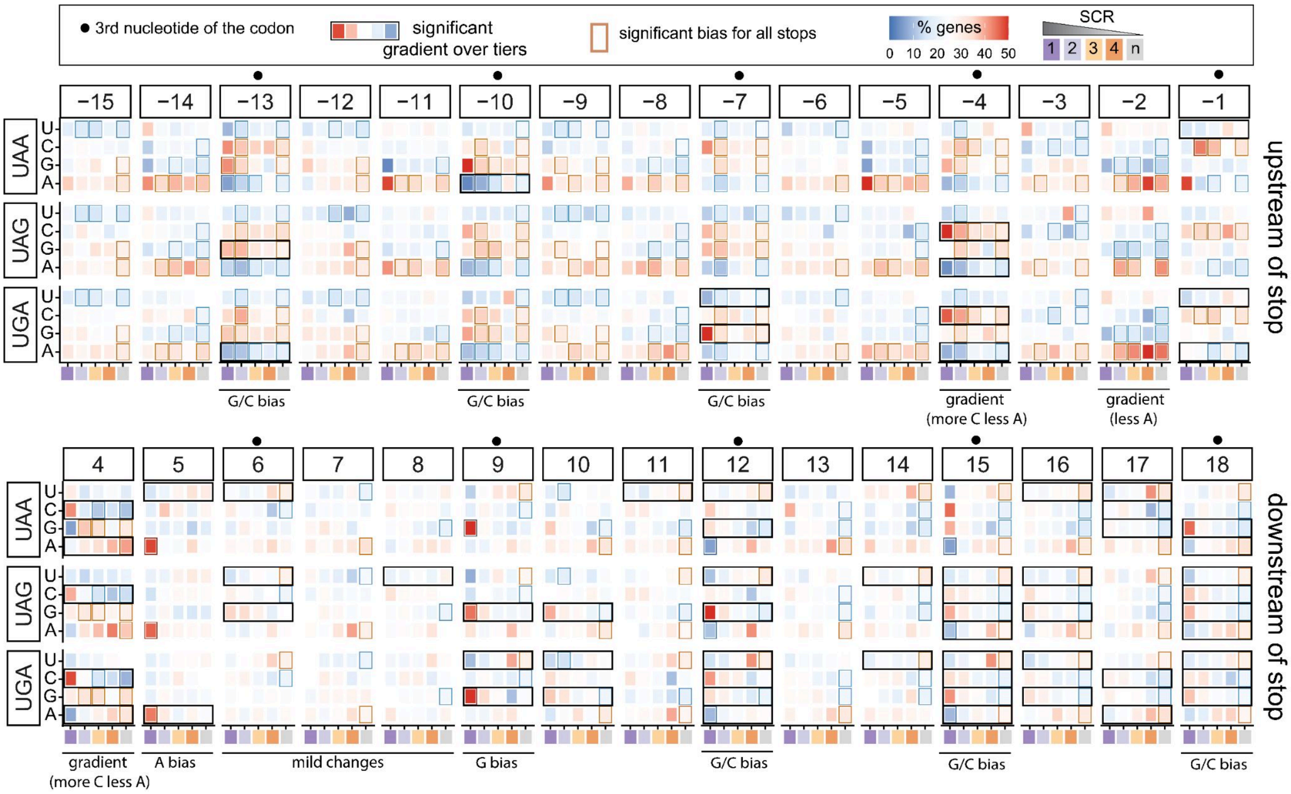


**Supplementary Figure S16.** Extended nucleotide context around stop codons.

For each tier (x-axis) and each stop codon (facet rows), the percentage of genes containing each nucleotide at each position relative to the stop codon is shown (facet columns). Top panel represents position upstream of stop codon, bottom panel represents nucleotide context downstream of stop codons. Position +4 corresponds to the nucleotide immediately downstream of the stop codon, whereas position -1 corresponds to the nucleotide immediately upstream. Borders around tiles indicate significant deviation from the expected 25% frequency. Significance was assessed separately for each of the three stop codons (FDR < 0.05 for each stop codon) and additionally required a combined FDR < 0.001 using Fisher's method to aggregate p-values across all three stop codons. Black rectangles indicate positions identified by regression analysis, in which nucleotide frequencies across tiers were tested for association with SCR propensity. Only nucleotides showing significant gradients (FDR < 0.001) are highlighted.

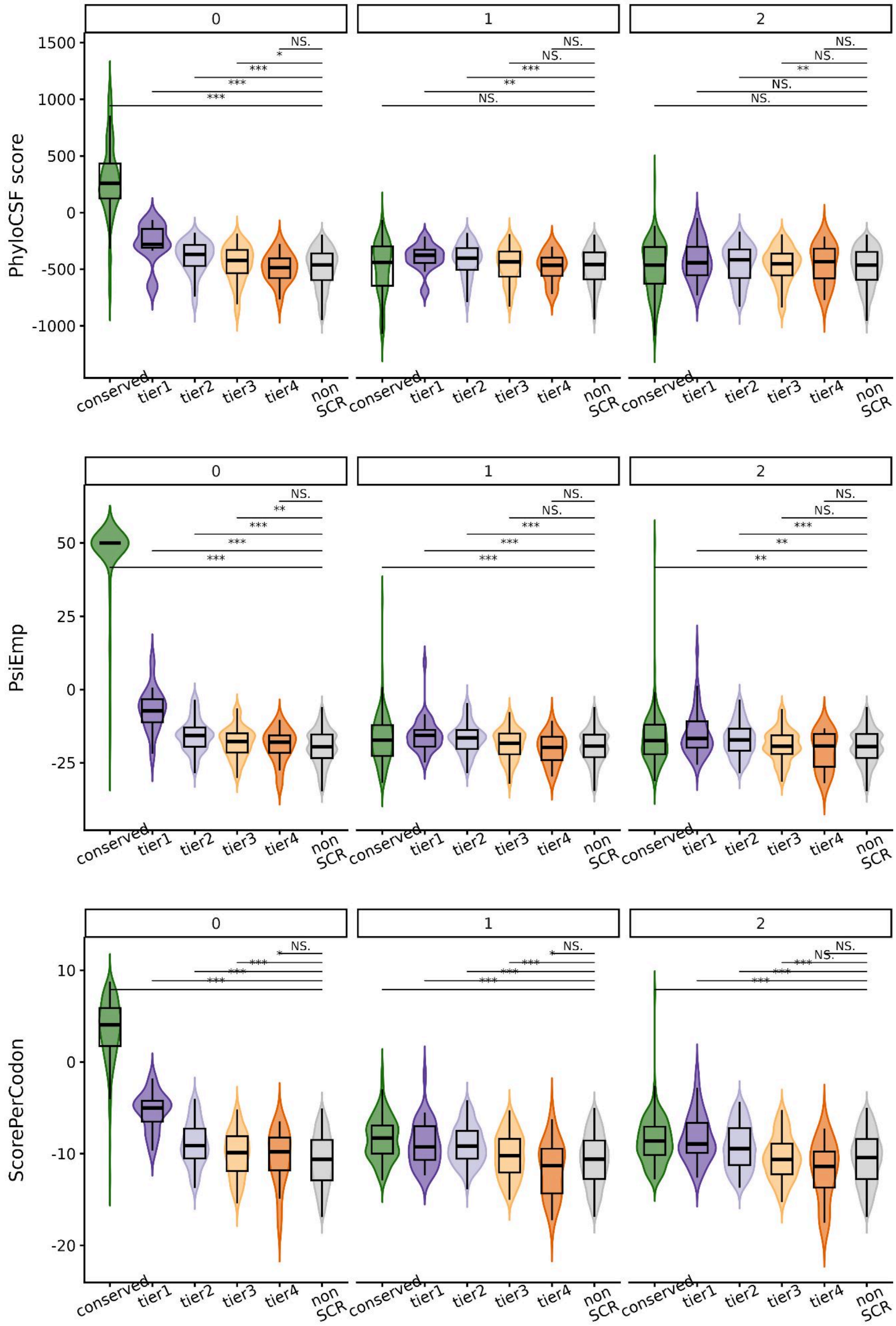


**Supplementary Figure S17.** PhyloCSF scores for genes in different tiers for 0 frame (in the same frame as canonical CDS) and two others (horizontal facets). Conserved SCR genes (Methods) were separated to assess how newly predicted candidates compare. Significance is indicated as NS, *, **, and *** ($p < 0.05$, 0.01, and 0.001, respectively; two-sided Wilcoxon test). Top: PhyloCSF scores calculated across the entire SCR extension region. Middle: PhyloCSF-ΨEmp scores, which account for the dependence of PhyloCSF scores on region length by comparing observed scores to empirical distributions of coding and non-coding regions of similar length. Bottom: Mean per-codon PhyloCSF scores, obtained by calculating PhyloCSF independently for each codon and summarizing the resulting distribution across the SCR extension

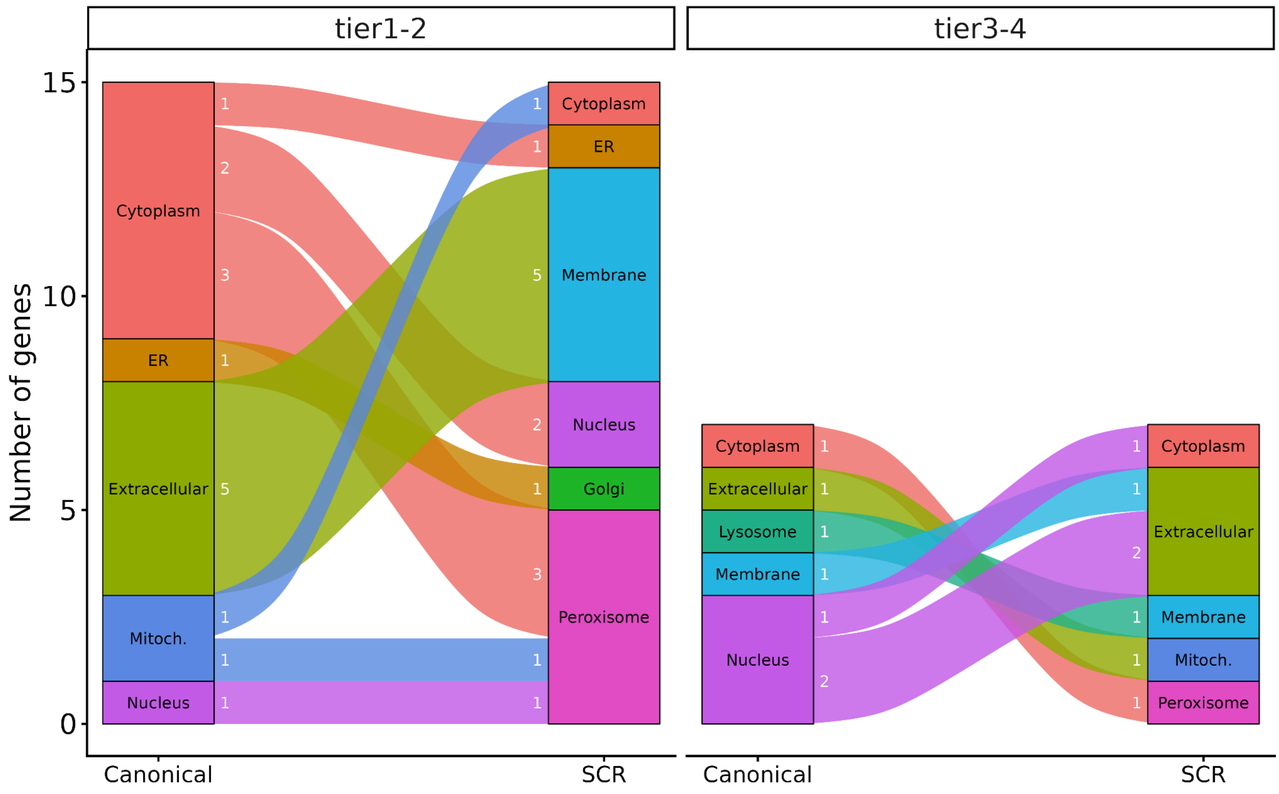


**Supplementary Figure S18.** Shifts in subcellular localization with SCR.
Changes in localization predicted by DeepLoc2 (Methods) between the canonical and SCR extended protein isoforms identified among candidate SCR genes are shown, grouped by tier1-2 (left) and tier 3-4 (right). ER: Endoplasmic reticulum. Mitoch: Mitochondrion.

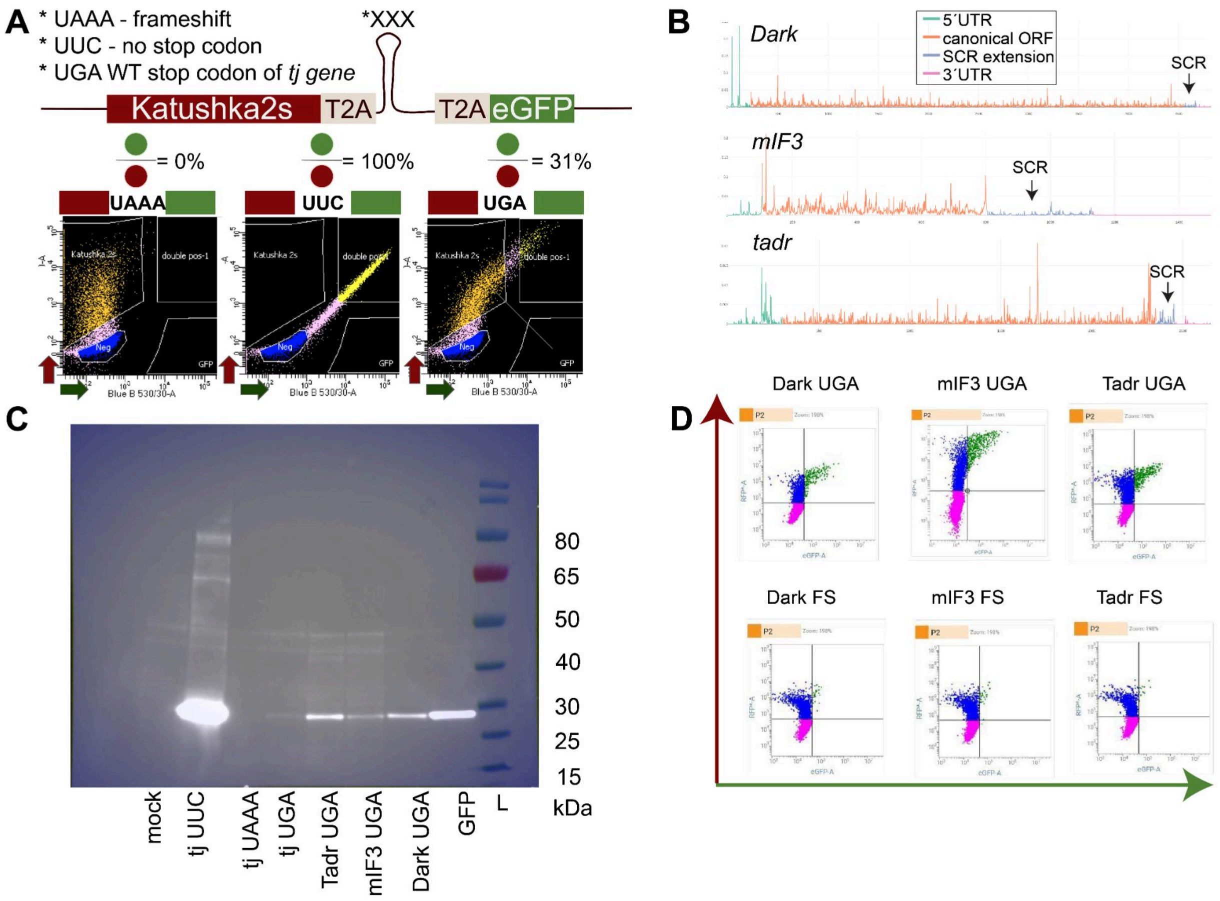


**Supplementary Figure S19.** Experimental assessment of SCR signals prior to large-scale screening.

**(A)** Pilot experiment using a dual-fluorescence reporter containing the SCR insert from the *tj* gene.
**(B)** Aggregated ribosome profiling density for three genes selected for SCR validation. The SCR region is indicated by an arrow and highlighted in blue. These genes were selected based on these profiles and tested for SCR activity using western blot.
**(C)** Western blot analysis using an anti-GFP antibody in S2 cells transfected with plasmids carrying the selected inserts (see Supplementary Table T4). Due to the presence of T2A sites, the expected band size is ~30kDa, corresponding to GFP alone (last lane).
**(D)** Flow cytometry profiles of selected candidates compared with negative controls (frameshifted constructs = FS).

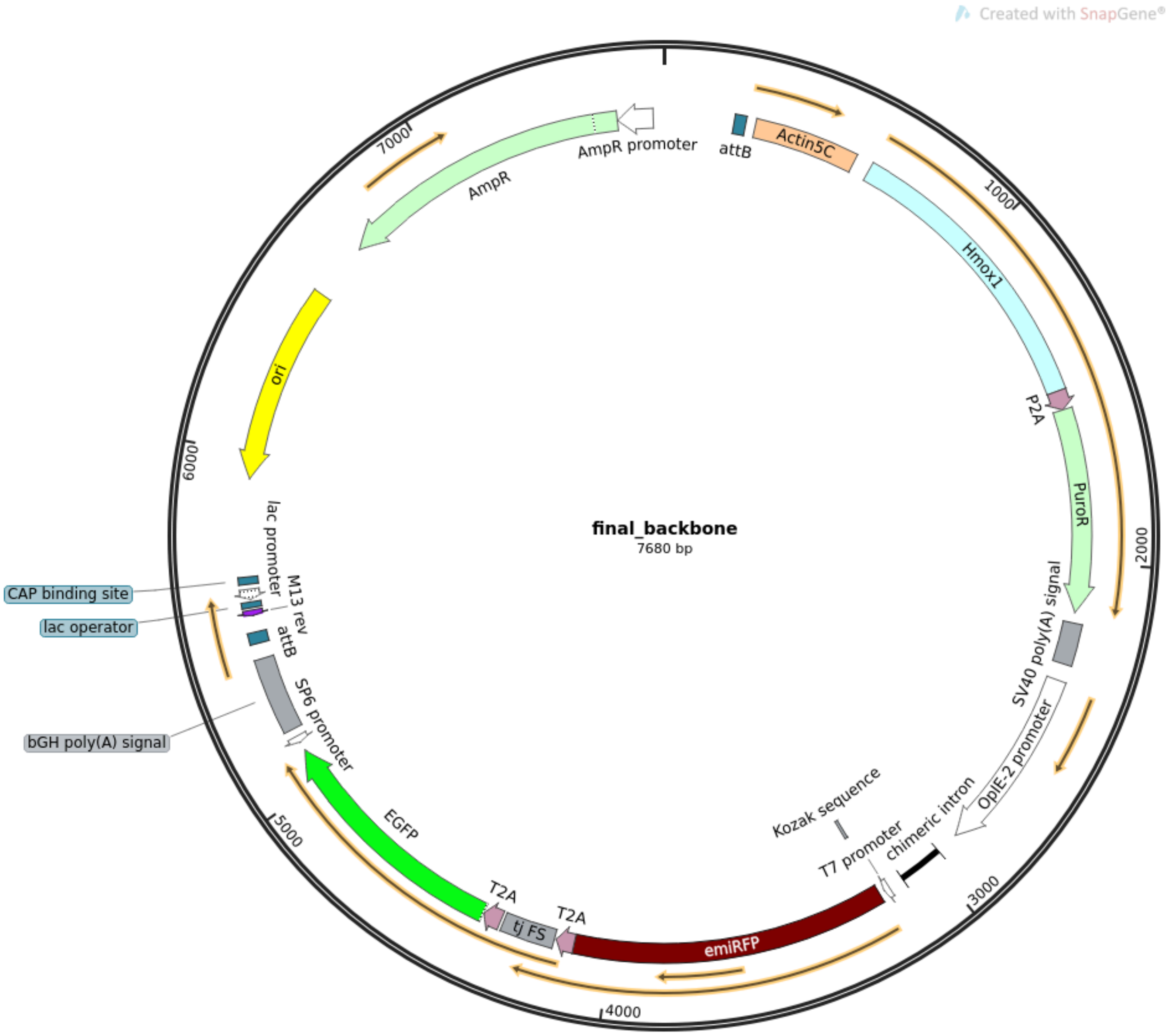


**Supplementary Figure S20.** Plasmid map of library backbone used in pooled screening.

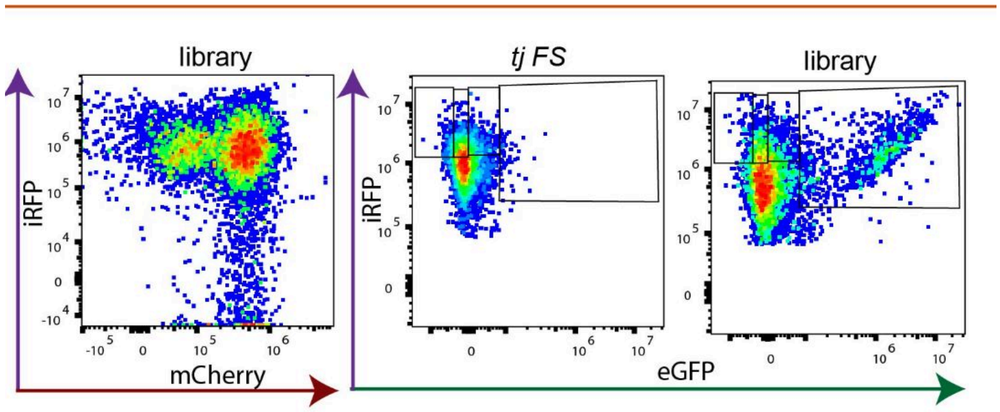


**Supplementary Figure S21.** Screening flow cytometry profiles.

**Left panel:** outcome of genomic integration based on mCherry (x-axis) and iRFP (y-axis) fluorescence. After integration, ~70% of cells are iRFP-positive, indicating high integration efficiency, although a fraction of cells still retains mCherry signal.
**Middle panel:** cells with integrated negative control (library backbone with *tj* frameshift insert, FS). x-axis: eGFP; y-axis: iRFP.
**Right panel:** cells with integrated library of SCR stimulators. x-axis: eGFP; y-axis: iRFP.

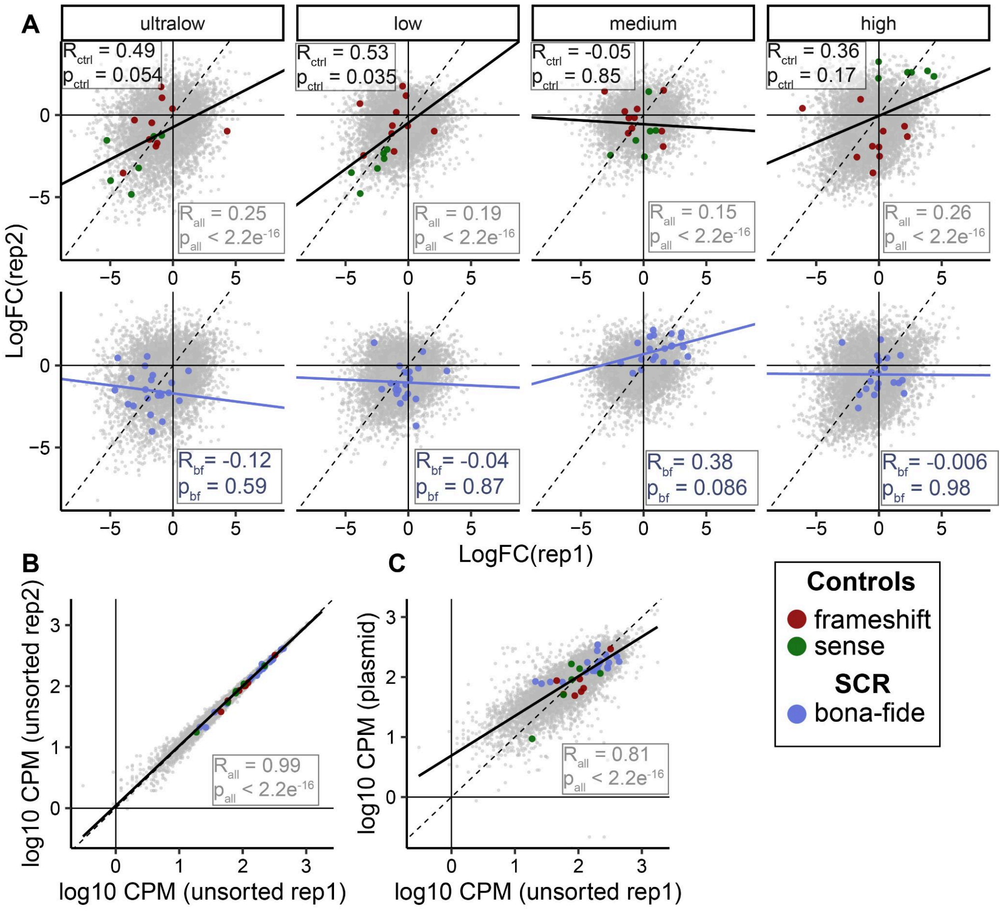


**Supplementary Figure S22.** Replicability of the screening.

**(A)** Enrichment of counts ($log_2FC$) in each sorted fraction relative to unsorted cells. Cells were sorted into four fractions (ultra-low, low, medium, and high) based on eGFP signal. Sorting was performed on two independent days, which were treated as biological replicates. For each insert, enrichment was calculated as the ratio of normalized counts in each fraction over unsorted cells (post-integration), independently for each replicate. Correlations between replicates are shown for each fraction. Pearson correlation coefficients (R) and associated p-values were computed for three groups of inserts: (i) all inserts (n = 10,052; grey), (ii) control sequences including frameshifted constructs ( no eGFP expected; n = 11) and sense controls (expected ~100% SCR; n = 6; black), and (iii) *bona fide* SCR genes previously validated by western blot and dual-fluorescence reporter assays, together with their evolutionary replicates (*Dark*, *mif3*, and *tadr*; light purple).

For sense constructs, enrichment is expected in the high fraction and depletion in the ultra-low fraction. For frameshifted constructs, enrichment is expected in the ultra-low fraction and depletion in the high fraction. For *bona fide* SCR elements, enrichment is expected in the medium fraction and depletion in the ultra-low fraction. Although correlations between replicates may be low, higher correlation coefficients are observed in the fractions where enrichment is predicted for each group.

**(B)** Correlation between $log_{10}$ CPM counts for replicates of unsorted cells after library integration.
**(C)** Correlation between $log_{10}$ CPM counts from replicate 1 of unsorted cells and counts obtained from sequencing of the plasmid library.

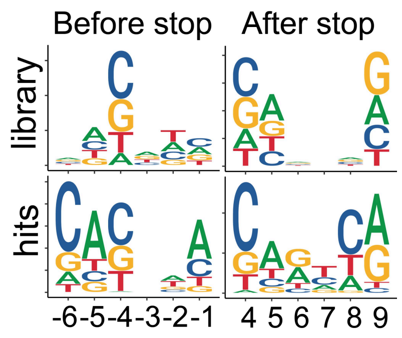


**Supplementary Figure S23.** Sequence logos of nucleotide composition (-6 to +6 nt relative to the stop codon). Top: input library. Bottom: screening hits (27 selected structures and their evolutionary replicates).

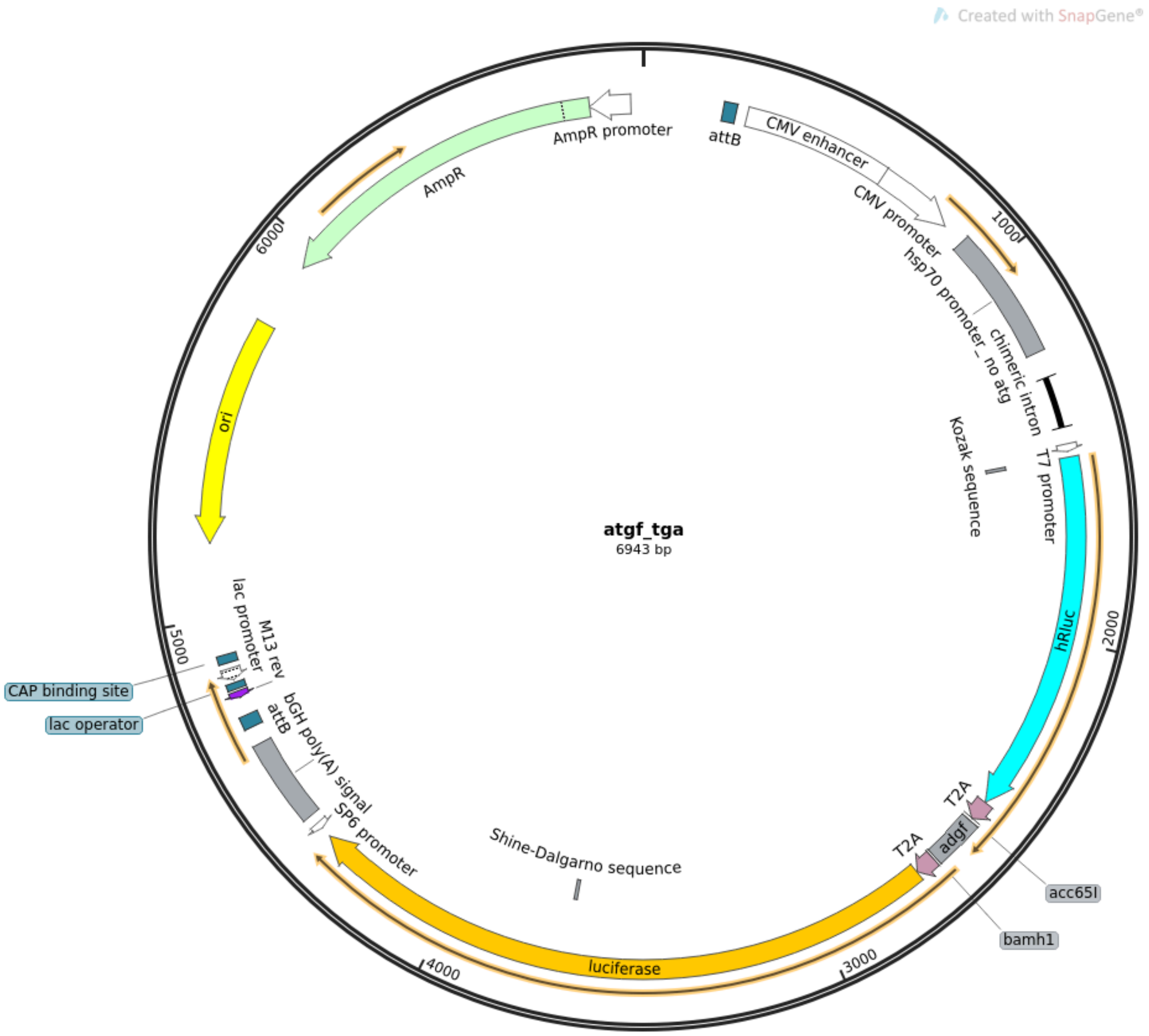


**Supplementary Figure S24.** Plasmid map of luciferase plasmid used in validation of the screening.

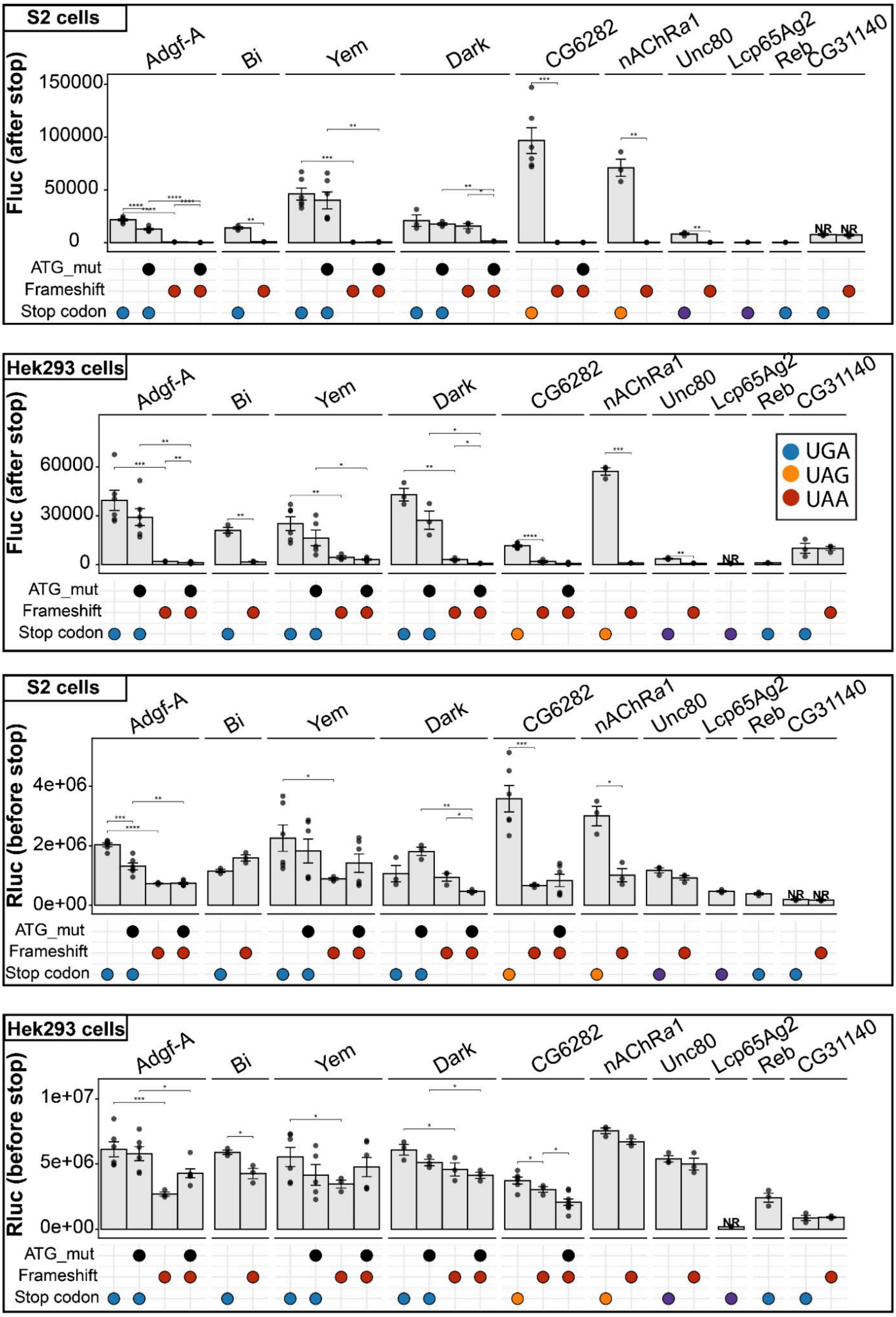


**Supplementary Figure S25. Absolute firefly luciferase (Fluc) and Renilla luciferase (Rluc) values measured in S2 and HEK293 cells for reporter constructs excluding the sense-control version.** Statistical significance is indicated as *, **, and *** ($p < 0.05$, 0.01, and 0.001, respectively) and was assessed using one-sided t-tests. Paired t-tests were applied when both groups contained three matched observations; otherwise, unpaired t-tests were used. NR - not reliable

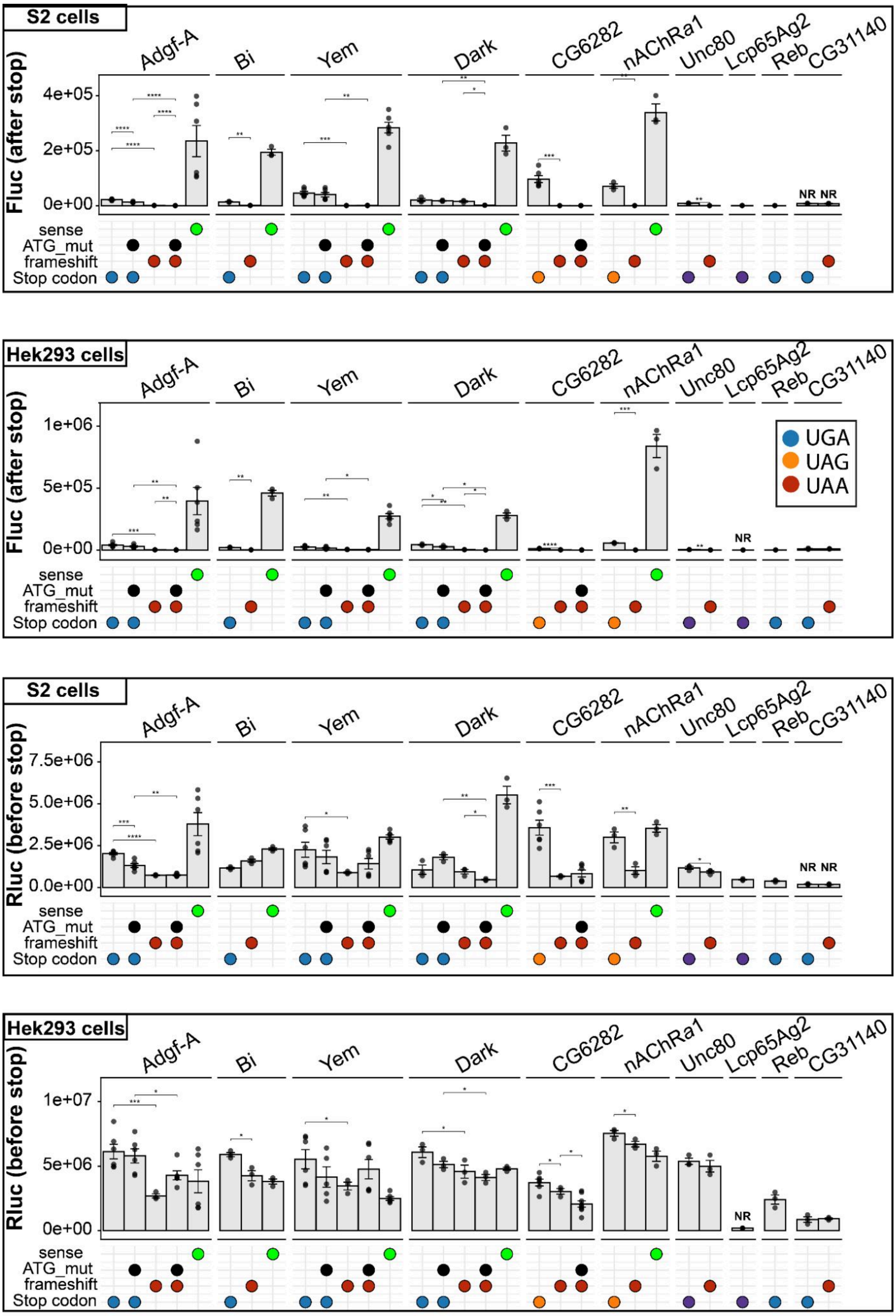


**Supplementary Figure S26.** Absolute firefly luciferase (Fluc) and Renilla luciferase (Rluc) values measured in S2 and HEK293 cells for reporter constructs including the sense-control **version.** Statistical significance is indicated as *, **, and *** ($p < 0.05$, 0.01, and 0.001, respectively) and was assessed using one-sided t-tests. Paired t-tests were applied when both groups contained three matched observations; otherwise, unpaired t-tests were used. NR - not reliable

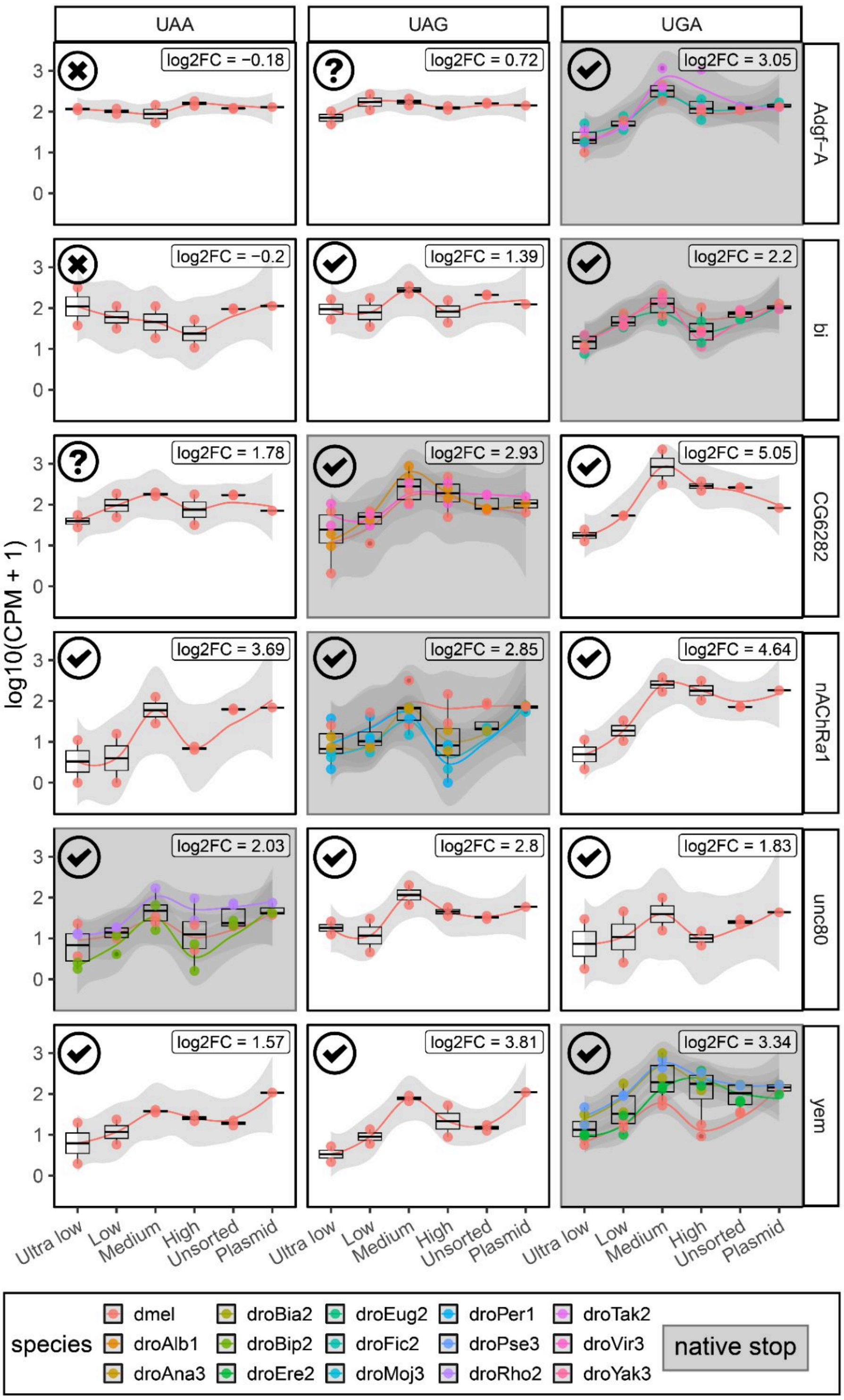


**Supplementary Figure S27. Activity of SCR stimulatory sequences (validated using dual luciferase assay) across different stop codons (UAA, UAG, UGA) measured in the screening assay.**

Each panel shows log10(CPM) read counts across sorted fractions. Red points indicate sequences from *Drosophila melanogaster*, whereas other colors correspond to evolutionary replicates. $Log_2FC$ represents the fold change between the medium fraction and the combined ultra-low and low fractions. Grey background indicates the native stop codon, and symbol shape in the left top corner represents the confidence of SCR activity.

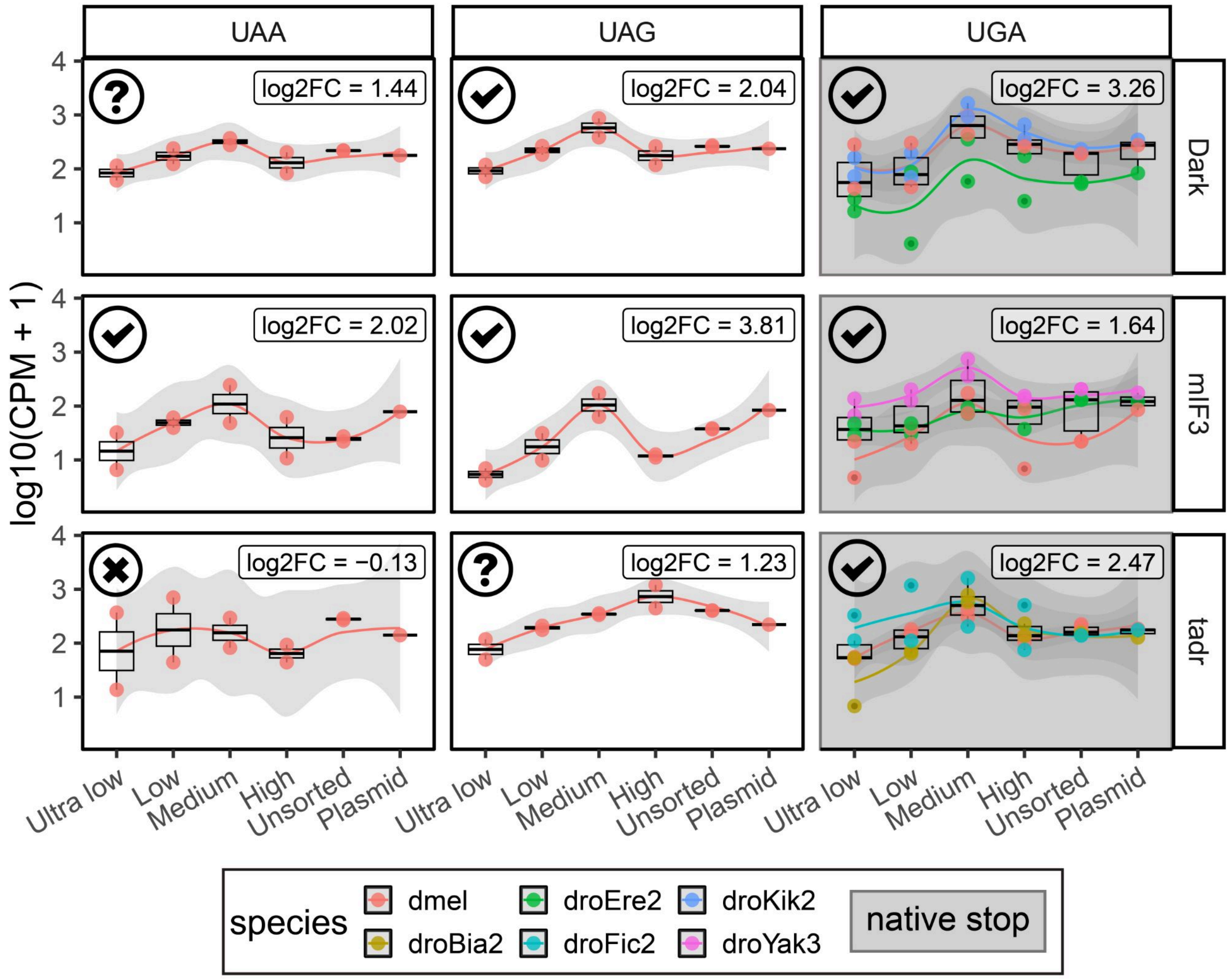


**Supplementary Figure S28.** Activity of SCR stimulatory sequences (bona-fide selected based on ribosome profiling) across different stop codons (UAA, UAG, UGA) measured in the screening assay.

Each panel shows log10(CPM, count per million) read counts across sorted fractions. Red points indicate sequences from *Drosophila melanogaster*, whereas other colors correspond to evolutionary replicates. $Log_2FC$ represents the fold change between the medium fraction and the combined ultra-low and low fractions. Grey background indicates the native stop codon, and the symbol shape in the left top corner represents the confidence of SCR activity.

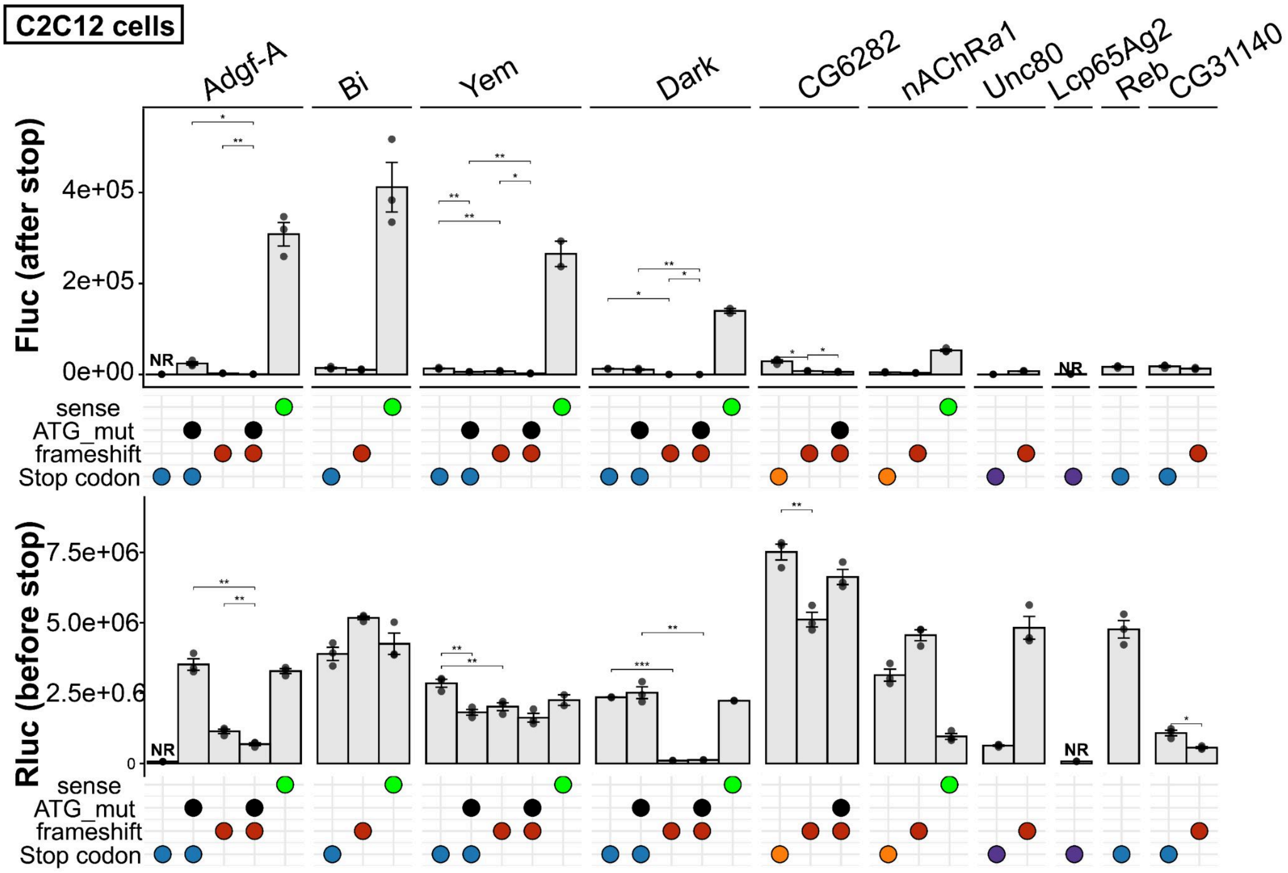


**Supplementary Figure S29.** Absolute firefly luciferase (Fluc) and Renilla luciferase (Rluc) values measured in C2C12 cells for reporter constructs including the sense-control version. Statistical significance is indicated as *, **, and *** ($p < 0.05$, 0.01, and 0.001, respectively) and was assessed using one-sided t-tests. Paired t-tests were applied when both groups contained three matched observations; otherwise, unpaired t-tests were used. NR - not reliable (Methods).

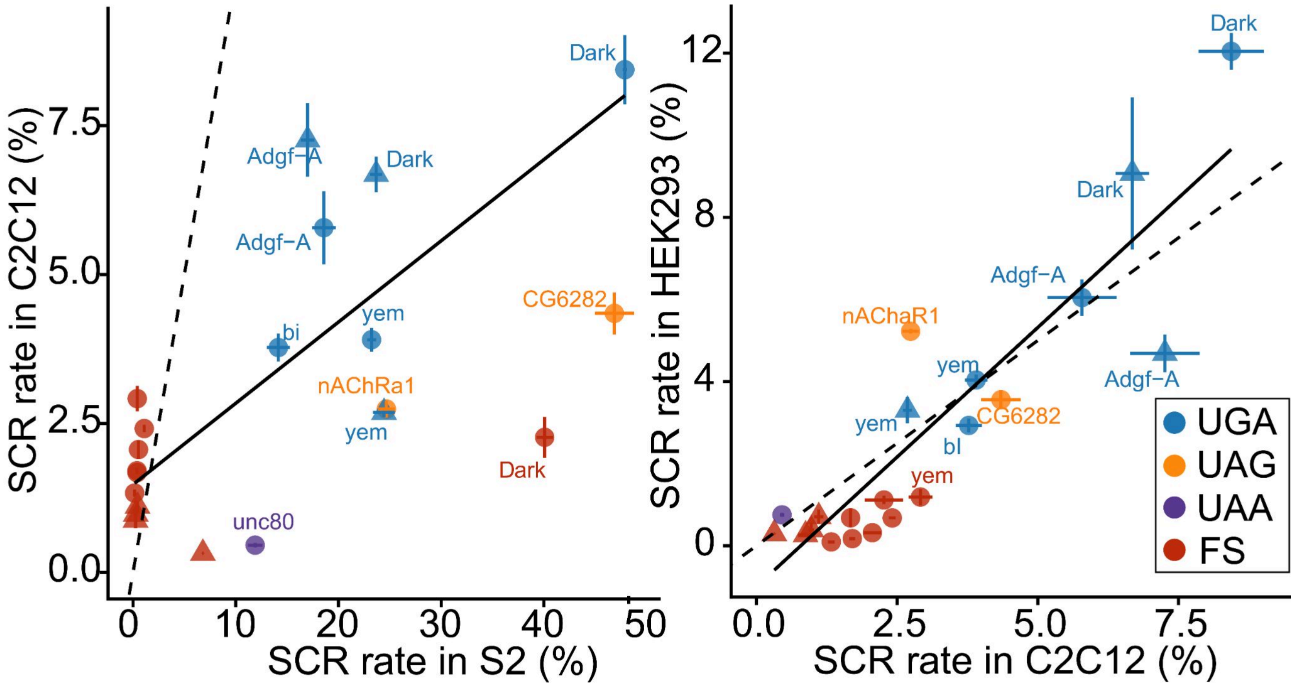


**Supplementary Figure S30. (A)** Correlation between SCR rates measured in S2, HEK293 and C2C12 cells via dual-luciferase. Only elements that passed validation were included. The color indicates the stop codon of the various constructs, with red ("FS") indicating frameshifted variants. Point shape indicates whether downstream AUG codons were mutated. The dotted line corresponds to a slope of 1, indicating identical SCR rates in both cell types. Correlation statistics for S2 vs. C2C12: R = 0.63, p.value = 0.002; or R = 0.74, p.value = 0.00063 when excluding CG6282 UAG and Dark FS. Correlation statistics for C2C12 vs. HEK293: R = 0.89, p = $3.42 \times 10^{-8}$. An analogous S2 vs HEK293 plot is presented in Figure 3.

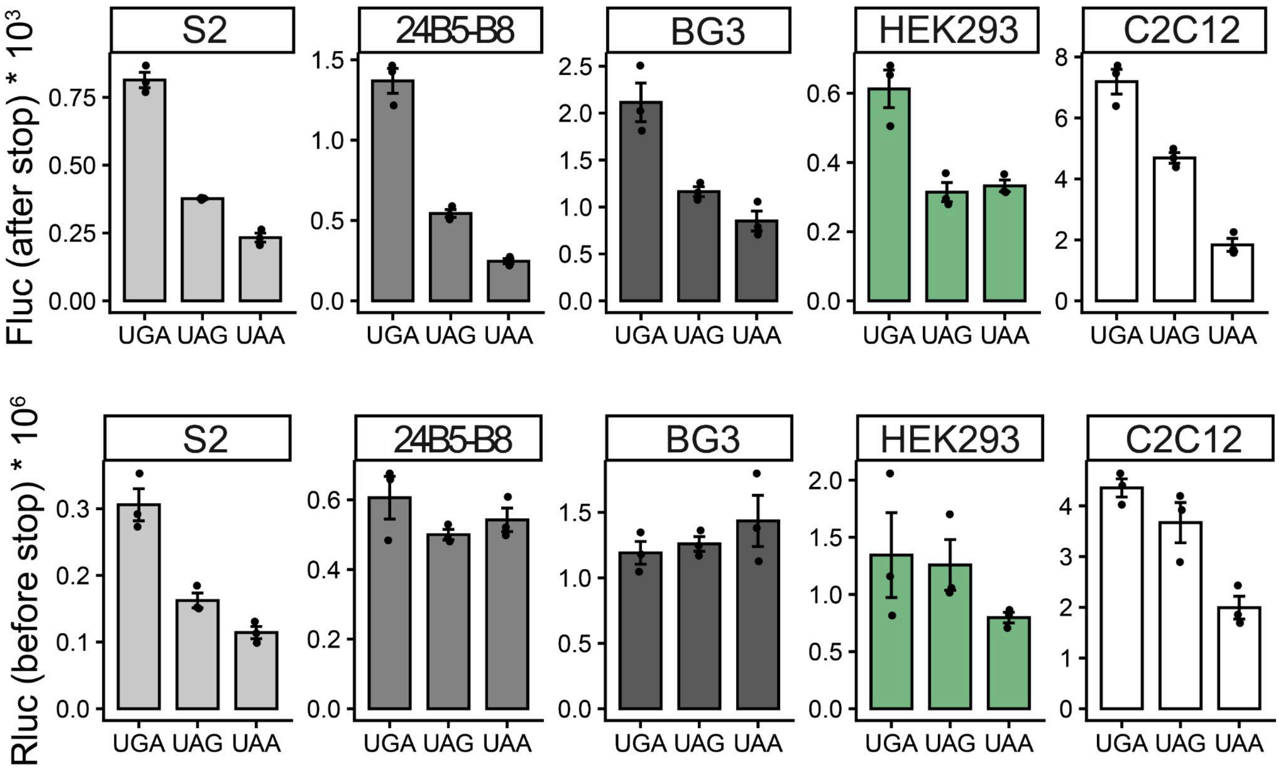


**Supplementary Figure S31.** Raw luminescent values from transfection experiments using reporters lacking stimulatory elements in Drosophila S2 cells, 24B5-B6, BG3 and mammalian HEK293 cells, and C2C12 cells.

These reporters do not contain stimulatory sequences and were designed to estimate basal termination efficiency. The nucleotide at the +4 position was set to C for all stop codons to mimic a weak termination context. The exact local context was GGUACC - stop - CGAUCC. Absolute values differ between cell lines due to differences in transfection efficiency and experimental protocols. (Rluc: Renilla; Fluc: Firefly)

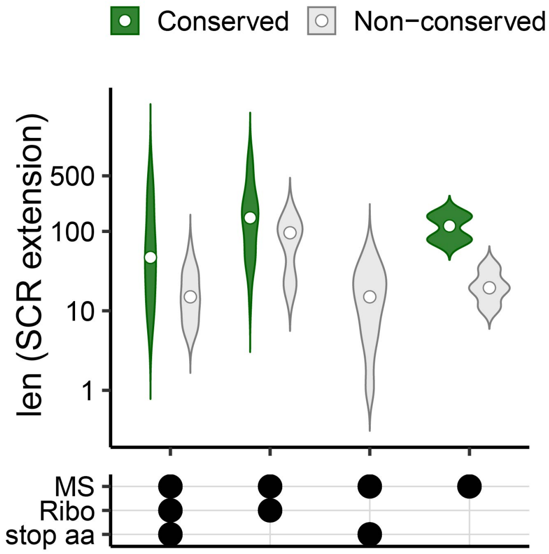


**Supplementary Figure S32.** Distribution of protein SCR extension lengths, grouped by detection method (MS, mass spectrometry; Ribo, ribosome profiling) and by peptide evidence type: peptides covering the stop codon, which allow inference of the amino acid incorporated at the stop site, or peptides detected only within the SCR extension.

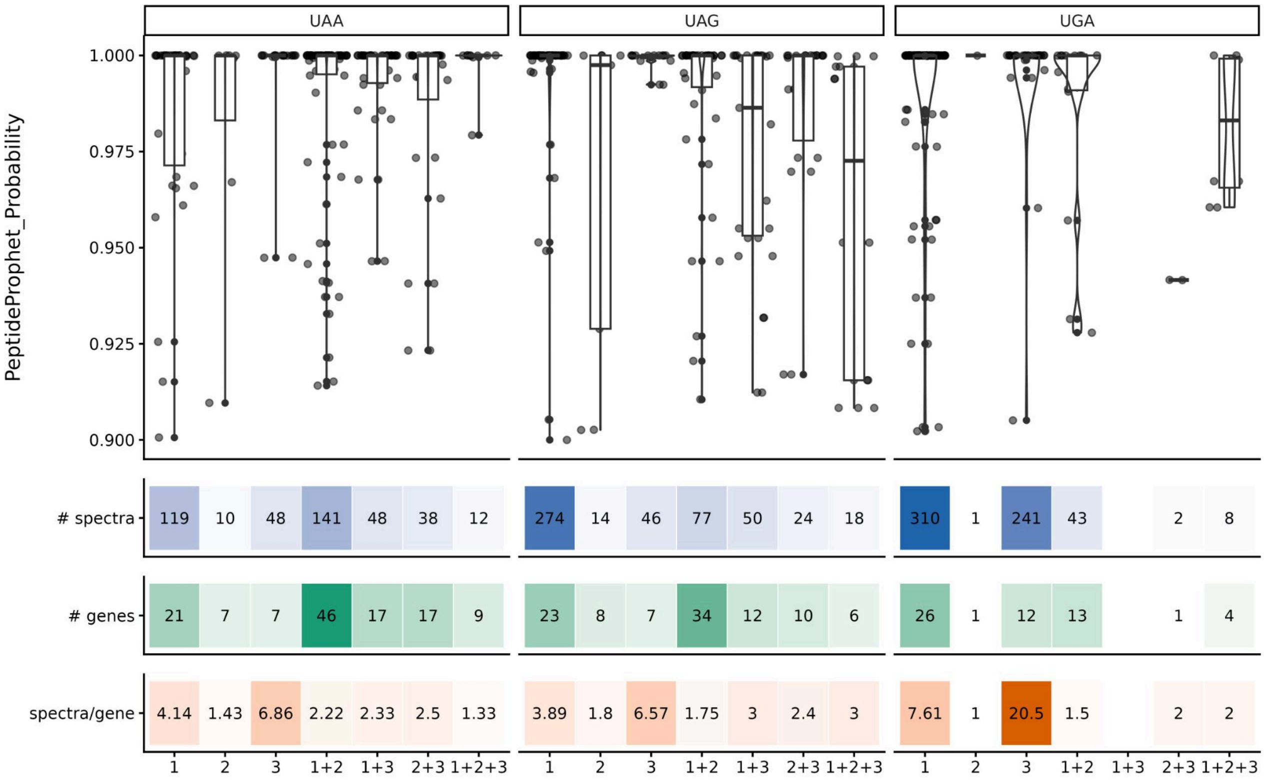


**Supplementary Figure S33.** Analysis of codon–anticodon mismatch configurations associated with amino acid incorporation during SCR. Top panels show the distribution of peptide identification confidence scores (PeptideProphet probability) for substitutions classified according to the inferred mismatch position between the stop codon and the decoding tRNA anticodon. Bottom heatmaps summarize the number of supporting spectra, independent genes, and normalized spectra-per-gene ratios for each mismatch category.

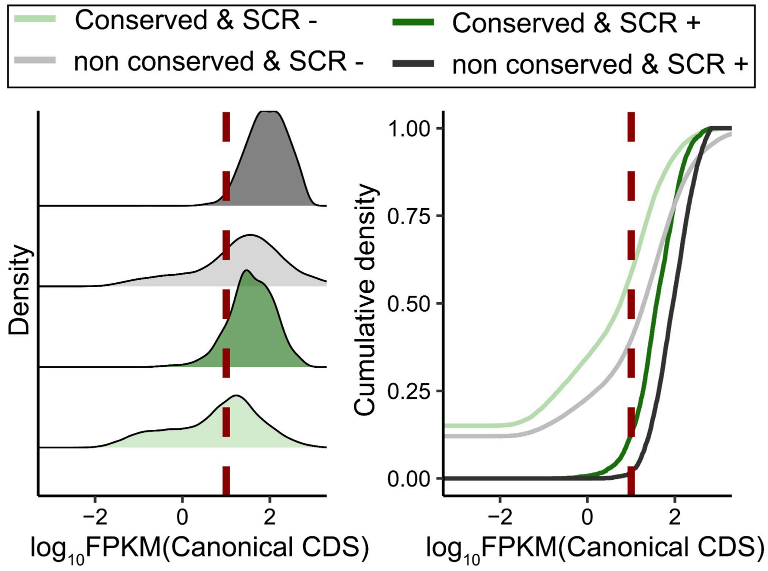


**Supplementary Figure S34.** Defining expression threshold for detectability of SCR.

The figure plots the distribution of expression values (Ribo-Seq FPKM of canonical CDS) in tier 1-2 genes, split by category as follows. Sample-specific SCR was predicted according to numerous metrics capturing Ribo-Seq density and robustness on the SCR extension region (Methods). Genes were categorized based on whether sample-specific SCR was predicted in at least two samples ("SCR +") and on whether they were detected as conserved SCR genes in Jungreis *et al* [23]("Conserved"). The red dashed line indicates the selected expression threshold for reliable SCR detection (FPKM > 10): ~90% of *bona fide* conserved SCR genes where sample-specific SCR was detectable have expression greater than this threshold; in contrast, only 40% of conserved SCR genes with no sample-specific SCR pass the threshold.

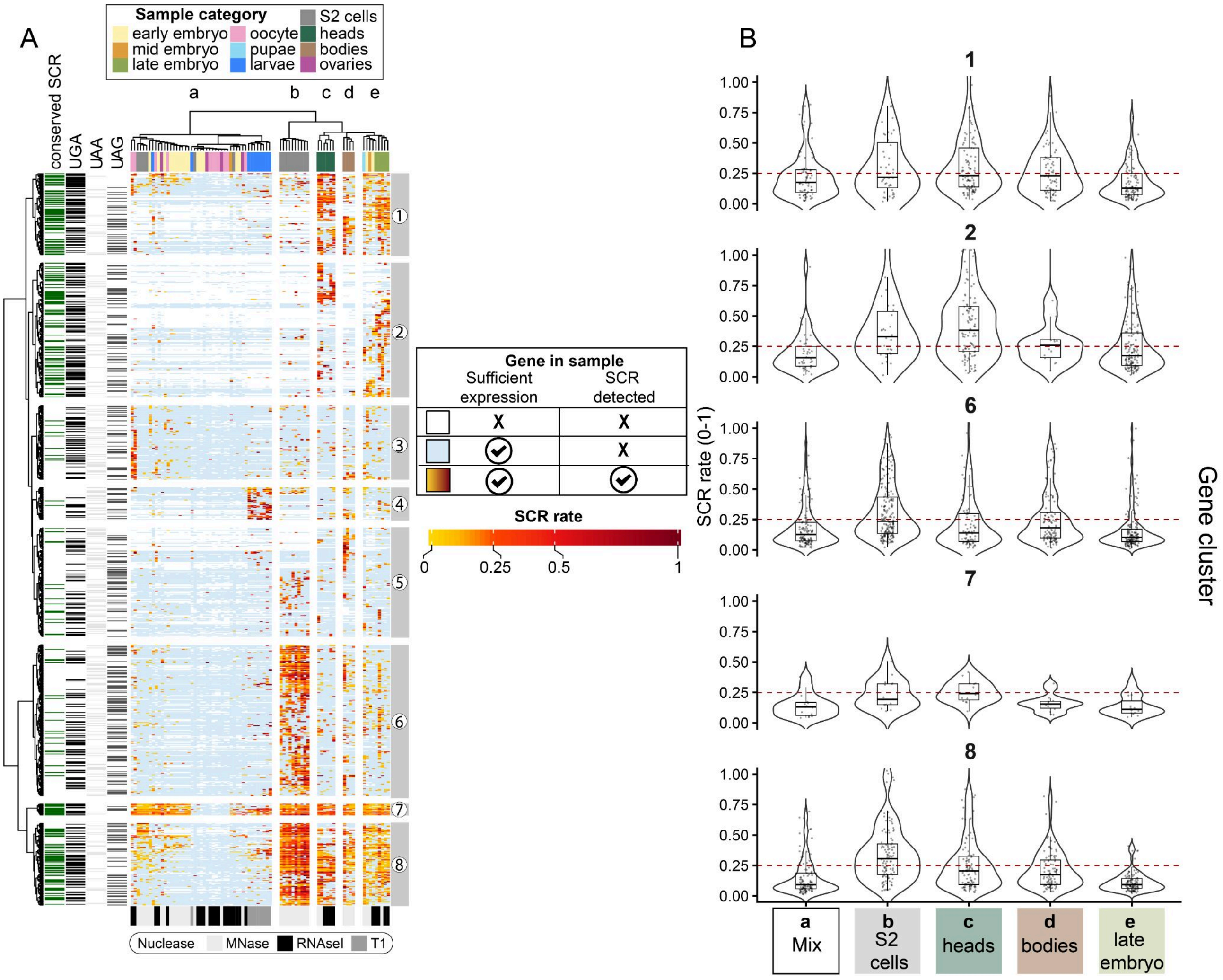


**Supplementary Figure S35.** Sample specific SCR rate estimation.

**A.** Clustered heatmap summarizing sample-specific SCR detection and SCR rate. Rows correspond to genes and columns to Ribo-Seq samples. Sample type is indicated by colored annotation bars at the top, and nuclease treatment is shown at the bottom. Heatmap colors represent both expression and SCR status: lowly expressed genes, considered insufficient for reliable SCR detection, are shown in white; expressed genes without detected SCR are shown in light blue; and genes with detected SCR are colored according to their estimated SCR rate, from low (yellow) to high (red) (black in Figure 5). Hierarchical clustering was performed using a simplified three-state matrix (not expressed, expressed without detected SCR, expressed with detected SCR), while the heatmap colors for SCR-positive genes reflect the estimated SCR rate. SCR rate was estimated from ribosome profiling data without noise correction as the ratio of ribosome density in SCR extensions (excluding the first 6 nt) to ribosome density in the canonical CDS (excluding the first 18 nt and the last 15 nt). Gene rows were annotated with stop codon identity and SCR predictions based on comparative genomics analysis from [23]

B. Distribution of SCR rates in samples with detected SCR across different sample groups. Sample groups were defined by splitting the column dendrogram into five categories (letters shown above the heatmap and in the x-axis): (a) a heterogeneous group of samples characterized by a low number of genes with detected SCR; (b) S2 cells; (c) heads; (d) bodies; and (e) a mixed group composed predominantly of late embryo samples. Five gene clusters (numbers shown on the left side of the heatmap and vertical facets) displaying sample-specific SCR patterns were selected to examine differences in SCR rates between sample groups (x axis). For each gene, SCR rates were calculated only in samples where SCR was detected (yellow-red cells in the heatmap in panel A). To aggregate measurements for each gene within each sample group, SCR rates were first log2-transformed, averaged across samples belonging to the same group, then back-transformed to the linear scale for visualization. The red dashed line indicates a SCR rate of 25%.

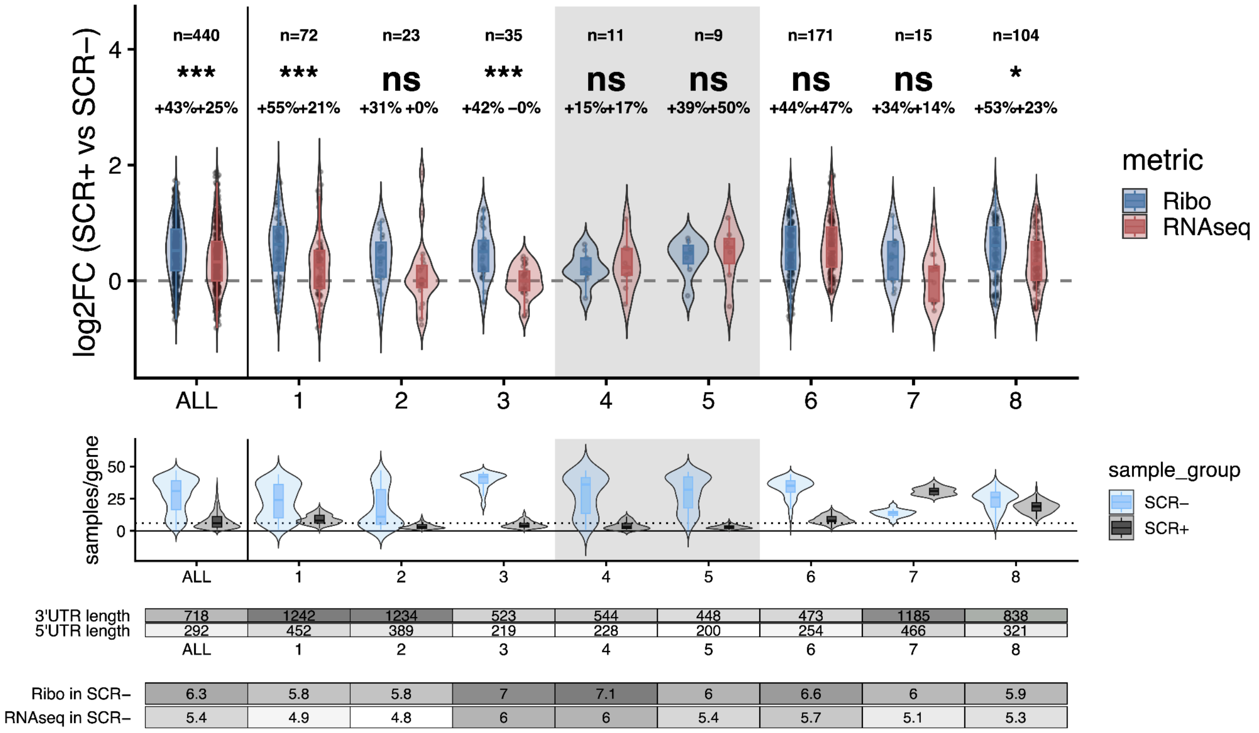


**Supplementary Figure S36.** Expression changes associated with SCR. Top: violin/boxplots showing gene-wise log2FC between the averages in SCR-positive (SCR+) and SCR-negative (SCR-) samples in Ribo-Seq (blue) and RNA-seq (red) expression, grouped by gene cluster (Methods). Percentages indicate median expression increases in SCR+ samples. Paired Wilcoxon tests adjusted by Benjamini-Hochberg were used to assess the difference between Ribo and RNA log2FC values: ns, not significant; * FDR < 0.05; *** FDR < 0.001. Mid: number of samples per gene available in this analysis, grouped by cluster. Only genes with at least 6 SCR+ and 6 SCR- samples were employed. Clusters with less than 12 genes are marked in grey and are omitted from Figure 5. Bottom: annotation panel showing median basal expression (SCR- samples, log2(FPKM)) and 5′UTR/3′UTR lengths per cluster.

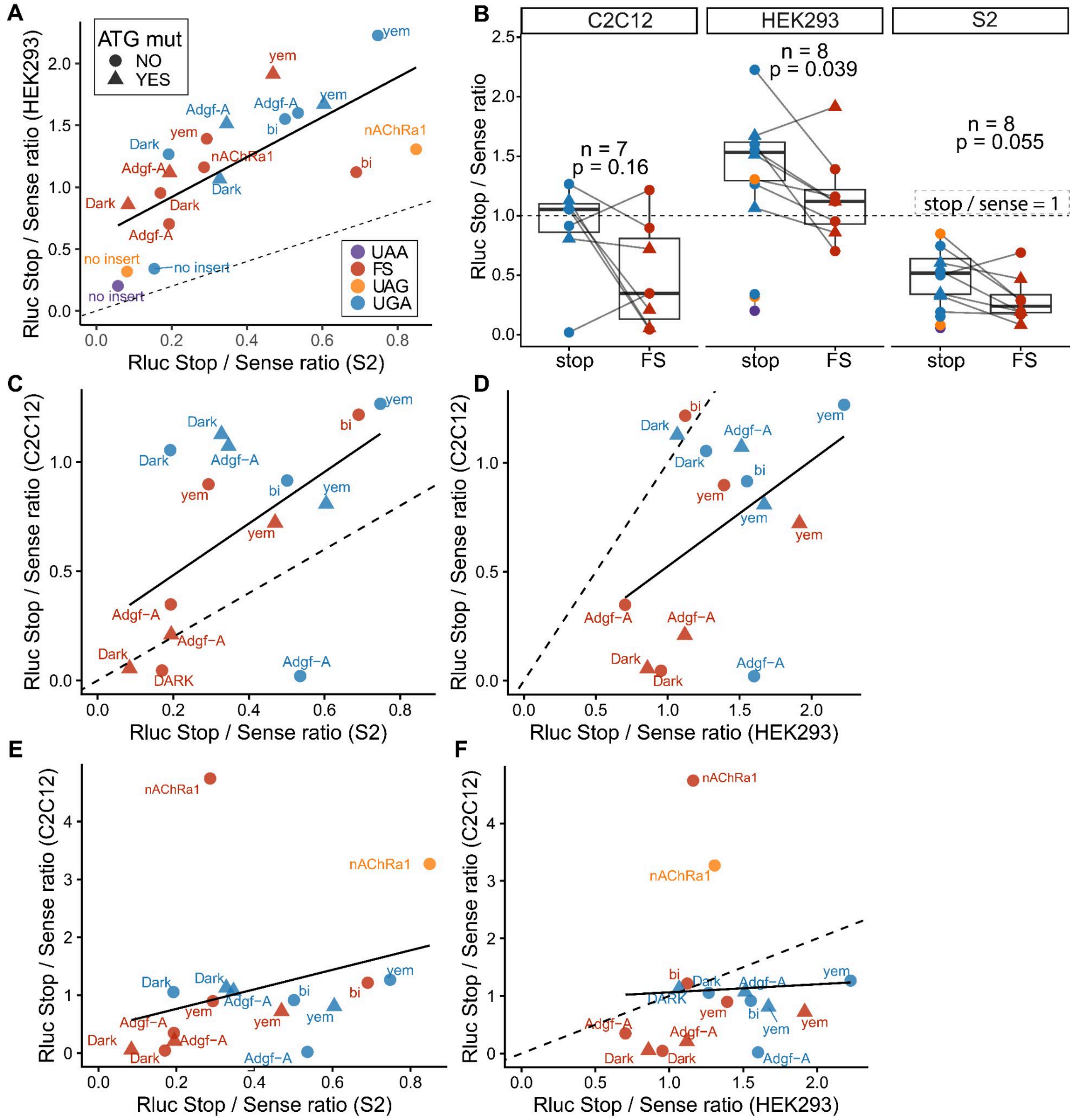


**Supplementary Figure S37.** SCR affects the expression of the reporter upstream of stop codon.

**(A).** Correlation of SCR-mediated enhancement of reporter signal in S2 cells and HEK293 cells. For each construct, Renilla luciferase (Rluc) signal (i.e. upstream of the stop codon) was normalized to the corresponding sense control. Constructs containing downstream ATGs were also tested after ATG mutation (triangles), while circles represent constructs without mutation (or lacking downstream ATGs). Pearson correlation coefficient (R = 0.72; p = 0.0004).

**(B).** Comparison of SCR constructs and their paired frameshift (FS) controls. P-values were calculated using a paired Wilcoxon test and are indicated in the plot.

**(C).** Correlation of SCR-mediated enhancement of reporter signal in S2 cells and C2C12 cells. See Panel A.Pearson correlation coefficient (R = 0.53; p = 0.048). The *nAChRa1* element appeared as outlier in C2C12 and was excluded here; the plot including it is available in panel F.

**(D).** Correlation of SCR-mediated enhancement of reporter signal in HEK293 cells and C2C12 cells. See panel A. Pearson correlation coefficient (R = 0.44; p = 0.1111). The *nAChRa1* element appeared as outlier in C2C12 and was excluded here; the plot including it is available in panel F.

**(E),(F).** Analogous plot to (C) and (D) without removing 1 outlier in C2C12 cells - *nAChRa1* (was excluded from this analysis because it consistently exhibited abnormally low Renilla luciferase signal in sense version, suggesting a construct-specific artifact). Pearson correlation coefficient (R = 0.31; p = 0.238 for C2C12 vs. S2 and R = 0.404; p = 0.869 for C2C12 vs. HEK293)

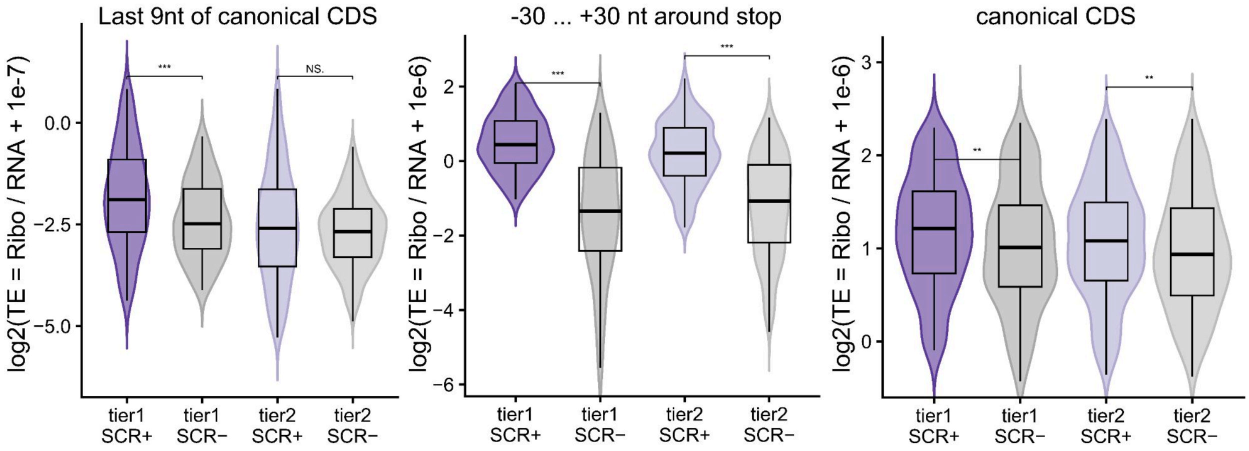


**Supplementary Figure S38.** Translation efficiency (TE) values for each gene in each sample.

Genes in which SCR was detected in a given sample (SCR+) are colored according to their tier classification, whereas genes that exhibited sufficient expression but did not show SCR are shown in grey (SCR-). For each gene, one value was calculated per category as the mean of log2-transformed TE values across samples. Left panel: TE calculated using read counts from the last 9 nt of canonical CDS. Middle panel: TE calculated using read counts from the region surrounding the stop codon. Right panel: TE calculated using read counts from the canonical CDS, trimmed by 18nt after the start and 15nt from stop to omit peaks associated to initiation and termination. Significance is indicated as NS, *, **, and *** ($p < 0.05$, 0.01, and 0.001, respectively; Wilcoxon test).

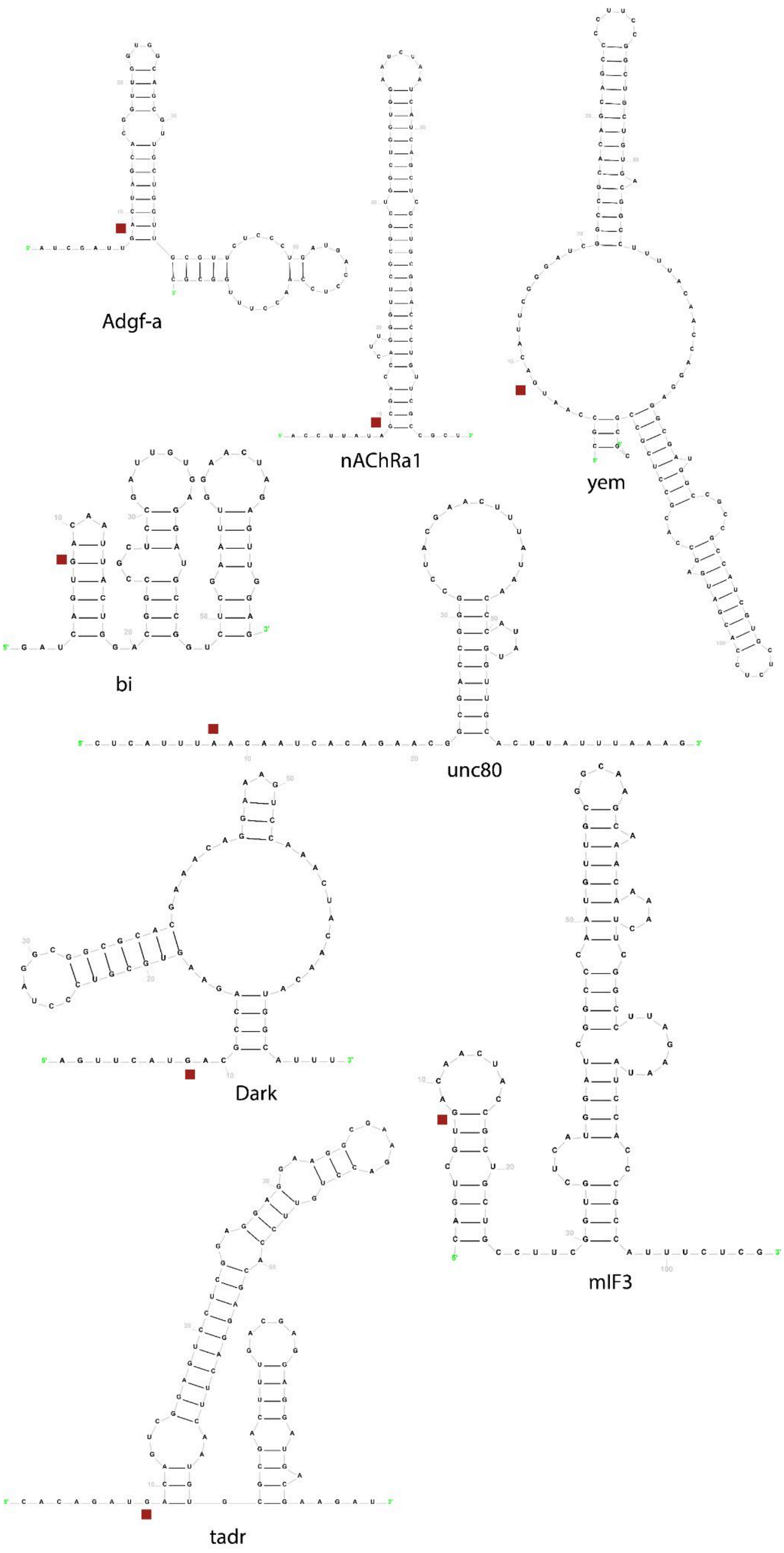


**Supplementary Figure S39.** RNAfold predictions of RNA secondary structures for genes identified in the screen. Assuming that RNA structures immediately upstream of the stop codon are likely to be melted by helicase activity, structure prediction was performed using sequences extending from 6 nt upstream of the stop codon onward. Stop codons are highlighted by red squares.

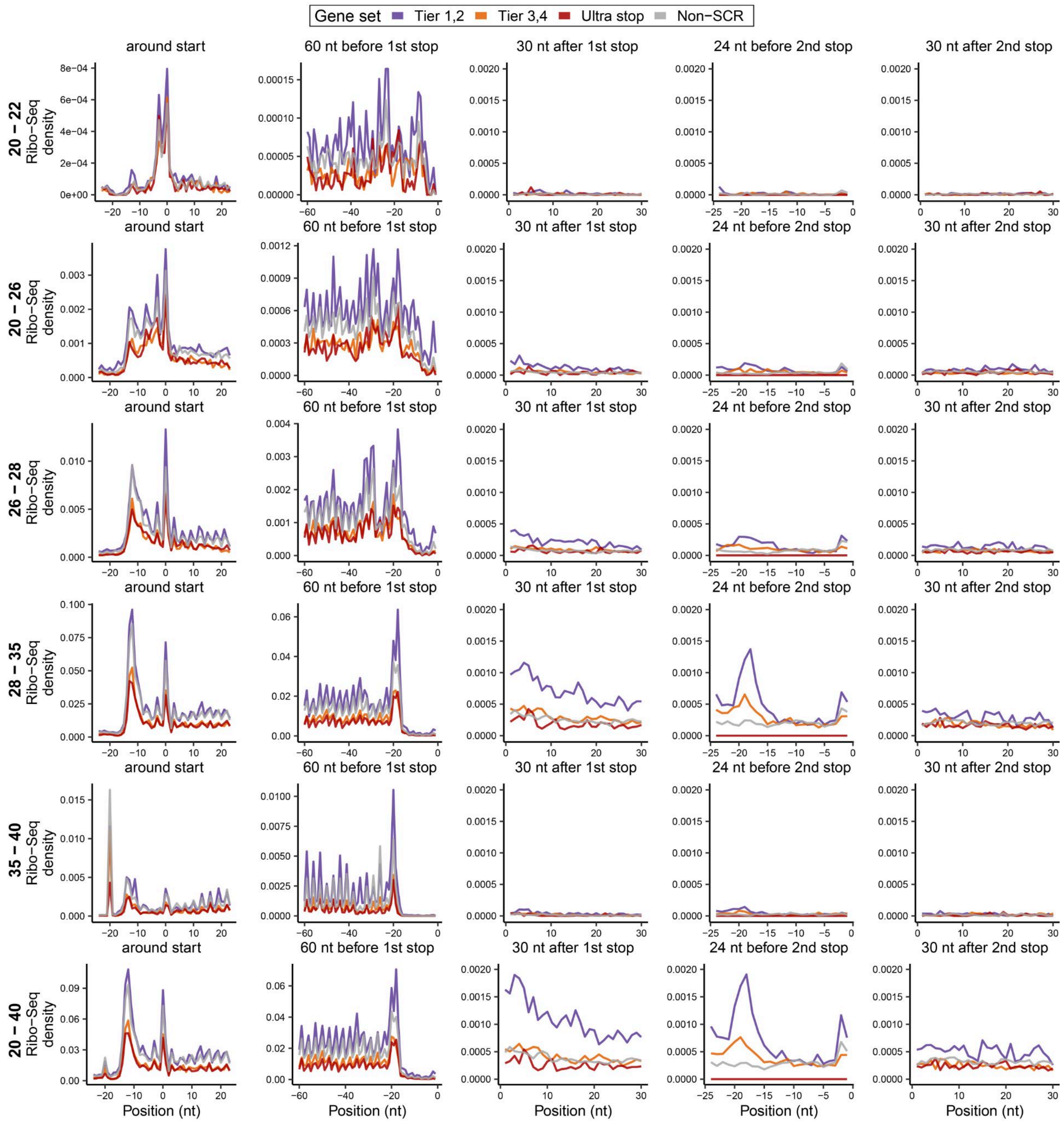


**Supplementary Figure S40.** Aggregated Ribo-Seq profiles across different read length ranges.

Genes were grouped into tiers based on ribosome profiling derived scores, with ultrastop and non-SCR genes included as controls. Reads from BAM files were filtered by length (y-axis facets). For each gene, mean read density at each nucleotide position was first computed by averaging across all BAM files. Aggregated Ribo-Seq profiles were then generated by averaging these per-gene profiles across genes within each group over defined windows around start and stop codons (x-axis facets). Read length ranges analyzed were 20-22 nt (associated with specific ribosome conformations), 20-26 nt, 26-28 nt, 28-35 nt, 35-40 nt, and 20-40 nt. Reads were aligned by their 5′ ends, and no P-site offset correction was applied.

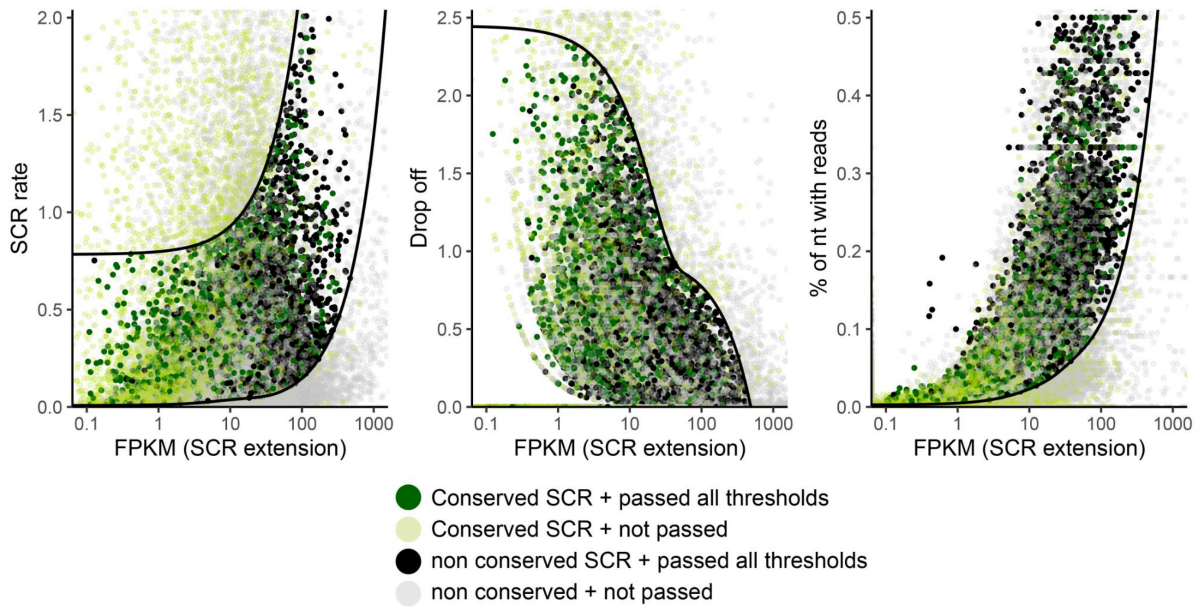


**Supplementary Figure S41.** Expression-dependent thresholds used for SCR detection.

Each point represents one gene in one sample, plotted against the FPKM of the SCR extension. Black curves indicate the expression-dependent thresholds learned from conserved SCR genes-sample values (light green and dark green dots together were used) using GAMs. A gene is classified as SCR-positive in a given sample only if it simultaneously satisfies all filtering criteria: RT rate within the learned lower and upper bounds, sufficient nucleotide coverage across the extension, acceptable drop-off after the second stop codon, expression ≥ 0.1, and coverage spanning at least the interquartile region of the extension. Point colors indicate whether a gene is conserved or non-conserved and whether it passes the full SCR criteria in that sample. Thus, even genes with values within the threshold for one parameter may still be classified as SCR-negative if they fail any of the others.

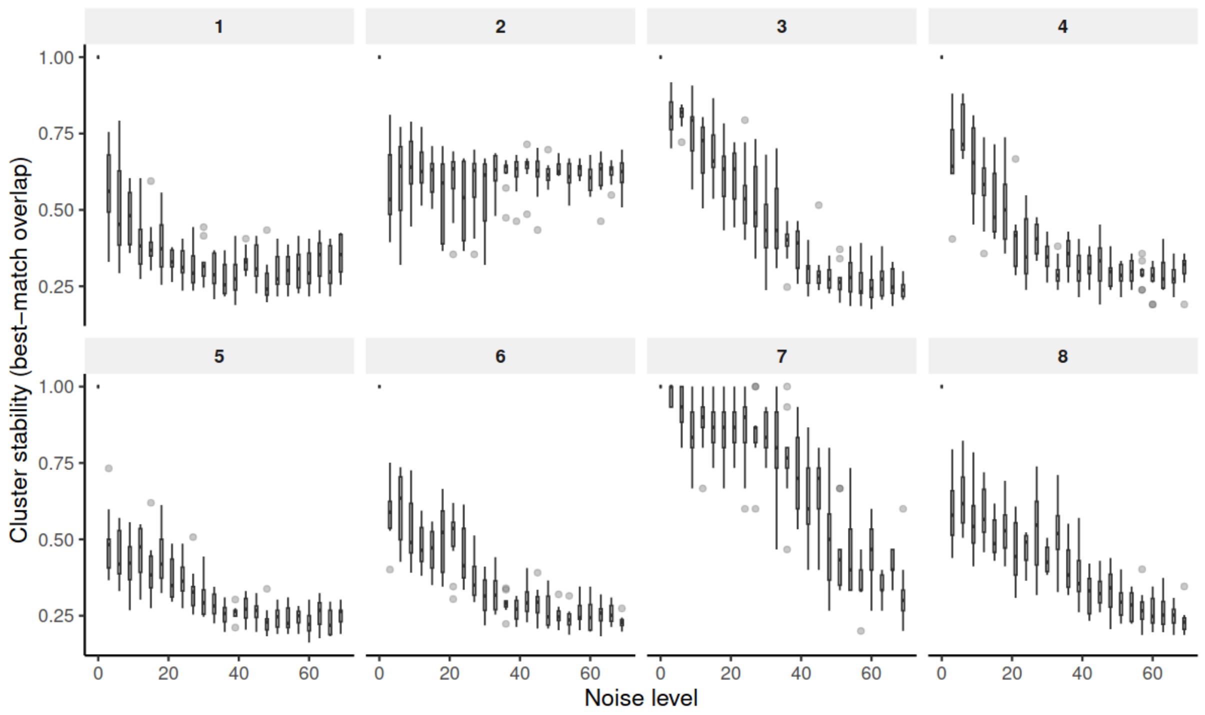


**Supplementary Figure S42.** Stability of clusters after noise addition.

Cluster stability is shown as the fraction of genes that remain assigned to the same cluster as in the original dendrogram after introducing noise (see Methods). Facets correspond to cluster numbers. The x-axis indicates the percentage of matrix values randomly swapped with fixed values representing expression states: 1 (not expressed; white in the heatmap), 2 (expressed but no SCR detected, lightblue in the heatmap), and 10 (SCR detected, black in the heatmap).

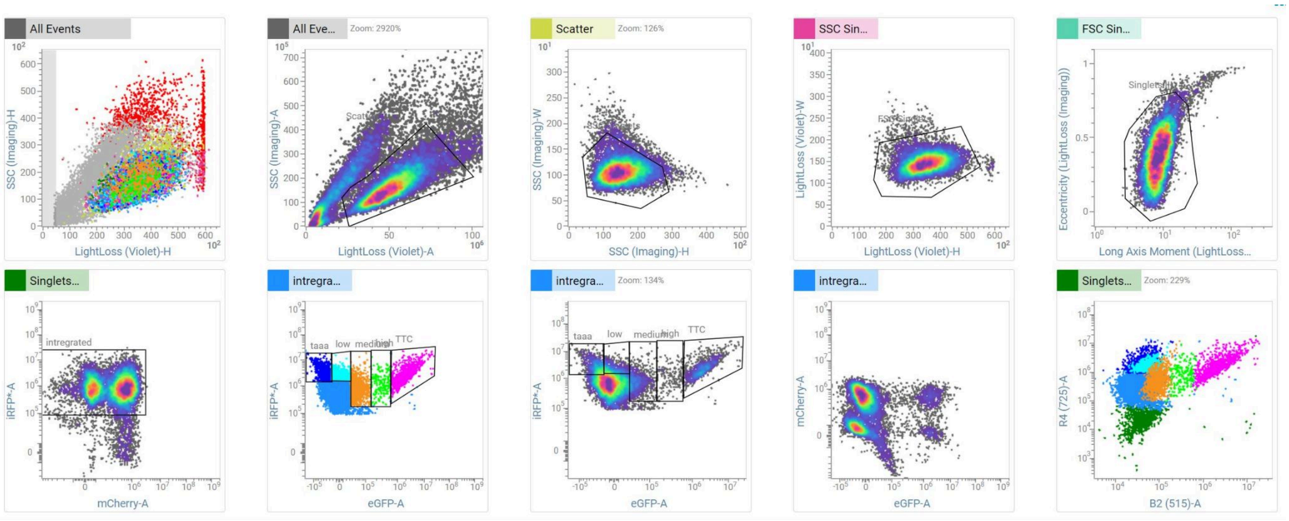


**Supplementary Figure S43.** Full gating strategy applied on the S8 sorter to identify live, singlet, iRFP-positive cells and sort them into four fractions based on GFP intensity.
